# APOE4 promotes cerebrovascular fibrosis and amyloid deposition via a pericyte-to-myofibroblast transition

**DOI:** 10.1101/2025.09.04.674192

**Authors:** Braxton R. Schuldt, Dominic Haworth-Staines, Andrea Perez-Arevalo, Diede W.M. Broekaart, Ashley Harlock, Leon Wang, Anna Bright, Georgia Gallagher, Grace Rabinowitz, Alison M. Goate, Towfique Raj, Ana C. Pereira, Joel W. Blanchard

**Affiliations:** Nash Family Department of Neuroscience and Friedman Brain Institute, Icahn School of Medicine at Mount Sinai, New York, NY, 10029, USA; Ronald M. Loeb Center for Alzheimer’s Disease, Icahn School of Medicine at Mount Sinai, New York, NY, 10029, USA; Institute for Regenerative Medicine, Black Family Stem Cell Institute, Icahn School of Medicine at Mount Sinai, New York, NY, 10029, USA; Department of Neurology, Icahn School of Medicine at Mount Sinai, New York, NY, 10029, USA

**Keywords:** APOE4, cerebrovascular degeneration, pericyte, myofibroblast, TGF-β, fibrosis, fibronectin, cerebral amyloid angiopathy, Alzheimer’s disease, iPSC modeling

## Abstract

Cerebrovascular disease is a major but poorly understood feature of Alzheimer’s disease (AD). The strongest genetic AD risk factor, APOE4, is associated with cerebrovascular degeneration, including vascular amyloid deposition and fibrosis. To uncover how APOE4 promotes cerebrovascular pathology, we assembled a single-cell transcriptomic atlas of human brain vasculature. In APOE4 carriers, pericyte abundance was significantly reduced and accompanied by the emergence of a myofibroblast-like cell population co-expressing contraction and extracellular matrix genes. Immunostaining confirmed non-vascular myofibroblasts in APOE4 human and mouse brains. We show that APOE4 pericytes transition into myofibroblasts that secrete fibronectin, which promotes vascular amyloid accumulation. Computational and experimental analyses identified elevated TGF-β signaling as the driver of this pericyte-to-myofibroblast transition. Inhibition of TGF-β restored pericyte coverage and reduced vascular fibrosis and amyloid to APOE3 levels, revealing a targetable mechanism linking APOE4 to cerebrovascular pathology in AD.

**Summary:** Cerebrovascular disease is a prominent but poorly understood component of Alzheimer’s disease (AD). The strongest genetic risk factor for AD, APOE4, is associated with multiple cerebrovascular pathologies, including vascular amyloid accumulation and fibrosis. APOE4 also renders the cerebrovasculature fragile to the point where the only disease-modifying AD therapeutic, anti-amyloid monoclonal antibodies, are warned against use in APOE4 carriers due to a high risk of hemorrhage and edema. To determine how APOE4 impacts cerebrovascular cells and pathogenesis, we assembled a high-resolution single-cell transcriptomics atlas of human cerebrovasculature from APOE4 carriers and non-carriers. In APOE4 individuals, we found that the number of microvascular pericytes was significantly reduced. Notably, loss of pericytes in APOE4 carriers was accompanied by the emergence of a unique population of mural cells characterized by high expression of contraction and extracellular matrix (ECM) genes, hallmarks of myofibroblasts. Immunostaining revealed the presence of myofibroblasts surrounding the microvasculature in APOE4 human hippocampi that were absent in age-matched APOE3/3 controls, even in cases of AD. Myofibroblast presence coincided with a significant increase in fibronectin and amyloid surrounding the vasculature. Myofibroblasts were also present around the microvasculature of aged APOE4/4 but not APOE3/3 mice, suggesting myofibroblast appearance is a direct effect of the APOE4 genotype and not a technical artifact of processing human tissue. To determine the mechanisms and functional implications of APOE4-mediated myofibroblast, we leveraged a vascularized human brain tissue (miBrain) derived from induced pluripotent stem cells. Similar to the post-mortem human brain, APOE4/4 miBrains showed significantly reduced microvascular pericyte coverage, coinciding with the emergence of myofibroblasts co-expressing ECM and contractile genes. Lineage tracing and genetic mixing experiments confirmed that the myofibroblasts emerge from APOE4 pericytes and secrete fibronectin-rich ECM. Knocking down fibronectin in APOE4 mural cells significantly reduced vascular amyloid accumulation. To determine how myofibroblast-like cells arise in the APOE4 brain, we performed computational analysis (NicheNet) on our post-mortem human single-cell transcriptomics cerebrovascular atlas. This predicted that TGF-β is the top causal driver of the pericyte-to-myofibroblast transition. Consistent with this prediction, we found that TGF-β ligands in astrocytes and receptors in mural cells are significantly upregulated compared to APOE3 controls. Chemical inhibition of TGF-β signaling in APOE4 miBrains and APOE4-TR mice significantly reduced myofibroblast presence, while concurrently increasing pericyte microvascular coverage and ultimately lowering vascular fibrosis and amyloid accumulation to APOE3 levels. Using a comprehensive three-pronged approach incorporating analysis of human post-mortem brain, APOE humanized mice, and human iPSC-derived vascularized brain tissue, we demonstrate that APOE4 causes increased TGF-β signaling, promoting a pericyte-to-myofibroblast transition that leads to vascular fibrosis and amyloid deposition. This provides critical insight into the mechanisms underlying APOE4 cerebrovascular dysfunction, highlighting new diagnostic and therapeutic strategies for a major AD risk population.

## Introduction

Alzheimer’s disease (AD) is a common and devastating neurodegenerative disorder characterized by progressive cognitive decline and brain atrophy. The vast majority of cases are classified as late-onset AD, where aging and genetic factors play a major role in disease risk^1^. Among these, the APOE4 allele stands out as the strongest genetic risk factor for AD identified through genome-wide association studies (GWAS)^2^. Remarkably, APOE4 differs from the more common APOE3 allele by a single amino acid (Cys112 ➔ Arg112), yet individuals carrying two copies of APOE4 face up to 16-fold increased risk of developing AD^3^. Furthermore, 40-65% of all individuals with AD carry at least one APOE4 copy, underscoring its relevance to disease pathogenesis^4^. Understanding how APOE4 contributes to AD progression is, therefore, critical to developing targeted therapies for this large and vulnerable patient population.

Since its identification as an AD risk variant, APOE4 has been implicated in dysfunction across nearly all major brain cell types. In neurons, APOE4 drives tau hyperphosphorylation and death^5–8^, in microglia, it promotes lipid accumulation, pro-inflammatory signatures, and impaired motility^9–14^, in astrocytes, it drives pro-inflammatory signalling^15–18^, and in oligodendrocytes, it disrupts myelination via cholesterol dysregulation^19^. Despite these widespread effects, APOE4 may exert its most profound influence on the cerebrovasculature^20–24^. Perivascular amyloid accumulation, blood-brain-barrier (BBB) breakdown, and microvessel degeneration are among the earliest changes that occur during AD pathogenesis and directly contribute to disease progression^25–27^. Furthermore, these vascular defects directly impair brain homeostasis and accelerate cognitive decline^28–36^. Even in the absence of AD pathology, APOE4 carriers show increased cerebrovascular dysfunction, suggesting that the vascular effects of APOE4 may be an upstream driver of AD pathogenesis^28,37–40^. While the advent of anti-amyloid monoclonal antibodies marked a major therapeutic milestone, their use in APOE4 carriers is limited by the elevated risk of amyloid-related imaging abnormalities (ARIA), including cerebral hemorrhage and edema^41,42^. The vulnerability of APOE4 carriers to the only FDA-approved AD therapeutic underscores a critical need: preventing or reversing APOE4-associated cerebrovascular damage may be essential to slowing or halting AD progression in this high-risk population.

Cerebral blood vessels are composed of endothelial cells, mural cells, and a basement membrane, with additional structural and functional support from astrocytic endfeet. Each of these components has been independently implicated in APOE4-mediated cerebrovascular degeneration. In endothelial cells, APOE4 promotes inflammatory and prothrombotic signaling while impairing tight junctions and compromising BBB integrity^43,44^. Pericytes are key contributors to cerebral amyloid angiopathy (CAA) and BBB breakdown, with APOE4 expression in pericytes directly driving these pathologies^28,33,45–50^. Beyond these canonical vascular cell types, single-cell studies have revealed an unexpected diversity of fibroblasts within brain barrier structures, suggesting emerging roles in cerebrovascular homeostasis, even in AD^51–56^. Supporting this, recent studies have identified protective genetic variants in vascular basement membrane components, such as fibronectin, that reduce AD risk in APOE4 carriers^57,58^. Moreover, the profibrotic cytokine TGF-β has been linked to vascular degeneration, further implicating dysregulation of the extracellular matrix in APOE4-associated cerebrovascular pathology^59–61^. Despite these findings across cell types and compartments of the APOE4 cerebrovasculature, the precise mechanisms through which APOE4 expression in vascular cells leads to cerebrovascular dysfunction and AD pathogenesis remain poorly defined.

To address this gap, we assembled a high-resolution transcriptomics atlas of the human cerebrovasculature by integrating previously published single-nuclei RNA-sequencing (snRNA-seq) datasets from APOE4 carriers and non-carriers. We complement this computational approach with postmortem immunostainings, isogenic iPSC models of the human brain, and humanized APOE knock-in mice. Through this comprehensive three-pronged approach (human postmortem, human isogenic iPSC models, and APOE-TR mice), we discover that APOE4 promotes a pericyte-to-myofibroblast transition in the brain. We establish that elevated TGF-β signaling in the APOE4 brain drives this transition, resulting in a loss of pericyte coverage and excessive fibronectin deposition along the vasculature. We show that fibronectin secretion from APOE4 myofibroblasts directly promotes perivascular amyloid accumulation and correlates with an earlier age of AD diagnosis and more severe cerebral pathology. Importantly, we demonstrate that TGF-β inhibition in APOE4 iPSC-derived human brain tissue and APOE4 mice reverses the pericyte-to-myofibroblast transition and reduces cerebrovascular fibrosis and amyloid accumulation, thus providing a direct mechanistic link between APOE4, TGF-β, vascular fibrosis, and AD pathogenesis.

## Results

### Harmonized single-nucleus atlas of the human APOE4 cerebrovasculature

To assemble a single-nuclei transcriptomics atlas of human cerebrovasculature from APOE3/3 and APOE4 carriers, we integrated three previously published snRNA-seq datasets of the postmortem human brain^11,62,63^ (**Figure 1A**). These studies were selected for their inclusion of *APOE* genotype information and complementary strategies to enrich vascular cell populations that are typically underrepresented in single-cell transcriptomic datasets. Cross-sample or cross-batch integration using Harmony was performed within each dataset, followed by vascular cell annotation. To minimize confounding variables, we then performed unbiased propensity score matching independently within each dataset to select a final cohort of APOE4 carriers and non-carriers matched for age, sex, AD status, and the number of vascular cell nuclei contributed per individual (**Extended Data 1A-C**). Vascular cells from these matched individuals were harmonized to generate an integrated cerebrovascular atlas (**Figure 1B; Extended Data 1D, 1E**), capturing diverse vascular subtypes across individuals and providing a comprehensive foundation for downstream analyses. Integration was performed using Harmony across the variables, subject, and dataset. Successful integration was confirmed by pseudobulk correlation analysis and variance partitioning, which demonstrated reduced dataset-driven effects while preserving biological variability (**Extended Data 1E-G**). After processing and quality control, the atlas consisted of ∼66,000 nuclei from 220 individuals (32,509 APOE3/3 (n = 110; F=57, M=53) and 33,179 APOE4 (n = 110; F=53, M=57; APOE3/4 = 94, APOE4/4 = 16)) (**Extended Data 1A, 1B**). The captured brain regions from the original studies included the hippocampus (Yang and Sun datasets), prefrontal cortex (Haney and Sun datasets), and angular gyrus, entorhinal cortex, midtemporal cortex, and anterior thalamus (Sun dataset). Clustering based on canonical cell-type-specific marker gene expression identified distinct populations of the major cerebrovascular cell types: astrocytes (*AQP4*), endothelial cells (*PECAM1*), pericytes (*PDGFRB*), and smooth muscle cells (SMCs) (*MYH11*) **(Figure 1B, Extended Data 1D, Supplementary Table S1)**.

**Figure 1.**
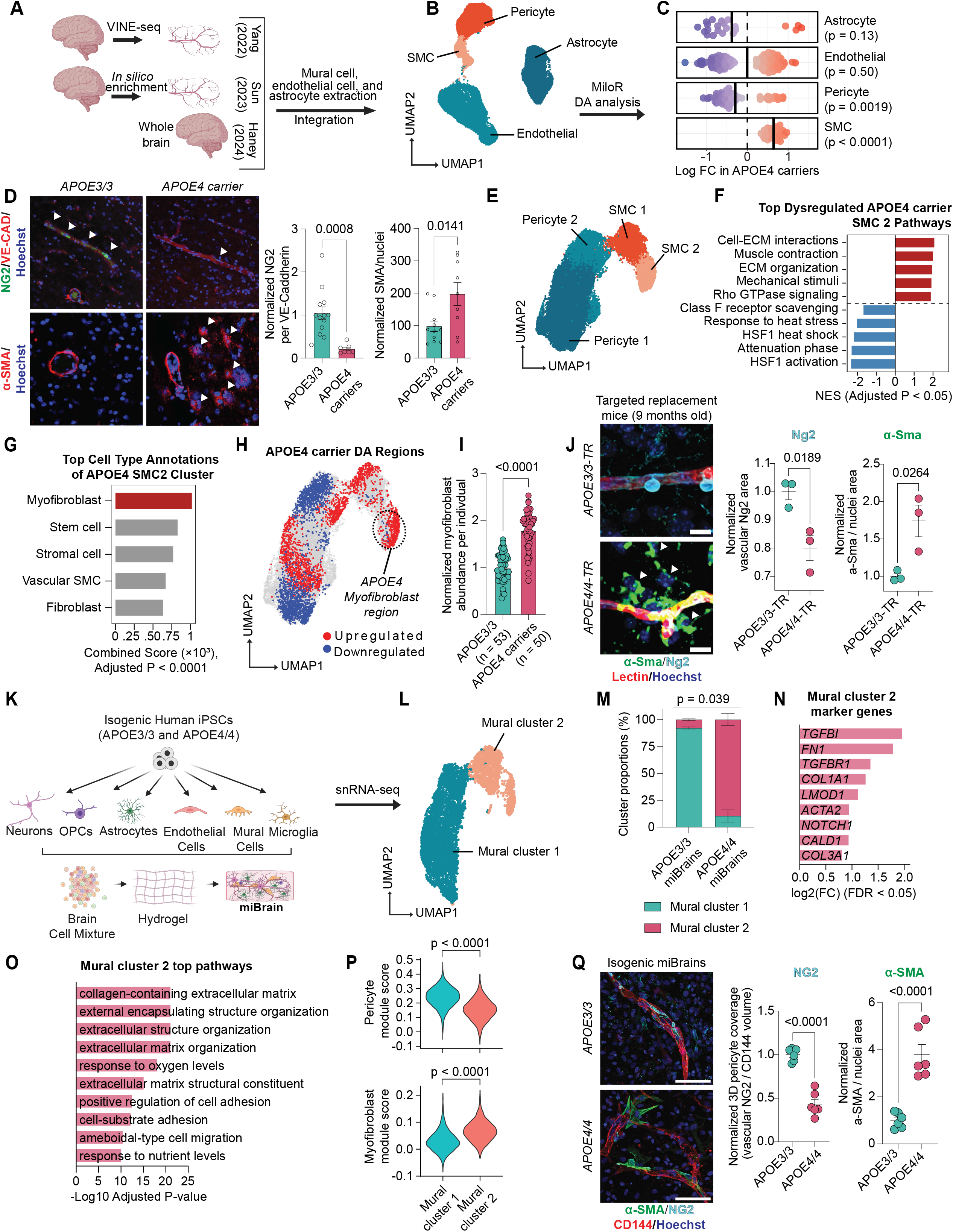
A myofibroblast-like mural cell population is enriched in the APOE4 brain. **A.** Schematic of cerebrovascular atlas generation from single nuclei RNA-sequencing of the post-mortem human brain in APOE4 carriers and non-carriers. **B.** UMAP of harmonized blood-brain barrier cell types. **C.** Significant differentially abundant (DA) cell neighborhoods per cell type in APOE4 carriers vs. non-carriers using the miloR package (spatial FDR < 0.05, positive logFC indicates enrichment in APOE4 carriers). Data points are individual cell neighborhoods. Solid vertical lines represent mean logFC for each cell type. A one-sample Wilcoxon signed-rank test with the Benjamini-Hochberg correction was performed on the distribution of logFC values for each cell type to test whether the median logFC significantly differed from 0. **D.** Representative images of post-mortem human hippocampus stained for markers of pericytes (NG2), smooth muscle cells (αSMA) and blood vessels (VE-CAD) in APOE3/3 and APOE4 carriers. Arrows indicate vascular pericytes (top) or non-vascular αSMA+ cells (bottom). Pericyte coverage for each genotype was calculated by quantifying the overlapping area of pericyte and blood vessel staining, expressed relative to APOE3/3. The area positive for αSMA was quantified, normalized to nuclei, and expressed relative to APOE3/3. Data points represent individuals (NG2: N = 13 APOE3/3 and N = 7 APOE4 carriers; αSMA: N = 11 APOE3/3 and N = 9 APOE4 carriers). Bars are group means ± SEM. P-values were calculated using an unpaired two-tailed Student’s t-test. **E.** UMAP of mural cell types after subclustering. **F.** Differential gene expression between APOE4 carriers and non-carriers for each mural cell type was performed using the MAST package. Genes were then ranked according to sign(logFC) * -log10(P-value) and input into the fgsea package using the REACTOME database. The top significantly upregulated and downregulated pathways in APOE4 carrier SMC_2 cells compared to APOE3/3 are shown (FDR < 0.05, ranked by adjusted p-value). Redundant pathways were merged to enhance readability. **G.** Top cell type annotations of SMC_2 cells from APOE4 carriers. The differentially expressed genes (p < 0.05, logFC > 0.25) upregulated in APOE4 SMC_2 cells from the pseudobulk approach in Extended Data 2D were input into EnrichR with the CellMarker database. Redundant cell types were merged to enhance readability. **H.** Differential abundance analysis on the mural cell clusters was performed with DAseq. Red and blue indicate enrichment and depletion in APOE4 carriers, respectively. Given the overlap of the APOE4-enriched SMC_2 region with ECM and contraction co-signatures, and the analyses performed in Extended Data 2H-J, this region was annotated as myofibroblasts. **I.** Subject-level model-estimated probability of myofibroblasts derived from a covariate-adjusted binomial generalized linear model (logit link) described in Extended Data 2M. Data points represent individuals (N = 53 APOE3/3 and N = 50 APOE4 carriers). Bars indicate marginal means ± SEM estimated from the model. Values were normalized to APOE3/3. P-values represent the marginal APOE genotype effect derived from the binomial GLM. **J.** Representative images of small vessels (<6 μm) in the corpus callosum of age- (9 to 10 months of age) and sex-matched APOE3/3 and APOE4/4-TR mice stained for markers of blood vessels (lectin), myofibroblasts (αSma), and pericytes (Ng2). Scale bar, 10 µm. For visualization only, vessels were cropped and a median filter was applied to the αSma channel. Arrows indicate non-vascular αSma+ myofibroblasts. The area positive for αSma was quantified, normalized to nuclei, and expressed relative to APOE3/3. Pericyte coverage was measured by quantifying the overlapping Ng2 and lectin area, normalized to APOE3/3. Data points represent mean values from individual mice (n = 6 images per mouse, n = 3 mice per genotype). Bars are mean group values ± SEM. P-values were calculated using unpaired two-sample Student’s t-tests. **K.** Schematic of the miBrain model. **L.** UMAP of miBrain mural cells derived from single-nuclei RNA-sequencing. **M.** Mural cluster proportions per genotype. Bars are mean group values ± SEM (n = 2 miBrains per genotype). P-values were calculated using an unpaired two-sample Student’s t-test with Welch’s correction. **N.** Representative marker genes of mural cluster 2. The MAST package was utilized to identify differentially expressed genes (|logFC| > 0.75, adjusted p < 0.05). **O.** The top enriched GO pathways in mural cluster 2. The differentially expressed genes identified in N. were input into ClusterProfiler. **P.** Pericyte and myofibroblast module scores per mural cluster. Scores were calculated using Seurat’s *AddModuleScore()* function based on the Pericyte_1 and myofibroblast marker genes from the human cerebrovascular atlas. **Q.** Representative images of isogenic APOE3/3 and APOE4/4 miBrains stained for markers of blood vessels (CD144), myofibroblasts (αSMA), and pericytes (NG2). Scale bar, 75 µm. Three-dimensional pericyte coverage was measured by the overlapping NG2 and CD144 volume, normalized to the total CD144 volume, and expressed relative to APOE3/3. The area positive for αSMA was quantified, normalized to nuclei, and expressed relative to APOE3/3. Data points represent mean values from individual miBrains (n = 6 images per miBrain, n = 6 miBrains per genotype). Bars are mean group values ± SEM. P-values were calculated using unpaired two-sample Student’s t-tests.

### APOE4 alters vascular cell composition in the human brain

Previous transcriptomic studies of the APOE4 vasculature have primarily focused on differential gene expression, with limited statistical power to detect changes in cell-type abundance^64^. By harmonizing multiple datasets into a single, integrated atlas, we were able to overcome these limitations and robustly assess whether APOE4 alters the cerebrovascular cell-type composition. We performed differential abundance analysis across all vascular cell types using MiloR, a k-nearest-neighbor graph-based approach designed to detect changes in non-discrete cell populations along differentiation or developmental trajectories^65^. Given the plasticity of vascular cell states and the absence of consistent markers to clearly delineate vascular subtypes^66^, this approach is well-suited for capturing abundance differences across transitional populations. Using MiloR, we identified 250 cell neighborhoods with significant differential abundance between APOE4 carriers and APOE3/3 individuals (spatial FDR < 0.05) (**Supplementary Table S2**).

### Pericyte loss in APOE4 carriers coincides with the emergence of atypical non-vascular SMCs

To examine the directionality of APOE4-associated changes in vascular cell composition, we plotted the fold changes of significantly altered neighborhoods for each cell type **(Figure 1C)**. APOE4 carriers showed a significant reduction of pericytes (0.29-log-fold decrease, p = 0.0019) and a significant increase of SMCs (0.65-log-fold increase, p < 0.0001) compared to APOE3/3 individuals. In contrast, astrocytes and endothelial cells exhibited no significant net changes in abundance (0.38-log-fold decrease in astrocytes, p = 0.13; 0.002-log-fold decrease in endothelial cells, p = 0.50). We confirmed these results with Dirichlet regression, controlling for age, sex, AD status, and dataset of origin, leading us to focus subsequent analyses on mural cells **(Extended Data 1H)**.

Age-related cognitive decline is associated with hippocampal cerebrovascular degeneration and elevated cerebrospinal fluid (CSF) levels of the pericyte injury marker soluble PDGFRβ (sPDGFRβ)^34,67,68^. APOE4 further accelerates cognitive decline and increases CSF sPDGFRβ in humans^28^, suggesting that the reduced pericyte abundance we observed in APOE4 carriers may reflect pericyte injury or a cell-state transition. To directly assess APOE4-associated pericyte loss, we performed NG2 immunostaining on postmortem human hippocampal sections. APOE4 carriers exhibited a significant reduction in NG2-positive pericyte coverage along hippocampal vasculature compared to age-matched APOE3/3 individuals (0.21 ± 0.042-fold change, corresponding to a 79% decrease relative to APOE3/3, p = 0.0008, **Figure 1D**). We next asked whether APOE4 pericyte loss corresponded with the increase in SMCs identified through differential abundance analysis. Human postmortem staining revealed significantly increased αSMA immunoreactivity in the APOE4 hippocampus compared to APOE3/3 controls (1.97 ± 0.35-fold increase, p = 0.0141, **Figure 1D**). However, while αSMA positivity was primarily confined to large vessels in APOE3/3 hippocampi, APOE4 carriers displayed abundant αSMA immunoreactivity throughout the parenchyma **(Figure 1D)**. Together, these findings indicated that APOE4 carriers exhibit both pericyte loss and the emergence of atypical non-vascular αSMA-positive cells, consistent with a mural cell phenotypic shift. This prompted us to further investigate whether APOE4 alters mural cell identity at the transcriptional level.

### A myofibroblast-like mural cell population is enriched in the APOE4 brain

To investigate APOE4-associated transcriptional changes in mural cells, we subclustered the original pericyte and SMC populations. Consistent with previous studies^62,63^, we identified two pericyte subtypes (Pericyte_1 and Pericyte_1) and two SMC subtypes (SMC_1 and SMC_2) **(Figure 1E, Extended Data 2A, Supplementary Table S3, S4)**. We then performed differential gene expression analysis between APOE4 carriers and APOE3/3 homozygotes within each mural cell subtype and employed gene set enrichment analysis (GSEA) to identify dysregulated pathways **(Extended Data 2B, 2C, Supplementary Table S5, S6).** Among all mural cell subtypes, the Pericyte_1 subcluster had the most differentially expressed genes (1210 genes), followed by SMC_2 (260 genes), SMC_1 (184 genes), and Pericyte_2 (98 genes) (**Extended Data 2B**). However, for the GSEA results, SMC_2 showed the greatest number of dysregulated pathways with 61, followed by Pericyte_1 (9 pathways), SMC_1 (7 pathways), and Pericyte_2 (6 pathways) (FDR < 0.05; **Extended Data 2C, Supplementary Table S5, S6**).

Due to the large number of SMC_2 dysregulated pathways, we hypothesized that transcriptional changes in this cluster may underlie the non-vascular αSMA staining pattern observed in the APOE4 postmortem human brain. Pathway enrichment analysis revealed that the most upregulated pathways in APOE4 SMC_2 cells were related to cellular contraction (e.g., muscle contraction, response to mechanical stimuli, RHO GTPase signaling) and extracellular matrix (ECM) production (e.g., cell-ECM interactions, ECM organization) (**Figure 1F**). To account for intra-individual cellular correlations, we confirmed upregulation of contraction- and ECM-related pathways in APOE4 SMC_2 cells using a pseudobulk approach (FDR < 0.05) (**Extended Data 2D, Supplementary Table S7, S8**). SMCs are classically described as adopting either a ***contractile state*** (high expression of *ACTA2* and *TAGLN* and low ECM gene expression), or a ***synthetic state*** (low contractile gene expression and high levels of ECM components, including *FN1* and various *collagens*)^69–71^. To determine whether the APOE4 SMC_2 population reflects an expansion of these canonical states or a distinct phenotype with simultaneous activation of both programs, we calculated contraction and ECM gene signature scores for mural cells in our cerebrovascular atlas using established marker genes (**Extended Data 2E)**. Consistent with the pathway analysis, both contraction and ECM signatures were significantly upregulated in SMC_2 cells from APOE4 carriers (contraction score: 1.50 ± 0.14-fold increase, p = 0.0242; ECM score: 1.59 ± 0.10-fold increase, p = 0.0071**; Extended Data 2F**). Importantly, the proportion of SMC_2 cells co-expressing both ECM and contractile signatures was significantly higher in APOE4 carriers (1.44 ± 0.09-fold increase, p = 0.0269; **Extended Data 2F**).

The concurrent expression of both signatures is a hallmark of ***myofibroblasts***, the principal effector cells of tissue fibrosis^72–77^. Although myofibroblasts have been extensively studied in peripheral organs^78–81^, they have also been reported in the brain following traumatic injury or ischemic events^82,83^, where some are thought to arise from pericytes that detach from the vasculature^82^. To determine whether the APOE4-enriched SMC_2 population resembles myofibroblasts, we performed unbiased cell-type annotation using EnrichR with the CellMarker database^84–86^. Myofibroblasts were identified as the top-matching cell type for the APOE4 SMC_2 gene signature **(Figure 1G, Supplementary Table S9)**. Consistent with this annotation, myofibroblast gene activity scores were significantly higher in SMC_2 cells from APOE4 carriers compared with APOE3/3 individuals (1.56 ± 0.12-fold increase, p = 0.0112; **Extended Data 2G**).

To determine whether myofibroblasts are selectively enriched in APOE4 carriers, we performed differential abundance analysis of mural cell clusters using DAseq. This analysis revealed a distinct subregion within the SMC_2 cluster that was significantly enriched in APOE4 individuals and spatially colocalized with the peak of ECM and contractile gene co-expression **(Figure 1H).** Differential gene expression and pathway enrichment analyses further showed that this APOE4-enriched subregion upregulated genes involved in cell–substrate junctions, ECM organization, and contractile fibers (e.g., *FN1*, *COL3A1*, *ACTA2*, *COL8A1*, *TGFβ1I1*), hallmark features of a myofibroblast state^87^. In contrast, the remaining SMC_2 cells expressed genes linked to vascular stability (e.g., *FLT1*, *EPAS1*, *TIMP3*, *ADAMTS9*) and cell-cell junctions **(Extended Data 2H, I, Supplementary Table S10, S11)**. These non-myofibroblast-like SMC_2 cells also downregulated cell migration pathways, consistent with a more quiescent, vessel associated phenotype (**Supplementary Table S11**). Unbiased annotation of the APOE4-enriched subregion again identified myofibroblast as the top-matching cell type **(Extended Data 2J, Supplementary Table S12)**. The expansion of this SMC_2 population in APOE4 carriers was further validated with Dirichlet regression and miloR (**Extended Data 2K, L**). Together, the transcriptional profile of APOE4 SMC_2 cells, the results of unbiased cell-type annotation and differential abundance analyses, and the presence of non-vascular parenchymal αSMA staining in postmortem brain tissue strongly support that the APOE4-enriched SMC_2 subpopulation represents myofibroblasts.

Previous studies have shown that cerebrovascular degeneration in APOE4 carriers occurs independently of amyloid and tau pathology^28^, but the mechanisms by which APOE4 drives cerebrovascular degeneration remain unclear. Furthermore, sex differences have been reported in the prevalence and severity of AD vascular pathology, including cerebral amyloid angiopathy^88^. We therefore asked whether the myofibroblast population that we identified reflects primarily an APOE4-dependent vascular phenotype or instead arises secondarily from AD pathology or sex-specific effects. For each individual in our atlas, we quantified the proportion of myofibroblasts and modeled these values using a binomial generalized linear model with APOE genotype, AD status, age, sex, and dataset origin as covariates. APOE×AD status and APOE×sex interaction terms were additionally incorporated to test whether disease state or sex modified the effect of APOE4 (**Extended Data 2M**). Model diagnostics confirmed a good fit with no significant violations of assumptions. Across all individuals, APOE4 emerged as the strongest predictor of myofibroblast abundance (odds ratio = 3.05, 95% CI: 2.22-4.19, p < 0.0001, **Extended Data 2M**) and APOE4 carriers exhibited significantly higher predicted myofibroblast proportions than APOE3/3 individuals when averaged across all covariates (1.79 ± 0.06-fold increase, p < 0.0001; **Figure 1I**). Importantly, stratified model predictions demonstrated that this effect was consistent across both sexes and across AD and non-AD individuals. Within APOE4 carriers, neither AD diagnosis nor sex significantly influenced myofibroblast abundance (**Extended Data 2N**). Finally, because the integrated datasets differed substantially in vascular cell recovery, we performed a sensitivity analysis to determine whether the observed APOE4 effect on myofibroblast abundance was driven by any single cohort. We fit a binomial generalized linear model using a leave-one-dataset-out strategy, sequentially excluding a dataset and refitting the model. Across all exclusion combinations, APOE4 genotype remained a significant predictor of increased myofibroblast abundance (**Extended Data 2O**). Taken together, these findings indicate that the observed myofibroblast population emerges due to APOE4 itself rather than secondarily to neurodegeneration, sex-specific pathology, or dataset artifacts.

To further test whether APOE4 promotes myofibroblast formation in the absence of AD or other cerebrovascular pathology, we analyzed aged APOE targeted replacement (TR) mice in which the murine *Apoe* gene is replaced with human *APOE3/3* or *APOE4/4* alleles. Bulk RNA-seq data^89^ from cerebral cortices of 4-month-old APOE3/3, APOE3/4, and APOE4/4-TR mice were scored for myofibroblast gene activity using GSVA with marker genes from our human cerebrovascular atlas. APOE4 gene dosage was positively associated with myofibroblast gene activity (β = 0.072, p = 0.015; **Extended Data 3A**). Immunofluorescence of hippocampal and cortical vessels further validated these findings. While both genotypes exhibited strong αSMA expression in large cerebral vessels (**Extended Data 3B**), APOE4/4 mice displayed a significant reduction in pericyte coverage of small vessels (< 6 µm) compared with APOE3/3 controls (0.80 ± 0.044-fold change; 20% decrease; p = 0.0189; **Figure 1J**). This pericyte loss coincided with increased αSMA immunoreactivity in cells proximal to small vessels and within the parenchyma, consistent with a myofibroblast identity (1.74 ± 0.21-fold increase; p = 0.0264; **Figure 1J**). We further validated this phenotype using another contractile protein highly expressed in myofibroblasts, Tagln, once again finding an increase in small vessel and parenchymal immunoreactivity in aged APOE4/4 mice (1.76 ± 0.20-fold increase; p = 0.0464; **Extended Data 3C)**. These results indicate that APOE4 promotes the induction of non-vascular myofibroblasts in aged mice in the absence of other AD genetic factors.

Finally, we assessed whether APOE4 is sufficient to induce myofibroblasts in a human context using miBrains, a 3D human brain tissue derived from isogenic APOE3/3 or APOE4/4 iPSCs differentiated into the six major brain cell types^90,91^ (**Figure 1K**). Our laboratory has previously generated a snRNA-seq dataset from isogenic APOE3/3 and APOE4/4 miBrains that captured the major cerebrovascular cell types^91^ (**Extended Data 3D**). Analysis of mural cells revealed two principal subclusters, with preferential enrichment of APOE3/3 cells in mural cluster 1 and APOE4/4 cells in mural cluster 2 (APOE3/3: 92.3% cluster 1, 7.7% cluster 2; APOE4/4: 10.5% cluster 1, 89.5% cluster2; p = 0.039; **Figure 1L, M; Extended Data 3E**). Notably, mural cluster 2 displayed enrichment of extracellular matrix and contractile pathways, closely resembling the transcriptional program observed in human APOE4 brains (**Figure 1N, O, Supplementary Table S13, S14**). A pseudobulk analysis confirmed upregulation of these pathways in APOE4/4 miBrain mural cells (**Extended Data 3F, Supplementary Table S15, S16**). Using markers from our human cerebrovascular atlas, we then assigned pericyte and myofibroblast module scores to the miBrain mural cell subclusters. This revealed a significant shift from a pericyte identity in mural cluster 1 to a myofibroblast identity in mural cluster 2 (p < 0.0001; **Figure 1P**). Consistent with these computational findings, immunofluorescence revealed that APOE4/4 miBrains exhibited a marked reduction in pericyte coverage compared to isogenic APOE3/3 miBrains (0.43 ± 0.056-fold change; 57% decrease; p < 0.0001), accompanied by increased non-vascular αSMA immunoreactivity (3.80 ± 0.43-fold increase; p < 0.0001; **Figure 1Q**). This finding was validated with the additional contractile marker TAGLN, demonstrating increased immunoreactivity in APOE4/4 miBrains (3.77 ± 1.22-fold increase; p = 0.048; **Extended Data 3G**). Together, these results demonstrate that APOE4 is sufficient to promote pericyte loss and the emergence of non-vascular myofibroblasts, recapitulating the transcriptional and morphological features observed in human and mouse APOE4 brains.

### APOE4 expression in mural cells is necessary and sufficient for myofibroblast accumulation

Having identified a robust APOE4-associated myofibroblast phenotype across three model systems, we next sought to determine whether this phenotype arises within mural cells or requires interaction with other APOE4-expressing cell types. To test this, we performed a hybrid genetic mixing experiment in which isogenic APOE3/3 mural cells (iMCs) were selectively integrated into APOE4/4 miBrains **(Figure 2A)**. As expected, APOE4/4 miBrains exhibited significantly reduced pericyte coverage and increased αSMA immunoreactivity compared to isogenic APOE3/3 controls (pericyte coverage: 0.39 ± 0.10-fold change, corresponding to 61% decrease relative to APOE3/3, p = 0.0018; αSMA: 2.99 ± 0.65-fold increase, p = 0.0214; **Figure 2B**). Strikingly, replacing APOE4/4 iMCs with isogenic APOE3/3 iMCs reversed these phenotypes, leading to a significant reduction in αSMA-positive cells and restoration of NG2-positive cell coverage along the vasculature (pericyte coverage: 1.60 ± 0.016-fold increase relative to APOE4/4, p < 0.0001; a-SMA: 0.34 ± 0.14-fold change, corresponding to a 66% decrease relative to APOE4/4, p = 0.0229; **Figure 2B**). These findings indicated that APOE4 expression in mural cells is necessary for pericyte loss and myofibroblast accumulation. We repeated this experiment with iMCs derived from two additional isogenic iPSC lines from an AD and non-AD individual with reciprocal gene editing strategies (Non-AD E3➔E4 [ADRC line] and AD E4➔E3 [sAD line]) (**Extended Data 3H, I**) and found similar results (**Extended Data 3J, K**). To assess whether APOE4 mural cells are sufficient to induce myofibroblasts, we performed the hybrid genetic mixing experiment in reverse, integrating APOE4/4 iMCs into an APOE3/3 miBrain (**Figure 2C**). Notably, this single change significantly reduced pericyte coverage and increased αSMA immunoreactivity to APOE4/4 miBrain levels (pericyte coverage: 0.84 ± 0.027-fold change, corresponding to a 16% decrease relative to APOE3/3, p = 0.0028; a-SMA: 3.21 ± 0.73-fold increase relative to APOE3/3, p = 0.0221; **Figure 2D**). Consistent with these results, immunostaining of iMC monocultures showed significantly higher αSMA and fibronectin levels in APOE4/4 iMCs compared to isogenic APOE3/3 controls across all donor lines (**Extended Data 4A-C**). Finally, we analyzed a previously published snRNA-seq dataset in which human APOE3 (iE3/Cre+) or APOE4 (iE4/Cre+) were conditionally expressed in vascular mural cells of Apoe-KO mice (**Figure 2E**). Isolation of mural cells revealed distinct pericyte and SMC clusters (**Figure 2F**). Similar to our postmortem human brain analysis, DAseq identified a specific SMC subregion enriched in aged iE4/Cre+ mice (**Figure 2G**). This subregion was enriched in genes and pathways related to cellular contraction and the ECM, and unbiased cell type annotation predicted myofibroblast as the top match (**Figure 2H-J, Supplementary Table S17-S19**), highlighting the sufficiency of APOE4 expression in mural cells for this fibrotic phenotype.

**Figure 2.**
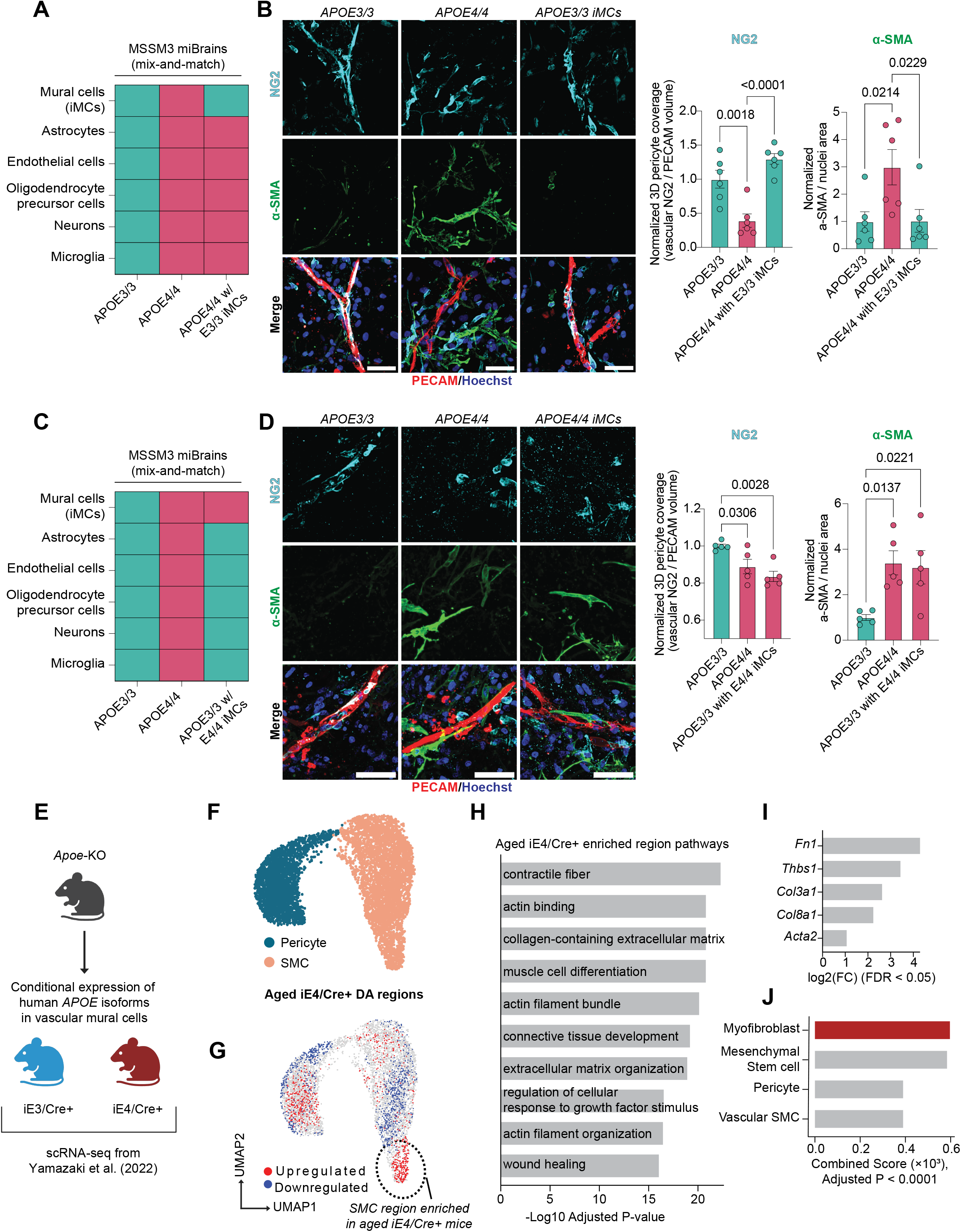
APOE4 expression in mural cells is necessary and sufficient for myofibroblast accumulation. **A.** Schematic of the hybrid genetic experiment in which APOE3/3 iPSC-derived mural cells (iMCs) were integrated into an APOE4/4 miBrain. **B.** Representative images of isogenic APOE3/3, APOE4/4, and APOE4/4 miBrains integrated with APOE3/3 iMCs stained for PECAM, αSMA, and NG2. Scale bar, 50 µm. Data points represent mean values from individual miBrains (n = 6 images per miBrain, n = 6 miBrains per genotype). **C.** Schematic of the hybrid genetic experiment in which APOE4/4 iPSC-derived mural cells (iMCs) were integrated into an APOE3/3 miBrain. **D.** Representative images of isogenic APOE3/3, APOE4/4, and APOE3/3 miBrains integrated with APOE4/4 iMCs stained for PECAM, αSMA, and NG2. Scale bar, 50 µm. Data points represent mean values from individual miBrains (n = 6 images per miBrain, n = 5 miBrains per genotype). For B. and D., three-dimensional pericyte coverage was measured by the overlapping NG2 and CD144 volume, normalized to the total CD144 volume, and expressed relative to APOE3/3. The area positive for αSMA was quantified, normalized to nuclei, and expressed relative to APOE3/3. Bars are mean group values ± SEM. P-values were calculated using a one-way ANOVA with Dunnett’s multiple comparisons test. **E.** Schematic of previously published mouse single-cell RNA-sequencing dataset with conditional human APOE expression in mural cells. **F.** UMAP of mural cells from this dataset. **G.** Differential abundance analysis on the mural cell clusters was performed with DAseq. Red and blue indicate enrichment and depletion in aged iE4/Cre+ mice, respectively. An SMC subregion enriched in aged iE4/Cre+ mice was annotated. **H-I.** Top pathways and representative marker genes of the annotated iE4/Cre+ region, respectively. The MAST package was utilized to identify marker genes of this region compared to other mural cells (|logFC| > 0.75, adjusted p < 0.05). These were then input into ClusterProfiler to identify the top GO pathways. **J.** Top cell type annotations of iE4/Cre+ enriched region. The marker genes from I. were input into EnrichR with the CellMarker database.

Lowering APOE protein levels in APOE4 models is associated with reduced vascular pathology, suggesting that APOE4 exerts a toxic gain-of-function effect^50,92^. To determine whether this same mechanism contributes to myofibroblast accumulation, we reanalyzed the dataset shown in Figure 2E, this time comparing iE4/Cre+ mice with *Apoe*-KO controls. Consistent with a toxic APOE4 gain-of-function, the myofibroblast population was significantly reduced in Apoe-KO mice relative to iE4/Cre+ mice, indicating that APOE expression is required for the APOE4-associated myofibroblast phenotype (0.11 ± 0.02-fold change, corresponding to a 89% decrease relative to iE4/Cre+, p < 0.0001; **Extended Data 4D**). To confirm this *in vitro*, we examined induced mural cell monocultures (iMCs) derived from APOE-KO iPSCs. Treatment of APOE-KO iMCs with recombinant APOE4 protein significantly increased α-SMA and fibronectin immunoreactivity, consistent with acquisition of a myofibroblast phenotype similar to that observed in APOE4/4 iMCs (a-SMA: 1.19 ± 0.059-fold increase, p = 0.0163; fibronectin: 1.50 ± 0.044-fold increase, p < 0.0001; **Extended Data 4E**). Taken together, these findings across multiple model systems point to a causal, toxic role for APOE4 expression in mural cells in driving the expansion of a myofibroblast population.

### APOE4 promotes a pericyte-to-myofibroblast transition

We next investigated how APOE4 mural cells generate the myofibroblast population. In multiple organ systems, including the brain, pericytes differentiate into myofibroblasts following injury^78,82^. A pericyte-to-myofibroblast transition (PMT) would explain the observed correlation between myofibroblast expansion and pericyte loss (**Figure 1D, J, Q**). To explore this possibility, we performed pseudotime analysis of our human cerebrovascular atlas. This revealed two distinct pericyte-derived trajectories: one first terminating in the APOE4-enriched myofibroblast population (pericyte-to-myofibroblast lineage) and another terminating in pericytes (pericyte-to-pericyte lineage) (**Figure 3A**). Gene signature scoring confirmed that the first trajectory was characterized by strong myofibroblast gene activity (AUC = 1.26, p < 0.0001) and the loss of pericyte identity (AUC = -3.01, p < 0.0001) (**Figure 3B**). To determine whether *APOE* genotype influences lineage conversion, we calculated curve weights for both trajectories. APOE4 mural cells were significantly enriched along the pericyte-to-myofibroblast trajectory, whereas APOE3/3 cells preferentially aligned with the pericyte-to-pericyte trajectory (p = 0.004; **Figure 3C**).

**Figure 3.**
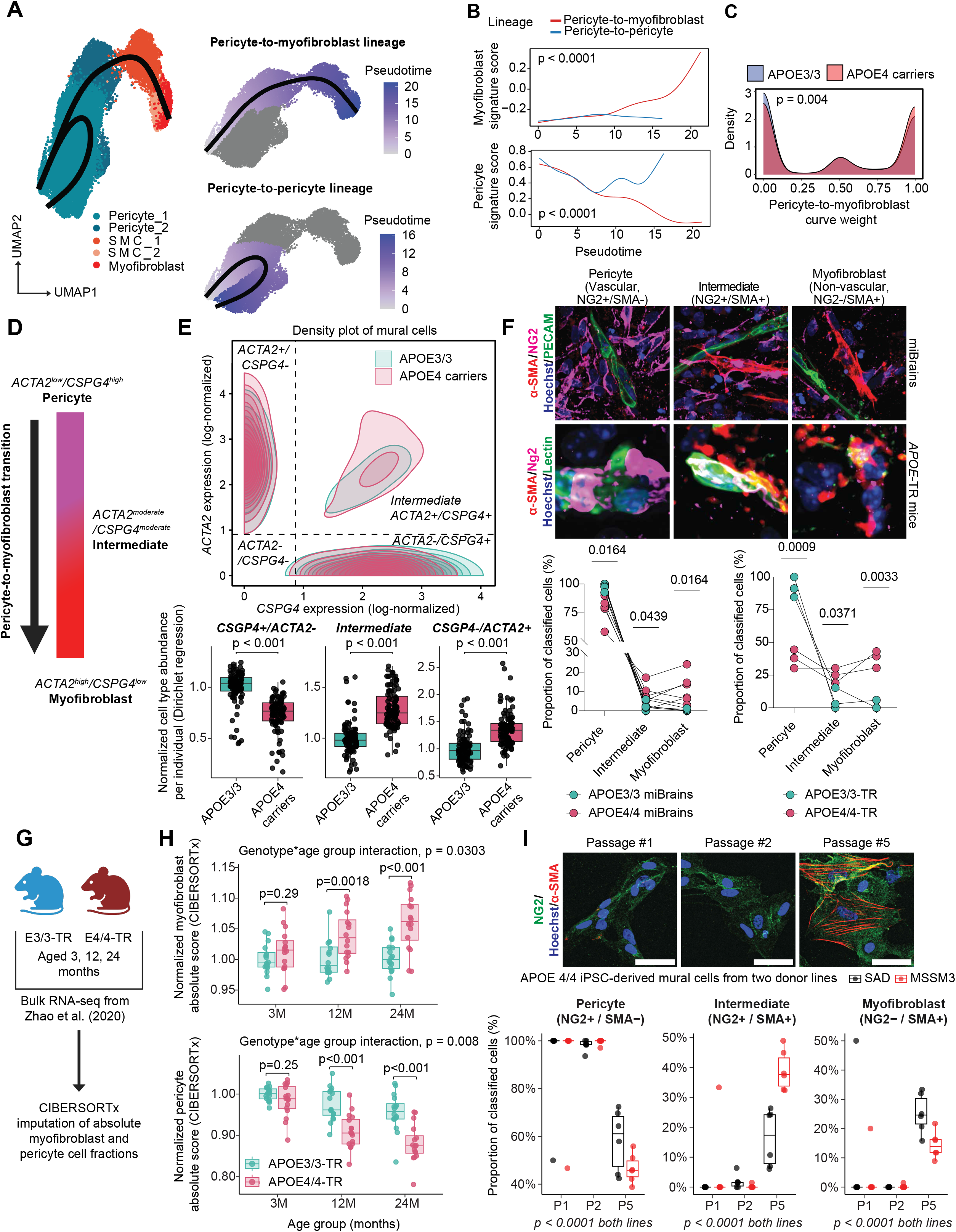
APOE4 promotes a pericyte-to-myofibroblast transition. **A.** Pseudotime analysis of the mural cell subclusters identified in Figure 1 using the slingshot R package. **B.** Generalized additive models (GAMs) were fit along each lineage using curated myofibroblast- and pericyte-associated gene signatures (see Methods) to validate the Slingshot-inferred trajectories. Signature score trajectories were then compared between lineages by computing the area under the curve (AUC) across pseudotime, and significance was assessed using a permutation-based p-value. **C.** Lineage bias between APOE4 carriers and APOE3/3 individuals. Distributions of Slingshot-derived curve weights were plotted for each lineage, stratified by APOE genotype. A value of 0 indicates a cell is exclusive to the pericyte-to-pericyte lineage. A value of 1 indicates a cell is exclusive to the pericyte-to-myofibroblast lineage. The p-value was calculated using the differentiationTest function in the condiments R package (see Methods). **D.** Schematic of *ACTA2* and *CSPG4* gene expression gradient during the pericyte-to-myofibroblast transition. **E.** Density plot of mural cells in non-AD individuals stratified by *ACTA2* and *CSPG4* expression. Cutoffs for gene expression were set as described in the Methods. Subject-level predicted proportions of each mural cell state were obtained from a covariate-adjusted Dirichlet regression model (Extended Data 5A). Counterfactual predictions were generated for each individual under APOE3 and APOE4 while holding covariates constant. Points represent individuals (n = 100 APOE3/3, n = 102 APOE4 carriers); boxes show the distribution of predicted values. Values were normalized to APOE3/3. P-values were calculated by testing whether the mean per-subject APOE4–APOE3 predicted difference differed from zero (Benjamini–Hochberg corrected for multiple states). **F.** Representative images of the pericyte-to-myofibroblast transition in APOE4/4 miBrains and TR-mice. APOE3/3 and APOE4/4 miBrains and TR-mice were stained for a vascular marker (CD144/lectin), αSMA, and NG2. Cells were classified into pericyte (vascular NG2+/ αSMA-), intermediate (NG2+/ αSMA+), or myofibroblast (nonvascular NG2-/ αSMA+) categories using intensity-based thresholds in CellProfiler. The number of cells in each category was normalized to the total number of classified cells. Data points represent mean values from individual miBrains or mice (n = 6 images per miBrain/mouse, n = 6 miBrains and n = 3 mice per genotype). P-values represent comparisons between genotypes for each cell state. P-values were calculated using a two-way repeated measures ANOVA with Geisser-Greenhouse correction and Tukey’s multiple comparisons test. **G.** Schematic of cell-type deconvolution of mouse bulk RNA-seq dataset via CIBERSORTx. **H.** Pericyte and myofibroblast absolute scores across ages and APOE-TR genotypes. CIBERSORTx absolute scores were computed as described in the Methods. Data points represent values from individual mice (n = 16 mice per age group and genotype). Boxes are group medians and IQRs. P-values were calculated as described in the Methods. **I.** Representative images of APOE4/4 iPSC-derived mural cells from two different donors stained for αSMA and NG2 over time. Scale bar, 50 µm. Cells were classified into pericyte (NG2+/ αSMA-), intermediate (NG2+/ αSMA+), or myofibroblast (NG2-/ αSMA+) categories using intensity-based thresholds in CellProfiler. The number of cells in each category was normalized to the total number of classified cells. Data points represent mean values from individual wells (n = 6 images per well, n = 6 wells per line and time point). P-values were calculated within each donor line using a binomial generalized linear model comparing Early (P1–P2) and Late (P5) stages using per-well counts (cells in each state relative to the total classified cells), with Holm correction for multiple testing.

In vivo, the pericyte-to-myofibroblast transition is reported to occur as a gradual continuum, rather than a binary switch, with cells progressively shifting from a pericyte to a myofibroblast identity^78,93^. This process includes an intermediate state in which cells co-express *CSPG4* (pericyte marker) and *ACTA2* (SMC/myofibroblast marker) (**Figure 3D**)^78,93^. We reasoned that if APOE4 promotes a PMT, APOE4 carriers should show not only pericyte loss and a gain of myofibroblasts, but also an enrichment of these intermediate *CSPG4+/ACTA2+* mural cells. To test this, we identified double-positive cells in our cerebrovascular atlas using defined gene expression cutoffs. APOE4 carriers exhibited a higher abundance of double-positive *CSPG4+/ACTA2+* cells compared to APOE3/3 individuals (**Figure 3E**). To confirm this enrichment, we modeled this proportion of each mural cell classification per individual using Dirichlet regression, with *APOE* genotype, AD diagnosis, age, sex, and dataset origin as covariates. We also included APOE x AD status as an interaction term test whether disease diagnosis modified the effect of APOE4. After covariate adjustment, APOE4 was associated with a significant shift in mural cell composition. Model-estimated proportions showed a reduction in CSPG4+/ACTA2− and an increase in CSPG4−/ACTA2+ cells, while intermediate CSPG4+/ACTA2+ cells were also significantly enriched in APOE4 carriers (1.29-fold increase, BH-adjusted p < 0.001; **Figure 3E**), supporting an APOE4-driven pericyte-to-myofibroblast transition. Notably, the intermediate CSPG4⁺/ACTA2⁺ mural cell population showed a higher model-estimated proportion in APOE4 individuals without AD compared to those with AD (**Extended Data 5A**), suggesting that APOE4-associated vascular remodeling may occur prior to overt neurodegenerative pathology.

To determine whether this intermediate state is also present in our experimental models, we quantified mural cells in miBrains and APOE-TR□mice as pericytes (vascular NG2+/αSMA–), intermediate cells (NG2+/αSMA+), or myofibroblasts (non-vascular□NG2–/αSMA+). To avoid misclassification, analyses were restricted to small vessels (< 6 μm) lacking SMCs, focusing on non-vascular αSMA+ cells. Consistent with our atlas findings, APOE4/4 miBrains and APOE4/4-TR mice exhibited a significant reduction in pericyte abundance (miBrains: 80.55 ± 4.90% APOE4/4 vs. 97.68 ± 1.08% APOE3/3, p = 0.0116; APOE-TR mice: 37.56 ± 4.09% APOE4/4-TR vs. 91.93 ± 4.46% APOE3/3-TR, p = 0.0009; **Figure 3F**). This decrease coincided with a significant increase in APOE4/4 myofibroblast abundance that was absent in isogenic APOE3/3 control (miBrains: 11.44 ± 3.14% APOE4/4 vs. 0.33 ± 0.22% APOE3/3, p = 0.0164; APOE-TR mice: 37.68 ± 3.72% APOE4/4 vs. 1.96 ± 1.96% APOE3/3, p = 0.0033; **Figure 3F**). Notably, in APOE4 mice and miBrains we also observed cells along small cerebral vessels strongly co-expressing αSMA and NG2, reflective of an intermediate transition state associated with the PMT (miBrains: 8.02 ± 2.25% APOE4/4 vs. 1.99 ± 0.88% APOE3/3, p = 0.0439; APOE-TR mice: 24.76 ± 3.28% APOE4/4 vs. 6.11 ± 4.72% APOE3/3, p = 0.0371; **Figure 3F**). To further validate the occurrence of a pericyte-to-myofibroblast transition in our experimental models, we quantified the average distance of each cell type from the vasculature in both APOE4/4 miBrains and APOE4/4-TR mice. In both model systems, vascular distance increased in a stepwise manner from pericytes to intermediate cells to myofibroblasts, supporting a model in which pericytes detach from vessels and migrate into the parenchyma while acquiring myofibroblast characteristics (APOE4/4 miBrains: R^2^ = 0.53, p = 0.0009; APOE4/4-TR mice: R^2^ = 0.81, p = 0.0028; **Extended Data 5B**). Taken together, these results support that APOE4 promotes a stepwise pericyte-to-myofibroblast transition in both human and *in vivo* models.

Given our finding that APOE4 promotes myofibroblast characteristics in iMC monocultures (**Extended Data 4A-C**), we also examined the APOE4-induced PMT in this context. Using CIBERSORTx^94^ deconvolution of a bulk RNA-seq dataset^50^, we mapped iMCs onto mural cell subclusters from our cerebrovascular atlas **(Extended**□**Data**□**5C)**. Without stratifying by *APOE* genotype, we found that iMCs transcriptionally align most with pericytes compared to other mural cell subtypes (raw data from previous study^50^, **Extended Data 5D**). However, investigating cell type identity scores between genotypes revealed two significant shifts: a loss of pericyte identity and a gain of myofibroblast identity in APOE4/4 iMCs relative to APOE3/3 (pericyte: 0.90 ± 0.010-fold change, corresponding to a 10% decrease relative to APOE3/3, adjusted p = 0.0286; myofibroblast: 1.66 ± 0.029-fold increase, adjusted p = 0.0005; raw data from previous study ^50^, **Extended Data 5E**). Differential expression analysis further confirmed upregulation of canonical myofibroblast pathways in APOE4/4□iMCs (raw data from previous study^50^, **Extended**□**Data**□**5F, Supplementary Table S20, S21)**. Together, these findings are consistent with an APOE4-mediated PMT.

Next, if myofibroblasts arise from pericytes rather than representing a pre-existing population, we would expect to observe a progressive shift in mural cell states over time. In the conditional APOE mouse dataset (**Figure 2E**), we observed an age-dependent decrease in pericyte abundance and age-dependent increase in myofibroblast abundance in iE4/Cre+ mice (pericyte: age x genotype p = 0.002; myofibroblast: age x genotype p = 0.007; **Extended Data 5G**). We then performed CIBERSORTx deconvolution of a longitudinal bulk RNA-seq dataset from APOE-TR mice^95^ using pericyte and myofibroblast marker genes derived from the conditional APOE mouse dataset (**Figure 3G**). At 3 months of age, APOE3-TR and APOE4-TR mice showed no significant differences in either pericyte or myofibroblast abundance. However, by 12 months APOE4-TR mice exhibited a reduction in pericyte signatures accompanied by a corresponding increase in myofibroblast signatures, which became more pronounced at 24 months. This was supported by a significant genotype-by-age interaction for both populations (myofibroblast: interaction p = 0.0303; pericyte: interaction p = 0.008; **Figure 3H**).

To further test whether pericytes give rise to myofibroblasts, we next examined differentiation potential within the miBrain snRNA-seq dataset. If pericytes transition into myofibroblasts, one would expect higher developmental potential in the pericyte state and lower potential in the myofibroblast state. To test this, we applied CytoTRACE2, a deep learning–based framework trained on 34 datasets that estimates developmental potential for each cell, where higher scores indicate less differentiated states. Consistent with a pericyte-to-myofibroblast transition, CytoTRACE2 predicted a gradient of decreasing developmental potential from the pericyte cluster toward the myofibroblast cluster (**Extended Data 5H**). We then integrated these scores with the k-nearest neighbor graph using CellRank2 to infer differentiation directionality, which revealed a clear trajectory from cells expressing pericyte signatures toward cells coexpressing myofibroblast signatures (**Extended Data 5I**).

Finally, to provide additional evidence that APOE4 pericytes convert to myofibroblasts in our model systems, we performed temporal analysis on iMC monocultures. In young cultures (passages one and two), nearly 100% of classified APOE4/4 iMCs were NG2+/αSMA-, consistent with a pericyte identity. However, in aged cultures (passage five), APOE4/4 iMCs significantly upregulated αSMA expression, shifting from a pure pericyte population to intermediate and myofibroblast states. These results were consistent across two donor iPSC lines (**Figure 3I**).

Collectively, these results across human tissue, iPSC-derived models, and APOE-TR□mice demonstrate that APOE4 expression in mural cells drives a pericyte-to-myofibroblast transition.

### Myofibroblast-derived fibronectin increases amyloid accumulation in APOE4 models

Myofibroblasts are key mediators of fibrosis, functioning as major sources of extracellular matrix (ECM) proteins, particularly collagens and fibronectin (*FN1*)^96^. Excessive ECM deposition by myofibroblasts leads to pathological tissue remodeling and organ dysfunction^97^. Recent studies have identified genetic variants in *FN1* associated with cognitive resilience in APOE4 carriers^57^, suggesting that altered fibronectin biology may influence disease outcomes in APOE4 carriers. We therefore examined whether APOE4-associated myofibroblasts promote fibronectin deposition in the brain vasculature. In APOE4/4-TR□mice, fibronectin immunoreactivity was significantly increased along small vessels compared to age- and sex-matched APOE3/3-TR□controls (2.02□±□0.19-fold increase,□p□=□0.0028), coinciding with elevated αSMA□staining (2.39□±□0.52-fold increase,□p□=□0.0447; **Figure**□**4A**). Consistent with these findings, APOE4/4 miBrains also exhibited increased fibronectin accumulation along the vasculature relative to isogenic APOE3/3 miBrains (1.56 ± 0.16-fold increase, p = 0.0058; **Figure 4B**). We confirmed the presence of vascular fibrosis in our APOE4 models using two additional ECM proteins, collagen I and collagen III, finding increased deposition of both proteins in APOE4/4-TR mice and APOE4/4 miBrains (**Extended Data 6A, B**).

**Figure 4.**
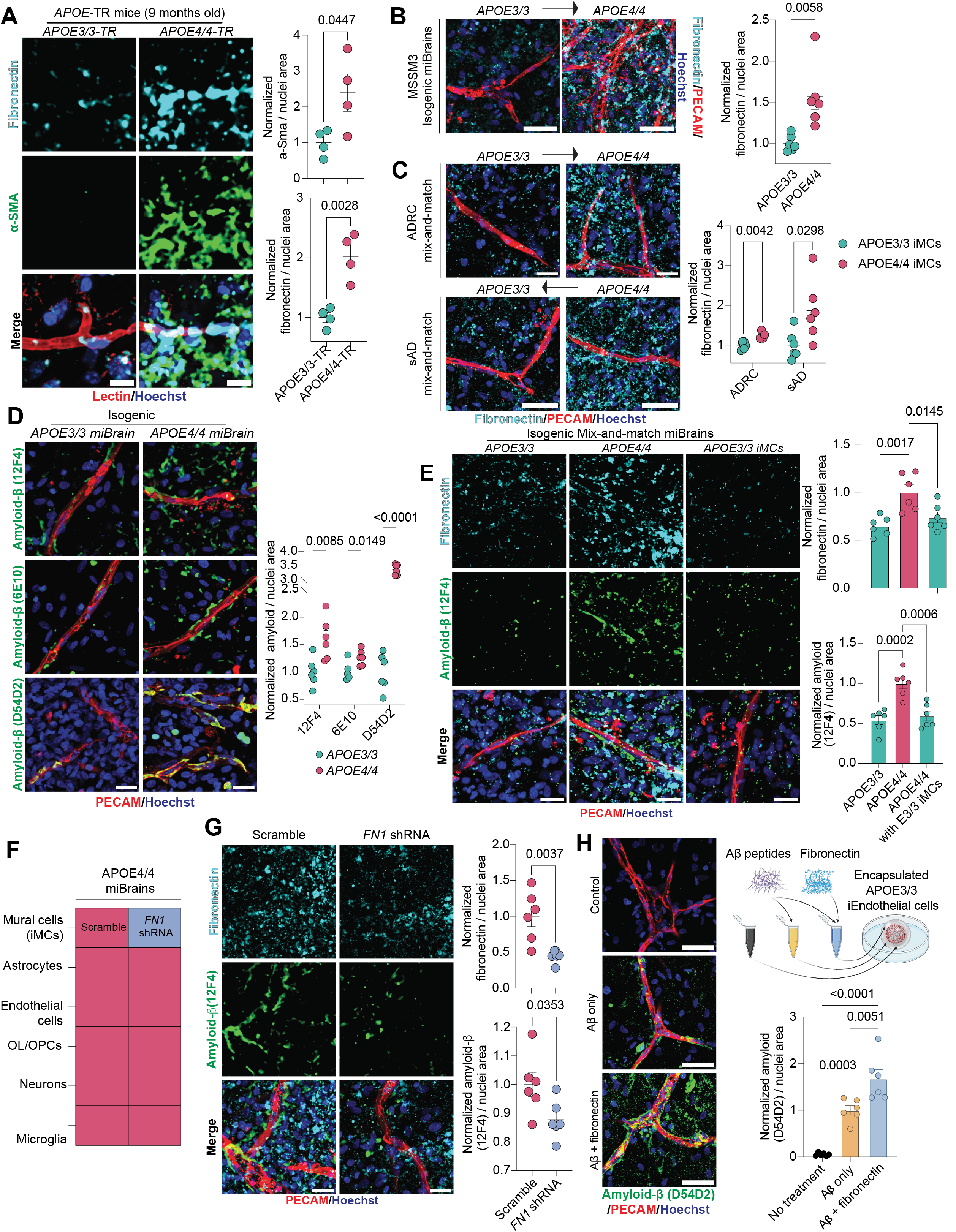
Myofibroblast-derived fibronectin drives amyloid accumulation in APOE4 models. **A.** Representative images of small vessels in the corpus callosum of APOE3/3-TR and APOE4/4-TR mice stained for lectin, αSma, and fibronectin. Scale bar, 10 µm. For visualization only, a median filter was applied to the fibronectin and αSma channels. **B.** Representative images of APOE3/3 and APOE4/4 miBrains stained for PECAM and fibronectin. Scale bar, 50 µm. **C.** miBrain experiments performed with additional isogenic pairs of iPSC-derived mural cells (iMCs) as previously described in Extended Figure 3I. ADRC scale bar, 25 µm. SAD scale bar, 50 µm. **D.** Representative images of isogenic pairs of APOE3/3 and APOE4/4 miBrains stained for three amyloid antibodies targeting different epitopes of the amyloid-β peptide. Scale bar, 25 µm. For visualization only, a median filter was applied to the amyloid channels. **E.** Representative images of isogenic APOE3/3, APOE4/4, and APOE4/4 miBrains integrated with APOE3/3 iMCs stained for PECAM, fibronectin, and amyloid (12F4). Scale bar, 25 µm. **F.** Schematic of *FN1* knockdown experiment in APOE4/4 miBrains. **G.** Representative images of APOE4/4 miBrains integrated with either iMCs transduced with scramble shRNA or shRNA targeting *FN1* stained for fibronectin and amyloid (12F4). Scale bar, 25 µm. **H.** Treatment of encapsulated APOE3/3 iPSC-derived endothelial cells (iECs) with fibronectin and amyloid mixtures. Scale bar, 50 µm. For the control group, iECs were left untreated. For the experimental groups, iECs were treated with either 20 nM amyloid-β 1-40 and 1-42 or 20 nM amyloid-β 1-40 and 1-42 mixed with 50 ng/mL fibronectin for 96 hrs. iECs were then fixed and stained for amyloid (D54D2). For all panels, the area positive for αSMA, fibronectin, or amyloid immunoreactivity was quantified, normalized to nuclei, and expressed relative to APOE3/3 (panels A, B, C), APOE4/4 (panels E, G), or iEC amyloid treatment (panel H). Data points represent mean values from individual miBrains (n = 6 images per miBrain, n = 4-6 miBrains per condition), mice (n = 6 images per mouse, n = 4 mice per genotype), or iECs (n = 6 images per iECs, n = 6 iECs per condition). Bars are mean group values ± SEM. Unless stated otherwise, p-values were calculated using unpaired two-sample Student’s t-tests (panels A, B, C, G) or a one-way ANOVA with Dunnett’s multiple comparisons test (panel E, H).

Myofibroblasts were the largest contributor of *FN1* expression among individuals from our cerebrovascular atlas (**Extended Data 6C**). Furthermore, most of the fibronectin deposition in APOE4/4-TR mice occurred near αSMA+ regions (72.66 ± 1.90%, p = 0.0013, **Extended Data 6D**), implicating myofibroblasts as a causal source of vascular fibrosis. To directly test this, we swapped APOE4/4 mural cells to APOE3/3 in an APOE4/4 miBrain, an approach that previously reversed APOE4-associated myofibroblast phenotypes (**Figure 2B**). Remarkably, this single swap led to a significant reduction in fibronectin protein across multiple donor lines (**Figure 4C**), suggesting that APOE4 mural cell-derived myofibroblasts are the main contributor to fibronectin accumulation.

In the central nervous system, perivascular fibroblasts are also a major source of fibronectin and contribute to fibrotic scar formation in response to brain injury^98,99^. To rule out the involvement of this cell type, we analyzed perivascular fibroblasts from the original snRNA-seq datasets corresponding to the same subjects included in our cerebrovascular atlas. We found that the myofibroblasts identified in this study exhibited significantly (p < 0.0001) higher *FN1* expression compared to fibroblasts (**Extended Data 6E**), and that *APOE* genotype had no significant (FDR > 0.05) effect on ECM-related pathways in fibroblasts (**Supplementary Table S22, S23**), further highlighting mural cell-derived myofibroblasts as the primary cell type responsible for APOE4-associated vascular ECM remodeling.

To assess how mural cell *FN1* expression relates to clinical characteristics of AD, we analyzed our cerebrovascular atlas using the Haney (2024) dataset^11^, which includes extensive clinical annotations. We focused on individuals with an AD diagnosis and pseudobulked mural cell *FN1* expression for each individual to specifically examine *FN1* within the context of the pericyte-to-myofibroblast transition. Consistent with our earlier findings, APOE4 carriers had significantly higher mural cell *FN1* expression compared to APOE3/3 individuals (1.39 ± 0.10-fold increase, p = 0.0065; **Extended Data 6F**). Supporting a role for fibronectin in AD progression, mural cell *FN1* expression was negatively correlated with the age of AD diagnosis (β = -8.77, p = 0.0007, **Extended Data 6G**). We also derived a global pathology score for each individual by averaging their total amyloid plaque, neurofibrillary tangle, Lewy body, and cerebral infarct scores reported in the original dataset. Individuals in the highest quartile of global pathology had significantly elevated mural cell *FN1* expression compared to the remaining individuals (1.43 ± 0.15-fold increase relative to the low pathology group, p = 0.0041; **Extended Data 6H**), suggesting a link between APOE4-driven fibronectin deposition and AD pathology.

To investigate the causal role of fibronectin in AD pathology, we leveraged the miBrain system. Because APOE4 carriers are highly susceptible to vascular amyloid deposition and fibronectin can directly bind amyloid^100–102^, we focused on perivascular amyloid-β accumulation. To first validate the miBrain as a model for this pathology, we immunostained isogenic APOE3/3 and APOE4/4 miBrains using three separate antibodies each targeting distinct amyloid−β epitopes. Consistent with observations in APOE4 carriers, APOE4/4 miBrains showed significantly increased amyloid-β immunoreactivity along the vasculature with all three antibodies compared to isogenic APOE3/3 controls (12F4: 1.61 ± 0.16-fold increase, p = 0.0085; 6E10: 1.26 ± 0.056-fold increase, p = 0.0149; D54D2: 3.38 ± 0.074-fold increase, p < 0.0001; **Figure 4D**).

If fibronectin causally contributes to amyloid-β pathology, then reducing myofibroblast-derived fibronectin should also lower amyloid accumulation. Consistent with this, APOE4/4 miBrains containing APOE3/3 iMCs displayed significantly reduced vascular amyloid deposition (12F4 antibody) compared to all APOE4/4 controls (0.59 ± 0.06-fold change, corresponding to a 41% decrease relative to APOE4/4, p = 0.0006; **Figure 4E**). This finding was validated with an additional amyloid antibody (D54D2) (**Extended Data 6I**). We also performed this genetic hybrid mixing experiment in reverse as previously described (**Figure 2C**), integrating APOE4/4 iMCs into an APOE3/3 miBrain. Strikingly, APOE4/4 iMCs were sufficient to increase both fibronectin and amyloid deposition (fibronectin: 1.15 ± 0.03-fold increase relative to APOE3/3, p = 0.0092; D54D2: 1.44 ± 0.18-fold increase relative to APOE3/3, p = 0.0335; **Extended Data 6J**). Together, these bidirectional experiments suggest that APOE4 expression in mural cells is both sufficient and necessary to drive vascular fibrosis and amyloid accumulation. Notably, we also observed increased APOE deposition along the vasculature of APOE4/4 miBrains, with significantly enhanced colocalization of APOE with both fibronectin and amyloid, suggesting that APOE may directly associate with the fibrotic ECM to promote amyloid deposition (**Extended Data 6K-L**).

Fibronectin directly binds to amyloid and may promote its aggregation^100^. To determine whether the reduction in amyloid pathology observed when APOE3/3 iMCs replaced APOE4/4 iMCs in the miBrain was mediated by decreased fibronectin, we transduced APOE4/4 iMCs with a shRNA targeting *FN1* or a scramble control. qRT-PCR of iMC monocultures confirmed robust *FN1* knockdown (0.037 ± 0.0018-fold change, corresponding to a 96.3% decrease relative to scramble, p = 0.0009; **Extended Data 6M**). FN1 knockdown and control APOE4/4 iMCs were then incorporated into APOE4/4 miBrains to directly assess the impact of fibronectin depletion on amyloid accumulation (**Figure 4F**). Consistent with our hypothesis, *FN1* knockdown led to markedly reduced fibronectin staining and a concomitant decrease in vascular amyloid deposition compared to scramble controls (fibronectin: 0.44 ± 0.036-fold change, corresponding to a 56% decrease relative to scramble, p = 0.0037; amyloid: 0.88 ± 0.027-fold change, corresponding to a 12% decrease relative to scramble, p = 0.0353; **Figure 4G**). To further validate the causal link between fibronectin and amyloid accumulation, we treated APOE3/3 iPSC-derived endothelial cells with amyloid alone or an amyloid-fibronectin mixture. Cells exposed to the amyloid-fibronectin mixture exhibited significantly greater amyloid accumulation than those treated with amyloid alone (1.68 ± 0.20-fold increase, p = 0.0051, **Figure 4H**). Together, these findings demonstrate that fibronectin secreted by APOE4-derived myofibroblasts promotes vascular amyloid accumulation, establishing a direct mechanistic link between myofibroblasts and AD progression in APOE4 carriers.

### Increased TGF-β signaling in the APOE4 brain promotes the pericyte-to-myofibroblast transition

Having identified a key role for APOE4-derived myofibroblasts in AD pathology, we next investigated the molecular mechanisms through which APOE4 promotes the PMT. We applied NicheNet^103^, a computational tool that predicts upstream ligands regulating target gene expression, using APOE4 myofibroblast marker genes identified in our human cerebrovascular atlas as input (**Figure 5A**, **Figure 1H**). NicheNet identified TGF-β as the top predicted driver of the PMT (**Figure 5B, Supplementary Table S24**). In addition to TGF-β, NicheNet also identified other candidate ligands, including BMP and FGF family members, although these ranked lower and showed less consistent regulatory potential across myofibroblast-associated targets. In agreement with its established role in fibrosis, TGF-β ligands were predicted to regulate multiple myofibroblast-associated genes, including *FN1*, *ACTA2*, and *COL1A2* (**Figure 5C, D**). Fibrosis in the CNS can also arise downstream of an interferon-gamma-mediated cascade, particularly in neuroinflammatory diseases such as multiple sclerosis^98^. However, APOE4 myofibroblasts show no change in interferon-gamma signaling at either the gene or pathway level (**Supplementary Table S10-S12**). These findings suggest that APOE4 cerebrovascular fibronectin and amyloid deposition occur through a TGF-β-dependent, IFN-γ-independent mechanism.

**Figure 5.**
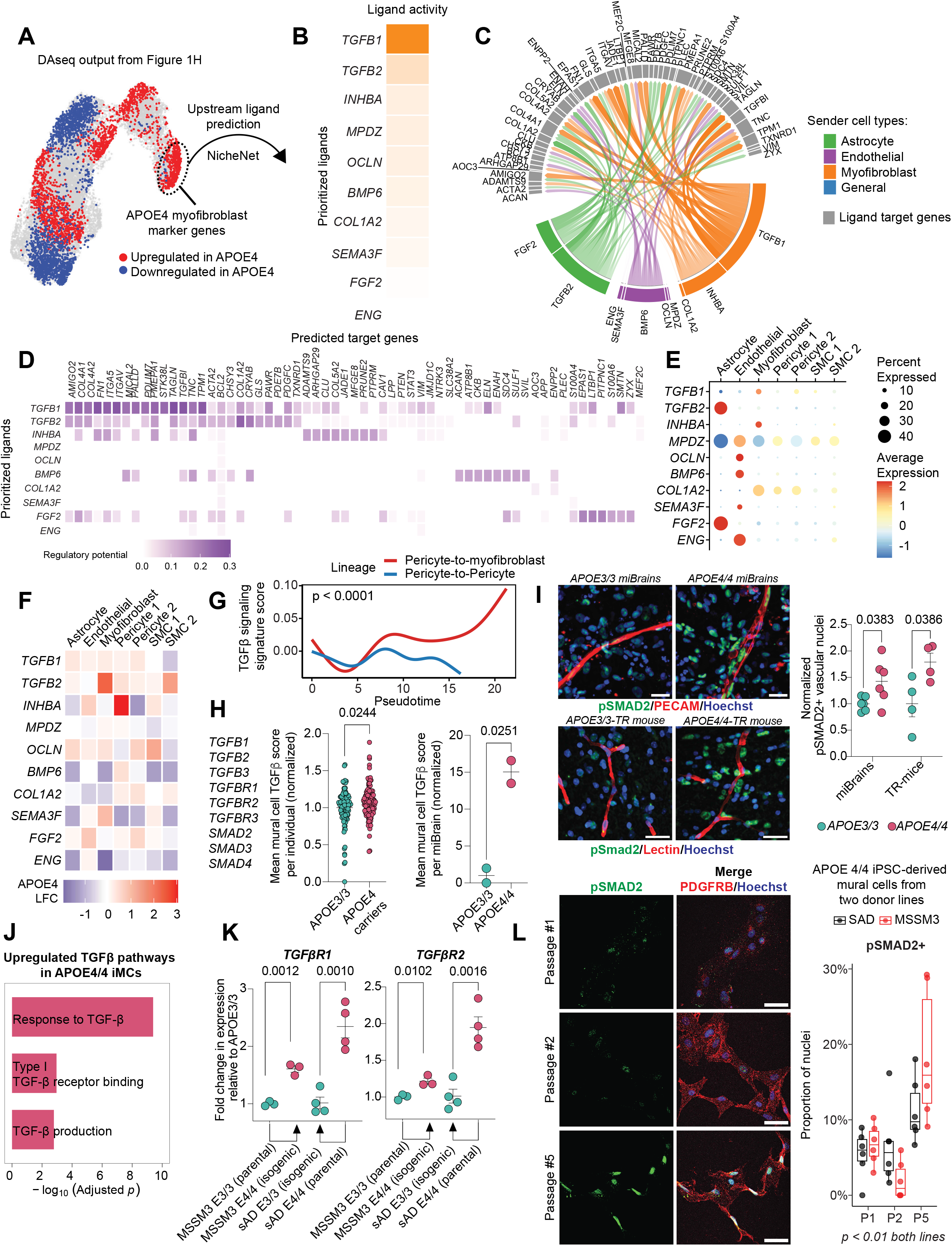
Increased TGF-β signaling in the APOE4 brain drives the pericyte-to-myofibroblast transition. **A.** APOE4 myofibroblast marker genes identified in Extended Data 2H were input into the R package NicheNet. **B.** Top predicted ligands driving myofibroblast marker expression. **C.** Cirkos plot of top ligands and their predicted target genes from NicheNet. **D.** Predicted target genes of the top ligands in B. **E.** Gene expression of top ligands per cell type. **F.** Ligand expression of APOE4 carriers relative to APOE3/3 per cell type. **G.** A generalized additive model (GAM) was fitted along each lineage using the hallmark TGF-β signaling gene set (see Methods). Signature score trajectories were then compared between lineages by computing the area under the curve (AUC) across pseudotime, and significance was assessed using a permutation-based p-value. **H.** TGF-β expression scores of mural cells from the cerebrovascular atlas and miBrain snRNA-seq datasets stratified by APOE genotype. TGF-β module scores were calculated per-cell using Seurat’s *AddModuleScore()* function based on the specified genes. The scores were then averaged per individual or miBrain. Data points represent individuals (n = 106 APOE3/3, n = 106 APOE4 carriers) or miBrains (n = 2 per genotype). Bars are mean group values ± SEM. P-values were calculated using unpaired two-sample Student’s t-tests with Welch’s correction. **I.** Representative images of APOE3/3 and APOE4/4 miBrains and TR-mice stained for a vascular marker (PECAM/lectin) and pSMAD2. Scale bar, 25 µm. For visualization only, vessels were cropped and a median filter was applied. Nuclei along the vasculature were classified as pSMAD2+ using an intensity-based threshold in CellProfiler, normalized to the total number of vascular nuclei, and expressed relative to APOE3/3. Data points represent mean values from individual miBrains or mice (n = 6 images per miBrain/mouse, n = 6 miBrains and n = 4 mice per genotype). Bars are mean group values ± SEM. P-values were calculated using unpaired two-sample Student’s t-tests. **J.** Upregulated TGF-β signaling GO pathways in APOE4/4 iPSC-derived mural cells (iMCs). Differentially expressed genes in the iMC bulk RNA-seq dataset (|logFC| > 0.25, adjusted p < 0.05 calculated from limma) were input into ClusterProfiler. The significantly upregulated TGF-β signaling GO pathways (FDR < 0.05) were then extracted for graphical representation. **K.** *TGF*β*R* expression in isogenic pairs of iMCs from two donor lines. MSSM3 line: FPKM values from iMC bulk RNA-seq dataset normalized to APOE3/3. Data points represent individual bulk RNA-sequencing replicates (n = 3 per genotype). Bars are mean group values ± SEM. Adjusted p-values were calculated using the R package limma to account for multiple comparisons. sAD line: *TGF*β*R* expression in iMCs via qRT-PCR relative to APOE3/3. Data points represent mean values from individual wells of iMCs (3 qRT-PCR replicates per well, 4 wells per genotype). Bars are mean group values ± SEM. P-values were calculated using unpaired two-sample Student’s t-tests. **L.** Representative images of APOE4/4 iPSC-derived mural cells from two different donors stained for pSMAD2 and PDGFRβ over time. Scale bar, 50 µm. Nuclei were classified as pSMAD2+ using an intensity-based threshold in CellProfiler and normalized to the total number of nuclei. Data points represent mean values from individual wells (n = 6 images per well, n = 6 wells per line and time point). P-values were calculated within each donor line using a binomial generalized linear model comparing Early (P1–P2) versus Late (P5) using per-well pSMAD2+ nuclei counts relative to total nuclei, with Holm correction for multiple testing.

Within our postmortem human cerebrovascular atlas, TGFβ1 was predominantly expressed by myofibroblasts and TGFβ2 by astrocytes, with only modest differences in ligand expression between APOE4 and APOE3/3 individuals (**Figure 5C, E, F**). However, TGF-β pathway activity increased along the inferred pericyte-to-myofibroblast trajectory. Specifically, TGF-β hallmark gene signature scoring along pseudotime revealed significantly greater pathway activation along the pericyte-to-myofibroblast lineage compared to the pericyte-to-pericyte lineage in human mural cells (AUC = 0.479, p < 0.0001; **Figure 5G**). A similar pattern was observed in miBrain mural cells, where TGF-β signaling scores increased along the CytoTRACE-derived differentiation gradient from cells expressing pericyte signatures toward those expressing myofibroblast signatures (r = 0.38, p < 0.0001, **Extended Data 7A, 5H-I**). Consistent with this trajectory-based activation, mural cells from both APOE4 carriers in human tissue and APOE4/4 miBrains exhibited significant upregulation of key TGF-β pathway genes relative to APOE3/3 controls (human snRNA-seq: 1.08 ± 0.02-fold increase, p = 0.0244; miBrain snRNA-seq: 15.03 ± 1.54-fold increase, p = 0.0251; **Figure 5H**). The APOE4 vs. APOE3 logFC for each individual gene used in this TGF-β score calculation is displayed in Extended Data 7B. To functionally validate the increase in TGF-β signaling along the APOE4 vasculature, we immunostained miBrains and APOE-TR mice for pSmad2, a canonical marker of TGF-β pathway activation. Consistent with our computational predictions, APOE4/4 miBrains exhibited significantly more pSMAD2+ vascular nuclei compared to APOE3/3 (1.43 ± 0.17-fold increase, p = 0.0383; **Figure 5I**). Similarly, APOE4/4-TR mice showed significantly more pSmad2 immunoreactive nuclei around cortical microvasculature compared to age-matched APOE3/3-TR mice (1.79 ± 0.17-fold increase, p = 0.0386; **Figure 5I**).

We next sought to identify the cellular source of increased TGF-β in APOE4 brains. Given our prior finding that APOE4 mural cells are both necessary and sufficient to generate myofibroblasts, we hypothesized that TGF-β pathway activation is intrinsic to APOE4 mural cells. Supporting this, APOE4/4 iMCs in monoculture exhibited significantly higher nuclear SMAD2 intensities compared to isogenic APOE3/3 controls (1.21 ± 0.060-fold increase, p = 0.0104; **Extended Data 7C**). Moreover, treating APOE3/3 iMCs with recombinant TGFβ1 (50 ng/mL) increased nuclear SMAD2 levels comparable to APOE4/4 cells (1.17 ± 0.016-fold increase, p = 0.0393; **Extended Data 7C**), indicating that elevated baseline TGF-β signaling in APOE4/4 mural cells accounts for their increased SMAD2 activation.

To explore the mechanism underlying this elevated signaling, we analyzed iMC bulk RNA-seq data. Consistent with our immunofluorescence findings, APOE4/4 iMCs exhibited significant upregulation of TGF-β pathway genes (**Figure 5J**). We reasoned that enhanced signaling could result from either increased TGF-β ligand production or upregulation of TGF-β receptors. Analysis of this bulk RNA-seq data revealed that both *TGF*β*R1* and *TGF*β*R2* were significantly upregulated in APOE4/4 mural cells compared to isogenic APOE3/3 controls (*TGF*β*R1:* 1.61 ± 0.057-fold increase, adjusted p = 0.0012; *TGF*β*R2:* 1.22 ± 0.042-fold increase, adjusted p = 0.0102, raw data from previous study^50^, **Figure 5K**). We validated these findings in another iMC isogenic pair via qRT-PCR (*TGF*β*R1*: 2.35 ± 0.20-fold increase, adjusted p = 0.0020; *TGF*β*R2*: 1.95 ± 0.15-fold increase, adjusted p = 0.0020; **Figure 5K**). Finally, to test whether receptor upregulation is functionally linked to myofibroblast induction, we inhibited TGF-β signaling using the SB431542 compound. This treatment significantly reduced αSMA and fibronectin expression (αSMA: 0.39 ± 0.033-fold change, corresponding to a 61% decrease relative to APOE4/4, p < 0.0001; fibronectin: 0.76 ± 0.085-fold change, corresponding to a 24% decrease relative to APOE4/4, p = 0.0308; **Extended Data 7D**).

Finally, we previously found that APOE4 iMCs transition from pericytes into myofibroblasts over time (**Figure 3I**). If TGF-β signaling is critical for this process, we should also observe an increase in TGF-β signaling over time that corresponds with myofibroblast emergence. Indeed, compared to young cultures (passage one and two), aged APOE4/4 iMC cultures (passage 5) displayed significantly higher pSMAD2+ cell proportions (**Figure 5L**).

Taken together, these results indicate that increased expression of TGF-β receptors in APOE4 mural cells drives enhanced TGF-β signaling and promotes the pericyte-to-myofibroblast transition.

### TGF-**β** inhibition reverses APOE4-driven myofibroblast accumulation, cerebrovascular fibrosis, and amyloid deposition

AD patients exhibit significantly elevated *TGF*β*1* mRNA levels that strongly correlate with CAA severity^61^ , and chronic TGF-β overexpression in mice induces cerebrovascular fibrosis and perivascular amyloid accumulation^59,60^. Our findings support a model in which heightened TGF-β signaling in APOE4 brains causes a PMT leading to vascular fibronectin and amyloid deposition. To directly test this mechanism, we inhibited TGF-β signaling in APOE4/4 miBrains using two chemically distinct ALK5 (TGFβR1) inhibitors (SB431452 and Galunisertib) (**Figure 6A**).

**Figure 6.**
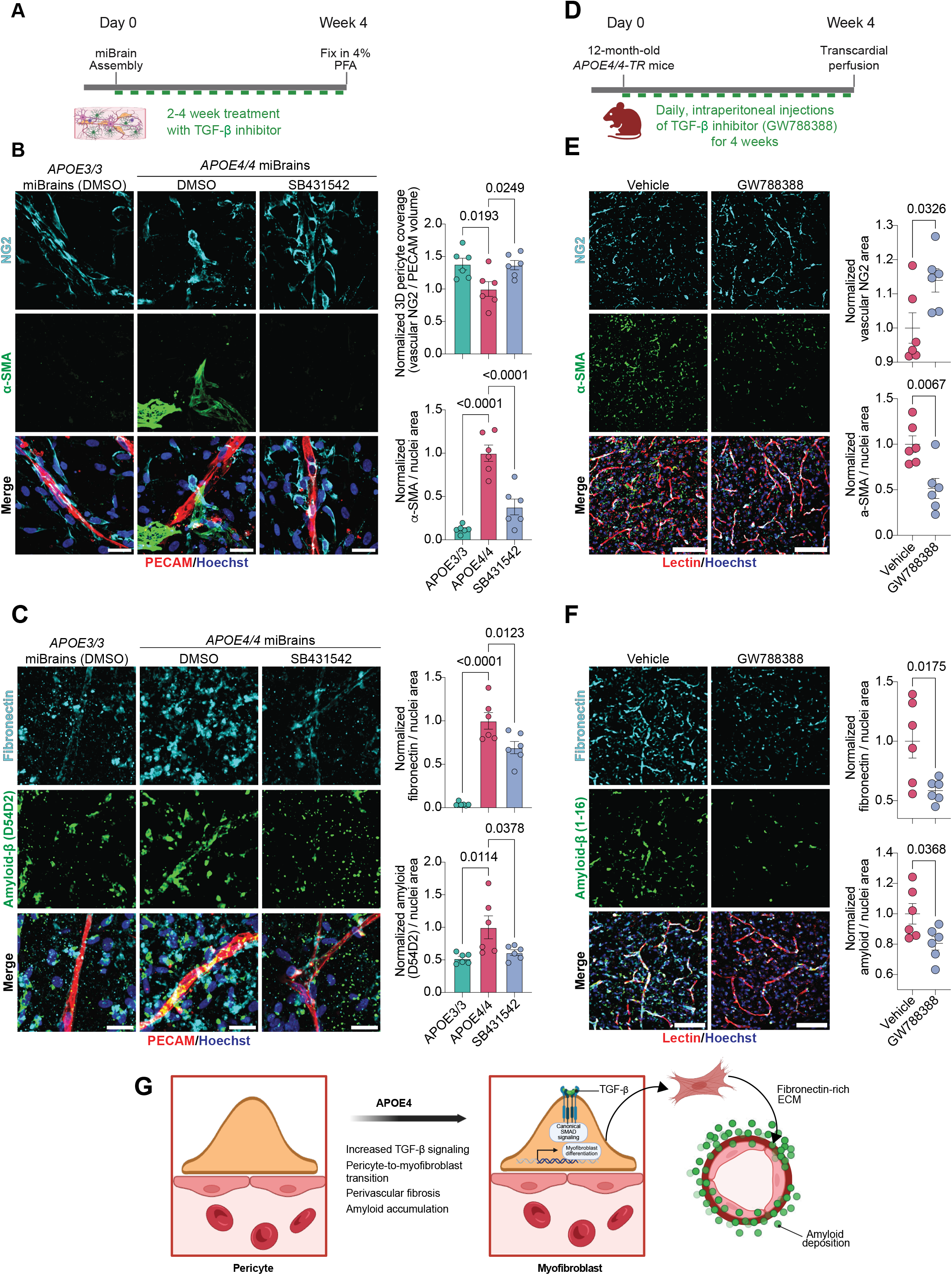
TGF-β inhibition reverses APOE4-driven myofibroblast accumulation, cerebrovascular fibrosis, and amyloid accumulation. **A.** Timeline of TGF-β inhibition in miBrains. **B.** Representative images of myofibroblast phenotypes after TGF-β inhibition in miBrains. Scale bar, 25 µm. For the control group, APOE3/3 and APOE4/4 miBrains were treated with a DMSO vehicle for two weeks. For the experimental group, APOE4/4 miBrains were treated with 10 µM SB431542 for two weeks. miBrains were then fixed and stained for PECAM, αSMA, and NG2. **C.** Representative images of cerebrovascular fibrosis and amyloid deposition in miBrains after TGF-β inhibition. Scale bar, 25 µm. For visualization only, a median filter was applied to the amyloid channel. For the control group, APOE3/3 and APOE4/4 miBrains were treated with a DMSO vehicle for 4 weeks. For the experimental group, APOE4/4 miBrains were treated with 10 µM SB431542 for 4 weeks. miBrains were then fixed and stained for PECAM, fibronectin, and amyloid (D54D2). For panels B and C, three-dimensional pericyte coverage was measured by the overlapping NG2 and PECAM volume, normalized to the total PECAM volume and expressed relative to APOE4/4 vehicle. The area positive for αSMA, fibronectin, or amyloid was quantified, normalized to nuclei, and expressed relative to APOE4/4 vehicle. Data points represent mean values from individual miBrains (n = 6 images per miBrain, n = 6 miBrains per condition). Bars are mean group values ± SEM. P-values were calculated using a one-way ANOVA with Dunnett’s multiple comparisons test. **D.** Timeline of TGF-β inhibition in aged APOE4/4-TR mice. **E.** Representative images of myofibroblast phenotypes after TGF-β inhibition in the cortex of aged APOE4/4-TR mice. Scale bar, 75 µm. Images were deconvolved using identical parameters. For visualization only, a median filter was applied. Mice were either treated with DMSO vehicle or 10 mg/kg GW788388 daily for 4 weeks and brain sections were stained for lectin, αSMA, and NG2. **F.** Representative images of cerebrovascular fibrosis and amyloid-β immunoreactivity in aged APOE4/4-TR mice after TGF-β inhibition. Scale bar, 75 µm. Images were deconvolved using identical parameters. For visualization only, a median filter was applied to the fibronectin and amyloid channels. Mice were either treated with DMSO vehicle or 10 mg/kg GW788388 daily for 4 weeks and brain sections were stained for lectin, fibronectin, and amyloid-β. For panels E and F, pericyte coverage was measured by quantifying the overlapping Ng2 and lectin area, normalized to vehicle. The area positive for αSma, fibronectin, or amyloid was quantified, normalized to nuclei, and expressed relative to vehicle. Data points represent mean values from individual mice (n = 6 images per mouse, n = 6 mice per condition). Bars are mean group values ± SEM. P-values were calculated using unpaired two-sample Student’s t-tests. **G.** Mechanistic model of the TGF-β-mediated pericyte-to-myofibroblast transition in APOE4 carriers.

Treatment of APOE4/4 miBrains with the TGF-β inhibitor SB431542 significantly reduced non-vascular αSMA immunoreactivity and led to a significant increase of pericyte vascular coverage to levels comparable to APOE3/3 controls (αSMA: 0.38 ± 0.091-fold change, corresponding to a 62% decrease relative to APOE4/4, p < 0.0001; pericyte coverage: 1.37 ± 0.072-fold increase relative to APOE4/4, p = 0.0249; **Figure 6B**), indicating a reversal of the PMT. Treatment of APOE4/4 miBrains with Galunisertib produced similar results (**Extended Data 7E**). Given the causal link we established between fibronectin secretion from myofibroblasts and perivascular amyloid accumulation, we next tested whether TGF-β inhibition could reduce both vascular fibrosis and amyloid burden. Consistent with this prediction, treatment of APOE4/4 miBrains with either SB431542 or Galunisertib significantly decreased fibronectin levels (SB431542: 0.69 ± 0.071-fold change, corresponding to a 31% decrease relative to APOE4/4, p = 0.0123; Galunisertib: 0.67 ± 0.030-fold change, corresponding to a 33% decrease relative to APOE4/4, p = 0.0007; **Figure 6C; Extended Data 7F**). Notably, these reductions in fibronectin were accompanied by significant reductions in amyloid deposition (D54D2 antibody), restoring levels to those seen in APOE3/3 control miBrains (SB431542: 0.61 ± 0.043-fold change, corresponding to a 39% decrease relative to APOE4/4, p = 0.0378; Galunisertib: 0.49 ± 0.034-fold change, corresponding to a 51% decrease relative to APOE4/4, p = 0.0124; **Figure 6C; Extended Data 7F**). Reductions in amyloid immunoreactivity following treatment with either drug were also observed with an additional amyloid antibody (12F4) (**Extended Data 7G**). To confirm that the observed findings were the result of inhibited TGF-β signaling as opposed to potential off-target effects, we stained for pSMAD2 and quantified the proportions of positive nuclei. This revealed a significant reduction in pSMAD2⁺ nuclei for both drugs, consistent with effective suppression of canonical TGF-β signaling (SB431542: 0.31 ± 0.077-fold change, corresponding to a 69% decrease relative to DMSO control, p = 0.0462; Galunisertib: 0.20 ± 0.066-fold change, corresponding to a 80% decrease relative to DMSO control, p = 0.0223; **Extended Data 7H)**.

Finally, we asked whether TGF-β inhibition could reverse the pericyte-to-myofibroblast transition, cerebrovascular fibrosis, and amyloid accumulation in vivo. To test this, we treated 12-month-old APOE4/4-TR mice with either a DMSO vehicle or the TGF-β inhibitor GW788388 for four weeks via daily intraperitoneal injections (**Figure 6D**). Following GW788388 treatment, we observed a significant reduction in pSMAD2+ nuclei, confirming suppression of canonical TGF-β signaling with this drug (0.67 ± 0.083-fold change, corresponding to a 33% decrease relative to DMSO vehicle, p = 0.0114; **Extended Data 7I**). Consistent with our miBrain results, TGF-β inhibition in aged APOE4/4-TR mice significantly increased pericyte coverage and reduced αSMA immunoreactivity relative to control (αSMA: 0.51 ± 0.11-fold change, corresponding to a 49% decrease relative to DMSO vehicle, p = 0.0067; pericyte coverage: 1.14 ± 0.034-fold increase relative to DMSO vehicle, p = 0.0326; **Figure 6E**), indicating a reversal of the pericyte-to-myofibroblast transition in vivo. Strikingly, this reversal was accompanied by a significant decrease in fibronectin deposition and amyloid-β immunoreactivity in drug-treated mice (fibronectin: 0.58 ± 0.039-fold change, corresponding to a 41% decrease relative to DMSO vehicle, p = 0.0175; amyloid-β: 0.81 ± 0.044-fold change, corresponding to a 19% decrease relative to DMSO vehicle, p = 0.0368; **Figure 6F**). Together, these findings demonstrate that APOE4 promotes a pericyte-to-myofibroblast transition through upregulation of TGF-β receptor signaling, driving vascular fibrosis and fibronectin-dependent amyloid accumulation, and that pharmacological inhibition of this pathway reverses these pathological phenotypes in vitro and in vivo (**Figure 6G**).L

## Discussion

Overall, the results presented here reveal that elevated TGF-β signaling drives a pericyte-to-myofibroblast transition in the APOE4 brain. This leads to a loss of pericyte coverage and fibrosis of the APOE4 vasculature, with fibronectin accumulation directly promoting perivascular amyloid deposition. Pharmacologically inhibiting TGF-β signaling in APOE4 in vitro and in vivo models reverses the pericyte-to-myofibroblast transition and attenuates cerebrovascular pathology. These results establish a causal link between TGF-β and cerebrovascular dysfunction in APOE4 carriers, which may influence AD progression in this large patient population. This study thus discovers a novel pathological mechanism in AD and reveals therapeutic opportunities for APOE4-driven cerebrovascular pathology. Importantly, targeting this mechanism may reduce the severe vascular side effects APOE4 carriers experience with anti-amyloid monoclonal antibody therapies.

AD has long been characterized by the presence of amyloid plaques and neurofibrillary tangles in the brain parenchyma^104^. However, cerebrovascular pathology is often one of the earliest manifestations of AD, sometimes occurring even before symptoms arise^27^. As part of disease progression, pericytes are lost along the vasculature leading to microvessel degeneration and the leakage of intravascular components into the brain parenchyma^48,105–108^. In addition to forming plaques, amyloid often accumulates along the vasculature in AD patients, leading to cerebral amyloid angiopathy^109^. CAA can occur in both large vessels and capillaries (Type 2 and Type 1 CAA, respectively), with APOE4 being particularly associated with capillary involvement^110^. Perivascular ECM accumulation is another feature of aging and AD, and several cerebrovascular ECM proteins, particularly fibronectin, are positively correlated with amyloid pathology^111–129^. Strikingly, TGF-β overexpression in mice leads to thickening of the cerebrovascular basement membrane that precedes amyloid accumulation and vascular dysfunction^60^. These findings suggest a link between TGF-β, perivascular fibrosis, and AD pathogenesis.

This study is the first to demonstrate that APOE4 directly increases TGF-β signaling along the cerebrovasculature, leading to vascular fibrosis and amyloid deposition via pericyte differentiation into myofibroblasts. Cerebrovascular degeneration is a well-characterized phenotype in APOE4 carriers that is linked to accelerated cognitive decline^28^. Independent of AD pathology, pericyte coverage is significantly reduced in APOE4 individuals and carriers of this allele are more susceptible to cerebral hemorrhages^28,101,130^. Additionally, perivascular amyloid accumulation is far more common in APOE4 carriers^101,102^. More recently, ECM dysregulation has also been implicated in APOE4-associated AD. APOE4 astrocytes and microglia upregulate matrisome pathways^10,131^ and solutes from the brain parenchyma, including amyloid, drain along the vasculature^132–135^. This suggests that perivascular accumulation of pro-aggregatory ECM proteins may drive impaired amyloid clearance^111,116^. Indeed, perivascular amyloid drainage is decreased in human APOE4-targeted replacement mice^136^. Strong evidence for a fibrotic phenotype in APOE4-mediated AD pathogenesis is a recent study that identified genetic factors protecting carriers of this allele from developing the disease^57^. They found that many protective variants are in ECM genes and that a particular loss-of-function *FN1* variant reduces the risk of developing AD by approximately 71%. Coinciding with these results, the brains of APOE4 carriers have increased perivascular fibronectin deposition compared to non-carriers^57,58^. To date, these APOE4-associated vascular pathologies – pericyte loss, amyloid deposition, and fibrosis – have all been investigated separately. However, this study uniquely identifies a TGF-β-dependent mechanism that links these phenotypes together into a single, targetable pathway. We show that inhibiting TGF-β can reverse cerebrovascular pathology in APOE4 which is, at least in part, due to reduced fibronectin accumulation. Therefore, the work presented here simultaneously explains a mechanism of AD risk and resilience and offers a novel therapeutic opportunity for APOE4 individuals.

While our data support a primarily cell-autonomous role for APOE4 in driving the pericyte-to-myofibroblast transition, several important questions remain. First, increased TGF-β ligand expression has been reported across multiple APOE4 cell types, including astrocytes and microglia, which raises the possibility that non–cell-autonomous signaling may contribute to mural cell reprogramming in APOE4 brains^137,138^. Defining the contributions of autocrine versus paracrine TGF-β signaling in vivo will be an important area for future study. Second, our findings that APOE expression is required for myofibroblast accumulation (**Extended Data 4D-E**) suggest that therapeutic strategies aimed at lowering APOE levels may represent an alternative approach to targeting downstream TGF-β signaling. Finally, APOE4 has been widely associated with lipid dysregulation across multiple CNS cell types, and cholesterol-lowering therapies have been shown to reduce myofibroblast accumulation and fibrosis in other organ systems^139–142^. Although we do not define the precise mechanism linking APOE4 to increased TGF-β signaling, these observations raise the possibility that altered lipid homeostasis may act upstream of, or in parallel with, TGF-β pathway activation in driving the pericyte-to-myofibroblast transition.

Overall, our study utilizes an orthogonal approach integrating single-nuclei transcriptomic and immunocytochemical analysis of the postmortem human brain with isogenic iPSC brain models and humanized APOE knock-in mice. From this rigorous approach, we discover for the first time a causal link between APOE4, TGF-β signaling, cerebrovascular fibrosis, and AD pathology, opening a path to new therapeutic opportunities in the treatment and prevention of AD in high-risk individuals.

Limitations of the Study:

This study has several limitations. The sample sizes for the miBrain snRNA-seq dataset (n = 2 per genotype) and iMC RNA-seq experiments (n = 3 per genotype) were modest, which may limit statistical power. In addition, the miBrain snRNA-seq dataset was previously generated in our laboratory using a model that includes neurons carrying the A53T mutation. However, our data indicate that the pericyte-to-myofibroblast transition occurs in a mural cell–autonomous manner, suggesting that the neuronal genotype is unlikely to drive the observed phenotype. Furthermore, pharmacological inhibition of TGF-β signaling in the miBrain model affects multiple cell types simultaneously, preventing definitive attribution of the rescue effects specifically to mural cells. Finally, although TGF-β inhibition mitigated APOE4-associated vascular phenotypes in our experimental systems, systemic inhibition of this pathway may have safety considerations given the broad physiological roles of TGF-β signaling.

## Supporting information

Supplementary Information Guide

## Resource availability

### 1. Lead Contact

Further information and requests for resources and reagents should be directed to the lead contact, Joel Blanchard.

### 2. Materials availability

This study did not generate new unique reagents.

### 3. Data and code availability

All datasets utilized in this study are from publicly available sources. The code used in this study is available at https://github.com/bschuldt99/Schuldt2026. Any additional information required to reanalyze the data reported in this paper is available from the lead contact upon request.

## Acknowledgements

We would like to thank Tatyana Kareva for technical and administrative support and the entire Blanchard Lab for scientific discourse. We acknowledge the use of BioRender for the generation of illustrations. This work has been funded in part with federal funds from NASA under contract #80ARC022CA004. This work was also supported by funding from the NIH/NIA (R01AG089533, UH3NS115064, U54AG090669), and the CureAlz Fund. BRS was supported by funding from the NIH (T32GM146636). ACP acknowledges funding from NIH Grants R01 AG063819; R01 AG064020; the Alzheimer’s Association; the Sanford J Grossman Charitable Trust; Brian and Tania Higgins Charitable Foundation; the Carolyn and Eugene Mercy Research Gift; the Robert J. and Claire Pasarow Foundation.

## Author Contributions

BRS performed computational analyses of the transcriptomic data. BRS, DHS, APA, LW, and AH performed the experiments and data analysis. The study was conceived and designed by BRS, JWB, DWMB, and ACP. BRS and JWB made the figures. BRS, JWB, ACP, LW, GG, GR, and AB wrote the paper. All authors critically reviewed the manuscript.

## Declaration of generative AI and AI-assisted technologies

During the preparation of this work, the authors used ChatGPT to assist with troubleshooting of errors that arose in R code. After using this tool, the authors reviewed and edited the code as needed and take full responsibility for the content of the publication.

## Methods

### Key Resources Table

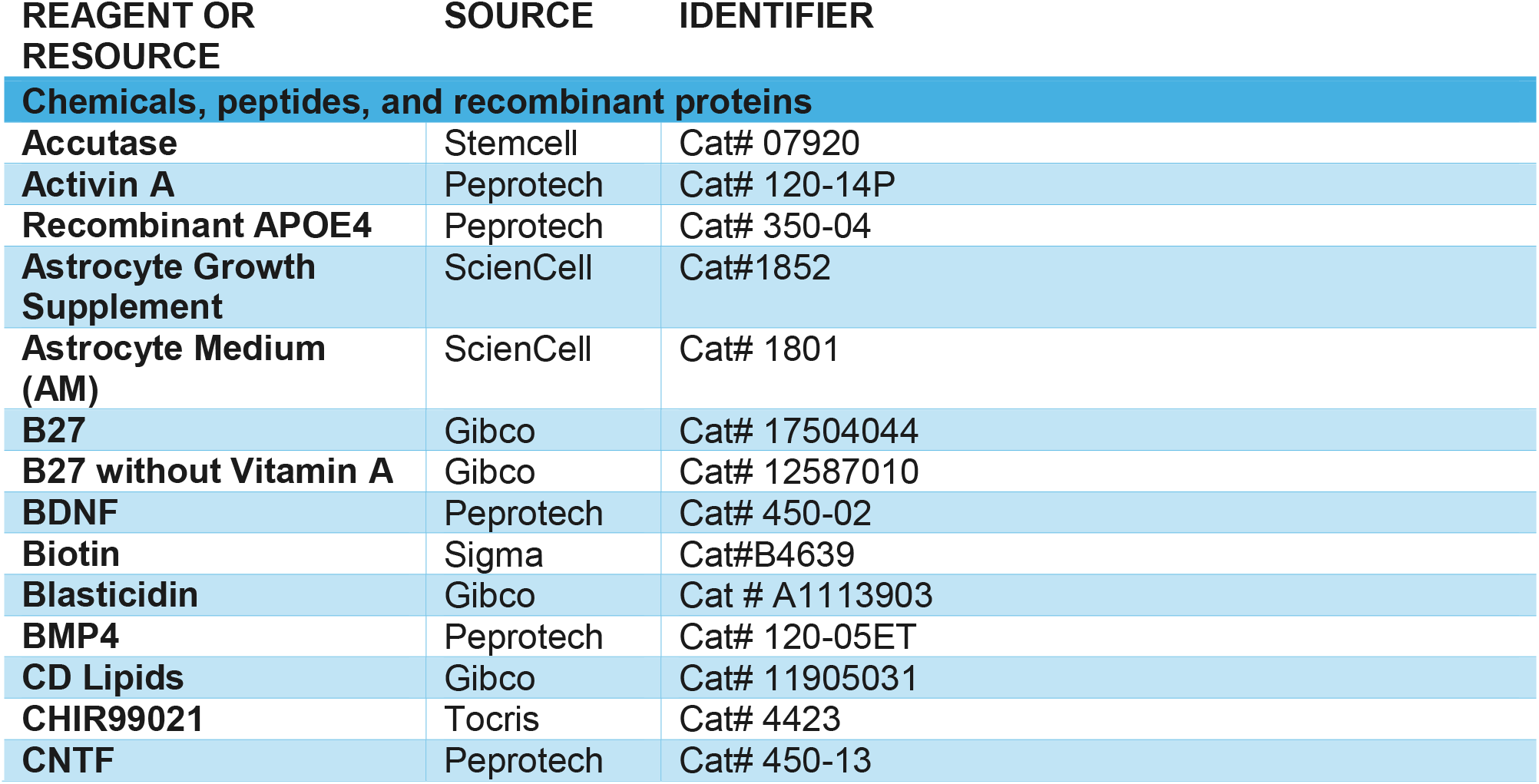
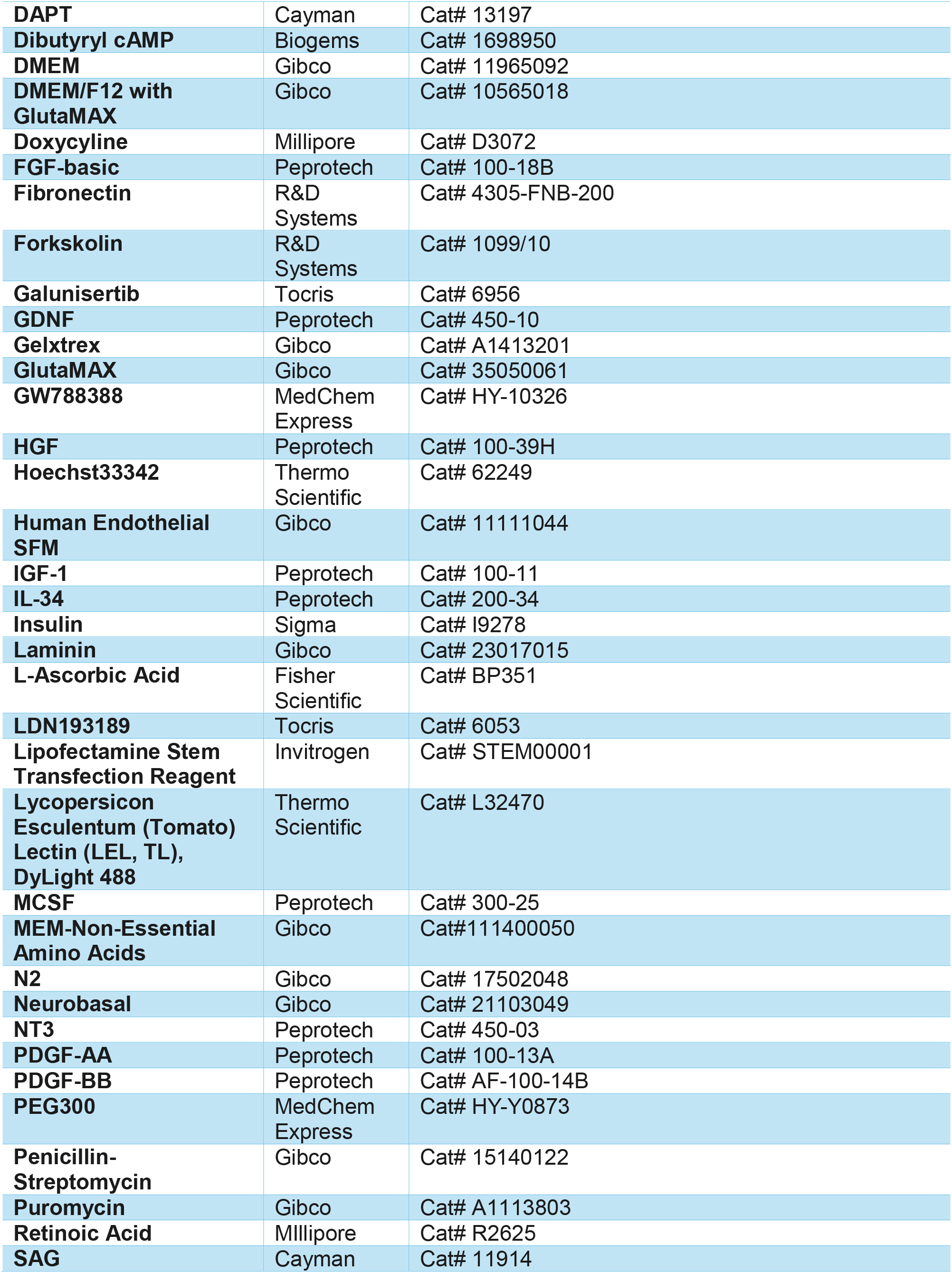
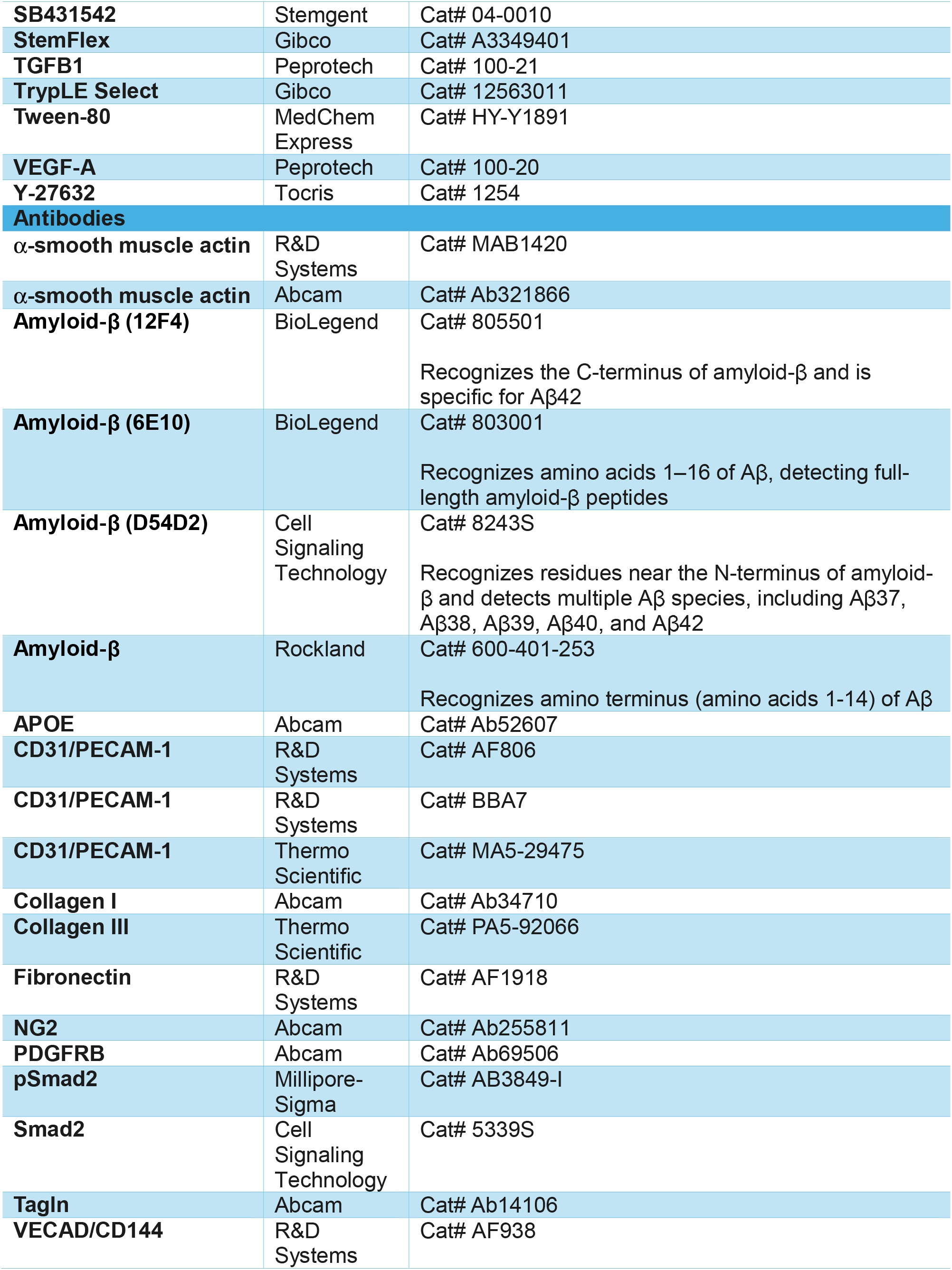
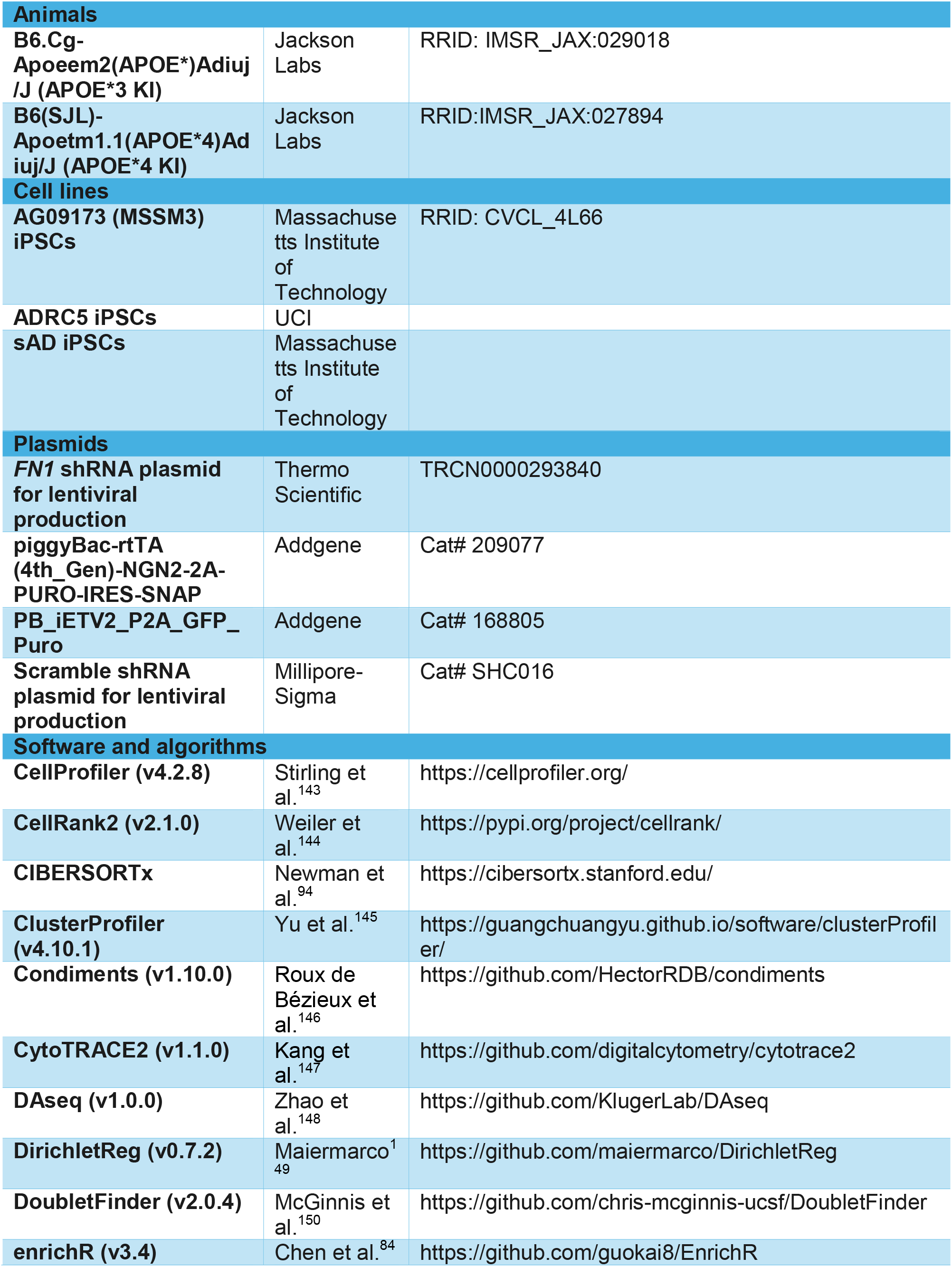
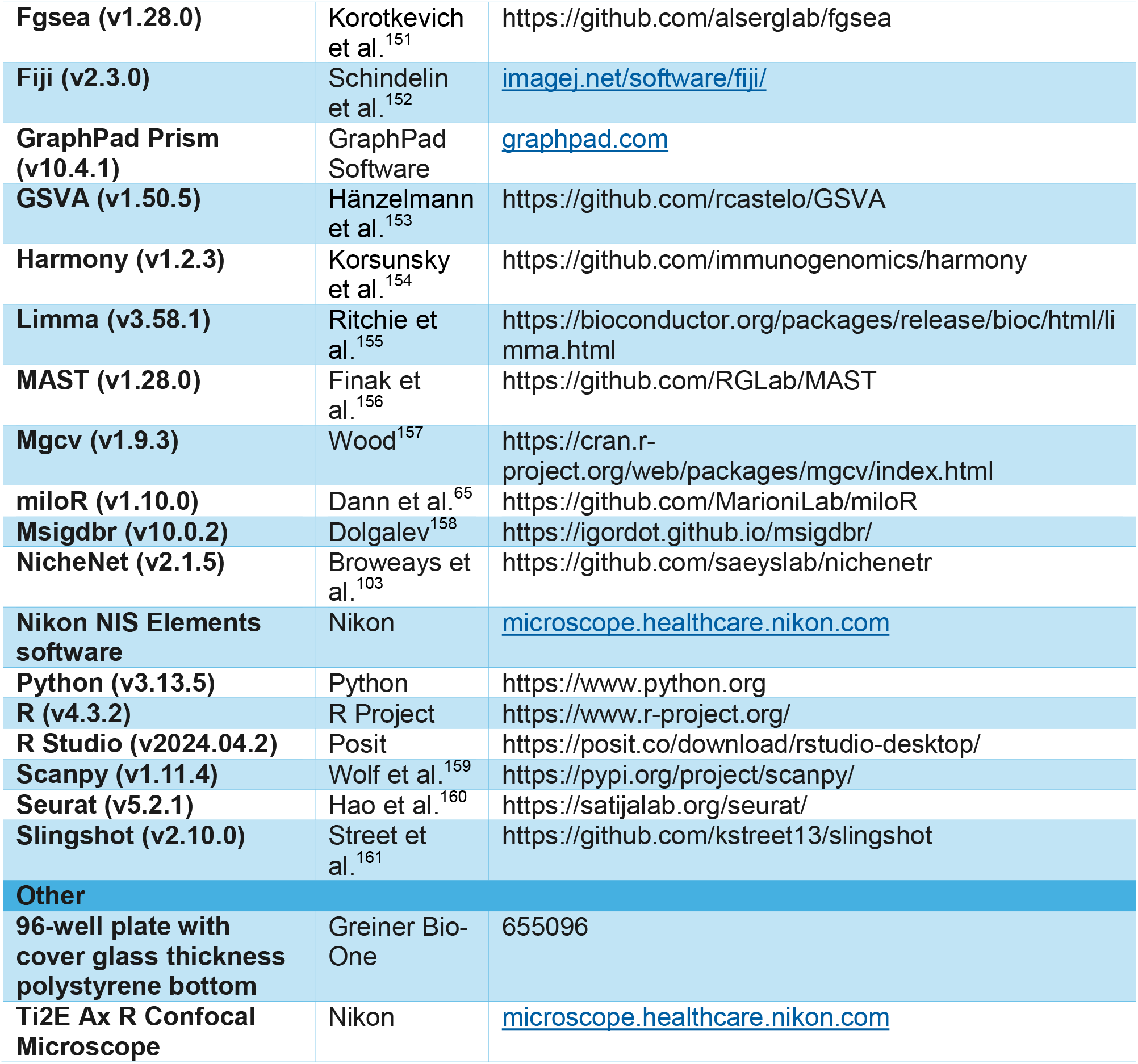

## EXPERIMENTAL MODEL AND STUDY PARTICIPANT DETAILS

### Cell lines

This study utilized iPSC lines from three different donors as described in the Key Resources Table.

### Mouse models

All experiments were performed in accordance with the Guide for the Care and Use of Laboratory Animals and were approved by the Institutional Animal Care and Use Committee at the Icahn School of Medicine at Mount Sinai (ISMMS). For all mouse experiments, C57BL/6J background mice with their endogenous murine *Apoe* replaced with human APOE3/3 or APOE4/4 were used. Aged, retired breeder APOE-TR mice were purchased from Jackson Laboratories (APOE3/3-TR strain number: 029018; APOE4/4-TR strain number: 027894). All mice used in APOE3/3- and APOE4/4-TR comparisons were females and ages ranged from 8-10 months. Mice used in the TGF-β inhibition experiment had equal numbers of males and females in both treatment and control groups (n = 6 per sex) and ages ranged from 12-13 months. Mice were housed in the ISMMS vivarium under standard conditions until experimentation.

## METHOD DETAILS

### Cerebrovascular atlas assembly and analysis

#### Processing and subject selection of individual snRNA-seq studies

Datasets from the following studies were downloaded and processed following the original methods as a guide. All analysis was performed in R with the Seurat package^160^ unless otherwise specified.

Yang et al., 2022:

Raw data from hippocampal samples were downloaded from GSE163577. To pass quality control, nuclei had to meet the following criteria: (1) more than 200 features, (2) less than 5000 features,and (3) less than 5% mitochondrial RNAs. Doublets were filtered out using DoubletFinder^150^. The standard Seurat pipeline of *NormalizeData(), FindVariableFeatures()* (nfeatures = 2000), and *ScaleData()* was performed. The top 30 principal components (PCs) were used for UMAP generation at a resolution of 0.2. Harmony was used for batch correction^154^. Clusters were annotated using the marker genes defined in the source publication. Clusters annotated as astrocytes, endothelial cells, and mural cells were then extracted for generation of the harmonized cerebrovascular atlas.

Sun et al., 2023:

Processed and clustered data was downloaded from the source publication. Only individuals with APOE3/3, APOE3/4, and APOE4/4 genotypes were included. Nuclei labeled as endothelial cells, smooth muscle cells, and pericytes were extracted for construction of the cerebrovascular atlas.

Haney et al., 2024:

Raw data of AD samples was downloaded from GSE254205. Nuclei that passed the following quality control parameters were included for further analysis: (1) more than 500 features, (2) more than 1000 reads, (3) less than 10% mitochondrial reads, and (4) less than 10% ribosomal reads. Doublets were removed using DoubletFinder. The standard Seurat pipeline of *NormalizeData(), FindVariableFeatures()* (nfeatures = 2500), and *ScaleData()* was performed. The top 20 principal components (PCs) were used for UMAP generation at a resolution of 0.2. Harmony was used for batch correction. Clusters were annotated using the marker genes defined in the source publication. Clusters annotated as astrocytes and vascular cells were extracted.

After extracting the cerebrovascular cell types from each dataset, propensity matching for APOE4 carriers and non-carriers was performed within each dataset to minimize confounding variables and equalize the number of individuals with each genotype. Specifically, the R package MatchIt^162^ was employed to match individuals using the following parameters:

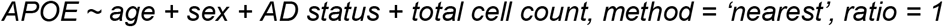

#### Cerebrovascular atlas integration and clustering

After propensity matching, data from the selected individuals were merged. The standard Seurat pipeline of *NormalizeData(), FindVariableFeatures()* (nfeatures = 2000), and *ScaleData()* was performed. Harmony was used for integration of the datasets with the following parameters:

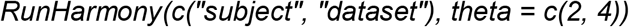

The top 15 principal components (PCs) were used for UMAP generation at a resolution of 0.5. Clusters with high expression of markers of two or more cell types were excluded from further analysis. After excluding a cluster, the Seurat pipeline and harmony integration was repeated. Ultimately, three rounds of clustering were performed and the final integrated UMAP was generated at a resolution of 0.2. Clusters were manually annotated using canonical marker genes for astrocytes, endothelial cells, pericytes, and smooth muscle cells (SMCs).

For mural cell subclustering, pericytes and smooth muscle cell clusters were extracted. *FindNeighbors(), FindClusters(),* and *RunUMAP()* were re-run with the top 15 PCs at a resolution of 0.2. Subclusters were annotated using previously described marker genes^62,63^.

#### Differential abundance analysis

miloR:

The miloR package was implemented in R^65^. For the entire cerebrovascular atlas, a k-nearest-neighbor (KNN) graph was generated using the *buildGraph()* function with parameters k = 50 and d = 15 Neighborhoods were generated with the *makeNhoods()* function with parameters prop = 0.1, k = 50, and d = 15 followed by *calcNhoodDistance()* with d = 15. Differential abundance between APOE4 carriers and non-carriers was assessed using the *testNhoods()* function with default parameters and the following design:

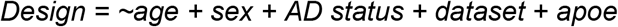

For miloR differential abundance analysis of the mural cell subclusters, the same pipeline was used as above. For all above functions, Harmony embeddings were used as the dimensionality reduction input.

Dirichlet regression on all vascular cell types:

For each individual, vascular cell types were aggregated and the number of nuclei assigned to each cell type was counted. The proportion of each cell type was then calculated relative to the total number of vascular cells within that subject. The resulting subject-level composition matrix (cell-type proportions that sum to one for each individual) was modeled using the DirichletReg package in R^149^. Cell-type proportions were used as the Dirichlet response variable and modeled as a function of APOE genotype, Alzheimer’s disease diagnosis, sex, age (modeled using a natural spline with 3 degrees of freedom), and dataset of origin. To account for differences in sampling depth across donors, models were weighted by the total number of cells contributed by each subject.

Dirichlet regression on mural cell subclusters:

For each individual, nuclei belonging to mural subclusters were counted and converted to proportions by dividing each subcluster count by the total number of mural cells for that subject. The resulting subject-by–mural subcluster proportion matrix (where proportions across mural subclusters sum to one per subject) was analyzed using Dirichlet regression with the DirichletReg package in R^149^. Mural subcluster proportions were used as the multivariate Dirichlet response variable and modeled as a function of APOE genotype, Alzheimer’s disease diagnosis, sex, standardized age (z-scored), and dataset of origin. To account for differences in the number of mural cells profiled per donor, models were weighted by the total number of mural cells contributed by each subject.

Dirichlet regression on mural cell states:

Mural cells were classified into three transcriptional states based on expression of the canonical pericyte marker *CSPG4* and the smooth muscle marker *ACTA2*. *CSPG4* and *ACTA2* expression values were extracted from the Seurat object metadata, and expression cutoffs were defined using the 1st percentile of non-zero expression for each gene across cells. Cells were then assigned to one of three mutually exclusive states: *CSPG4_high/ACTA2_low*, *CSPG4_high/ACTA2_high*, or *CSPG4_low/ACTA2_high*. Cells not meeting any of these criteria were excluded from downstream compositional analyses. For each individual, we aggregated cells by state to obtain subject-level counts and proportions, with proportions calculated relative to the total number of classified state-assigned mural cells per subject. Subject-level state compositions (three proportions summing to one) were modeled using Dirichlet regression^149^, with state composition as the multivariate response and APOE genotype, AD diagnosis, and an APOE×AD interaction as primary predictors, adjusting for standardized age, sex, and dataset of origin. To account for unequal sampling depth across individuals, models were weighted by the total number of classified state-assigned mural cells contributed by each subject.

DASeq:

The DAseq package was implemented in R^148^. To compute APOE4 carrier vs.non-carrier differential abundance in mural cell subclusters, Harmony embeddings were used as input to DAseq along with APOE genotype labels for each subject. A range of neighborhood sizes *(k.vector = seq(50, 500, 50))* was used to compute differential abundance (DA). Cells with DA scores above the 90^th^ percentile or below the 10^th^ percentile were classified as differentially abundant and visualized on UMAP coordinates. DA regions were identified using the *getDAregion()* function at a resolution of 0.5.

Binomial generalized linear model to estimate subject-level myofibroblast proportions:

For myofibroblast proportions, the number of cells in the SMC_2 DAseq region defined as myofibroblasts (**Figure 1H**) were counted for each individual. Myofibroblast proportion per individual was defined as the number of myofibroblast cells divided by the total number of SMC_2 cells. We modeled these subject-level proportions using a binomial generalized linear model (GLM) with a logit link in R (glm, family = binomial). Counts were represented in the response as a two-column matrix of successes and failures. The primary model tested whether the effect of APOE genotype differed by sex or AD diagnosis by including interaction terms for APOE×sex and APOE×AD, while adjusting for age and dataset of origin:

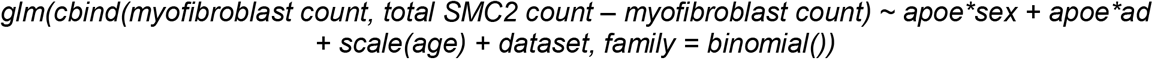

Age was standardized prior to modeling (z-scored) so that effect sizes corresponded to a change per standard-deviation increase in age. Model diagnostics were assessed with the DHARMA package^163^. Model coefficients for each for the variables were extracted and expressed as odds ratios with 95% confidence intervals. To report interpretable effect sizes, estimated marginal means were computed with the emmeans package and contrasts were performed on the logit scale, then exponentiated to obtain odds ratios (ORs) with 95% confidence intervals. In addition to reporting marginal main effects averaged over covariates (e.g., APOE4 vs APOE3), we quantified APOE-stratified contrasts within sex and within AD strata using Tukey-adjusted pairwise comparisons from emmeans for the APOE×sex and APOE×AD interaction terms.

Binomial GLM sensitivity analysis:

To assess robustness to potential dataset-specific effects, we performed a leave-one-dataset-out sensitivity analysis by refitting the model after excluding each dataset in turn (e.g., Sun+Yang only, Haney+Yang only, Haney+Sun only). Because the dataset variable is undefined once a dataset is removed and sample sizes are reduced, a simplified binomial GLM was used for the sensitivity analysis that omitted the dataset term and interaction terms while retaining the primary covariates (APOE genotype, sex, AD diagnosis, and standardized age). Individuals contributing fewer than 5 SMC_2 cells were excluded to improve model stability. For each dataset combination, the APOE4 versus APOE3 effect was summarized as an odds ratio with 95% confidence intervals derived from the fitted model.

#### Differential gene expression (DGE) analysis

Single-cell DGE analysis of mural cell subclusters:

Single-cell differential gene expression was performed with MAST^156^ by using the Seurat *FindMarkers()* function. Pairwise comparisons between APOE4 carriers and non-carriers were performed for each mural cell subcluster. Age, sex, AD status, and dataset were included as covariates. A differentially expressed gene was defined as |logFC| > 0.25 and an adjusted p-value < 0.05.

Pseudobulk DGE analysis of SMC_2 cluster:

For the mural cell subclusters, gene expression counts were aggregated across cells for each subject using the *AggregateExpression()* function in Seurat, grouping by subject, APOE genotype, and mural cell type. The data was filtered to only include SMC_2 cells from individuals contributing at least 5 cells. A voom-limma^155^ pipeline was then employed to model differential gene expression between APOE4 carriers and non-carriers. The following linear model was fit for each gene:

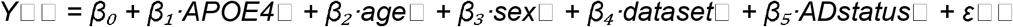

In this model, *Y*□□ represents the voom-transformed expression value of gene *i* in subject *j*; *APOE4*□ is an indicator variable for APOE4 genotype (with APOE3 as the reference); *age*□, *sex*□, *dataset*□, and *ADstatus*□ represent the subject-level covariates for age, sex, dataset, and AD status, respectively; β₀ is the model intercept; β₁*–*β₅ are the estimated coefficients for each predictor; and ε□□ is the residual error term. For each gene, statistics were calculated using the *eBayes()* function.

Myofibroblast marker genes:

DGE was performed with MAST by using the Seurat *FindMarkers()* function. Within the SMC_2 subcluster, a pairwise comparison was performed between the cells in the APOE4-enriched DAseq region (**Figure 1H**) and the remaining SMC_2 cells. A myofibroblast marker gene was defined as logFC > 0.25 and an adjusted p-value < 0.05.

#### Pathway analysis

Gene set enrichment analysis (GSEA) was performed using the fgsea R package^151^. The REACTOME database was input via the msigdbr R package^158^. Following APOE4 carrier vs. non-carrier differential gene expression analysis via MAST or voom-limma (see differential gene expression analysis methods section), genes were ranked according to the following formula:

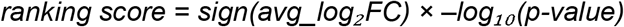

Genes were then sorted in decreasing order and input into the fgsea package. Enrichment scores were calculated using a running-sum statistic across the ranked gene list. The normalized enrichment score (NES) represents the enrichment score scaled to a null distribution generated by gene-set permutations, correcting for gene-set size and enabling comparison across pathways. The enrichment score, normalized enrichment score, p-values, and FDR-adjusted p-values were computed for each pathway. Significant pathways were defined as an adjusted p-value < 0.05. Redundant pathways were merged to enhance readability.

For Gene Ontology (GO) enrichment analysis, differentially expressed genes were input into the ClusterProfiler R package^145^ using the enrichGO() function with *ont = “ALL”.* Redundant pathways were merged with the *simplify()* function. Significant pathways were defined as an adjusted p-value < 0.05.

#### Unbiased cell type annotation

Cell type annotation was performed with the CellMarker_Augmented_2021 database via the enrichR package^84,164^. APOE4-enriched genes from the pseudobulk SMC_2 DGE analysis (logFC > 0.25, p < 0.05, see DGE analysis methods) or the myofibroblast marker genes (see DGE analysis methods) were input into enrichR with default parameters. Only cell types with a defined tissue of origin and at least 10 marker genes and 3 hits were included. Redundant cell types were merged to enhance readability.

#### Seurat module scores

Cellular contraction, extracellular matrix (ECM), and co-expression scores in mural cells: Module scores were calculated for each cell using the *AddModuleScore()* function in Seurat. The following genes were used for each module:

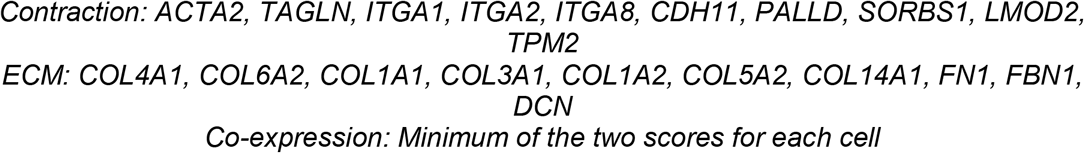

For comparison of module scores in APOE4 carriers vs. non-carriers, the data was filtered to only include SMC_2 cells from individuals contributing at least 5 cells. The module scores were then averaged across all cells per individual for subject-level comparisons between APOE genotypes.

TGF-β signaling in mural cells:

Mural cell subclusters were extracted and a module score was calculated for each cell using the *AddModuleScore()* function in Seurat. The following genes were used for the module:

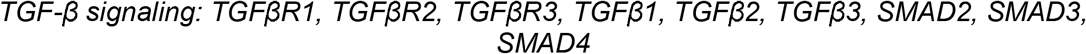

For comparison of TGF-β gene expression in APOE4 carriers vs. non-carriers, the module score was then averaged across all cells per individual for subject-level comparisons between APOE genotypes.

#### Myofibroblast gene activity scores

Gene set variation analysis was performed using the GSVA R package^153^. The data was filtered to only include SMC_2 cells from individuals contributing at least 5 cells. Gene expression counts were aggregated across cells for each subject using the *AggregateExpression()* function in Seurat, grouping by subject and APOE genotype. Myofibroblast marker genes were downloaded from a previous publication (cluster 11)^87^ and GSVA scores were assigned to each subject using the top 10 highly expressed myofibroblast marker genes in our dataset.

#### Pseudotime analysis

Lineage assignment:

Lineages were generated using the slingshot R package^161^. The UMAP of mural cell subclusters was input into slingshot with the Pericyte_2 cluster assigned as the start cluster.

Lineage bias:

For the lineages inferred by slingshot, each cell was assigned a pseudotime value along each lineage, as well as a corresponding curve weight. Curve weights represent the probability that a given cell belongs to a particular lineage when multiple branching trajectories are present. In this framework, cells located near a branch point may have fractional weights across lineages, whereas cells committed to a single trajectory have weights approaching 1 for that lineage and 0 for others.

After weight calculation, the R package condiments^146^ was then used to test for differential lineage assignment between APOE genotypes. Specifically, the *differentationTest()* function was performed for this anlaysis.

Generalized additive models (GAMs):

Three sets of module scores were assigned to mural cells using the *AddModuleScore()* function in Seurat with the following genes:

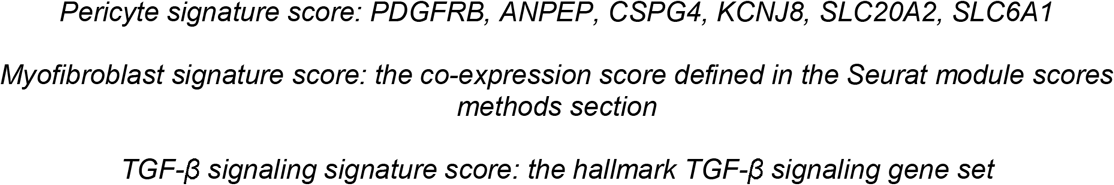

After assigning scores, separate GAMs with slingshot-derived lineage weights were fit for each module score with a smoothing parameter of *k = 7* using the mgcv R package^157^. To compare scores between lineages, the differences in areas under the curve (Δ-AUC: pericyte-to-myofibroblast – pericyte-to-pericyte lineage) was calculated for each GAM. Permutation testing (n = 1000) was then used to generate a null distribution of Δ-AUC values. A two-sided p-value was calculated as the proportion of permuted Δ-AUC values exceeding the observed Δ-AUC in magnitude.

#### NicheNet

The NicheNet R package was utilized for this analysis^103^. A sender-focused approach was performed. The DAseq APOE4-enriched myofibroblast region (**Figure 1H**) was assigned as the receiver cell type. The astrocyte, endothelial, pericyte_1, pericyte_2, SMC_1, SMC_2, and myofibroblast clusters were assigned as the sender cell types. Genes expressed in at least 10% of receivers or senders were used in the analysis, and the differential genes input into NicheNet for ligand prediction were the myofibroblast marker genes defined in the DGE analysis methods section.

#### Association of mural cell *FN1* expression with AD phenotypes

Mural cells from the Haney (2024) dataset^11^ were extracted from the cerebrovascular atlas and *FN1* expression was averaged per individual. Mural cell *FN1* expression was then correlated with age of AD diagnosis. A global cerebral pathology score was assigned to each individual by averaging the following metadata columns reported in the original dataset: *sum_lb_density, PlaqueTotal, TangleTotal, infarct_cerebral_total_volume.* Scores were then dichotomized into high (top 25%) and low (bottom 75%) pathology groups to compare mural cell *FN1* expression between them.

#### *FN1* gene expression in fibroblasts versus myofibroblasts

The original Yang and Sun snRNA-seq datasets each contained annotated fibroblast clusters. To enable direct comparison, we returned to these datasets and extracted fibroblast cells from the same subjects represented in our integrated cerebrovascular atlas, while myofibroblasts were defined as previously described and mapped back to the original objects by cell barcode. The datasets were harmonized and merged, and FN1 expression was pseudobulked by averaging log-normalized expression per subject within each cell type. Only subjects containing both fibroblasts and myofibroblasts were retained for analysis.

### miBrain single-nuclei RNA-seq analysis

#### Dataset processing and clustering

Raw data was downloaded from a previous dataset generated in our laboratory. To pass quality control, nuclei had to meet the following criteria: (1) more than 500 features, (2) less than 6500 features, and (3) less than 10% mitochondrial RNAs. Doublets were filtered out using DoubletFinder^150^. The standard Seurat pipeline of *NormalizeData(), FindVariableFeatures()* (nfeatures = 2000), and *ScaleData()* was performed. The top 15 principal components (PCs) were used for UMAP generation at a resolution of 0.2. Harmony was used for batch correction^154^. Clusters were annotated using canonical cell type marker genes. Clusters with high expression of markers of two or more cell types were excluded from further analysis. After excluding a cluster, the Seurat pipeline and harmony integration was repeated. Clusters annotated as astrocytes, endothelial cells, and mural cells were then extracted for downstream vascular cell analysis.

For mural cell subclustering, *FindNeighbors(), FindClusters(),* and *RunUMAP()* were re-run with the top 15 PCs at a resolution of 0.05. Subclusters were annotated using previously described marker genes^62,63^.

#### Differential gene expression (DGE) analysis

Single-cell DGE analysis of mural cell markers:

Single-cell differential gene expression was performed with MAST^156^ by using the Seurat *FindMarkers()* function. A pairwise comparison between APOE4/4-enriched mural cluster 2 and APOE3/3-enriched mural cluster 1 was performed. A differentially expressed gene was defined as |logFC| > 0.75 and an adjusted p-value < 0.05.

Pseudobulk DGE analysis of mural cells:

Mural cell gene expression counts were aggregated across cells for each miBrain using the *AggregateExpression()* function in Seurat, grouping by sample and APOE genotype. A voom-limma^155^ pipeline was then employed to model differential gene expression between APOE4/4 and APOE3/3 miBrains. The following linear model was fit for each gene:

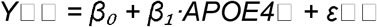

In this model, *Y*□□ represents the voom-transformed expression value of gene *i* in subject *j*; *APOE4*□ is an indicator variable for APOE4 genotype (with APOE3 as the reference); β₀ is the model intercept; β₁ is the estimated coefficients; and ε□□ is the residual error term. For each gene, statistics were calculated using the *eBayes()* function.

#### Pathway analysis

See section under ‘Cerebrovascular atlas assembly and analysis.’

#### Seurat module scores

Pericyte and myofibroblast scores in mural cells:

Module scores were calculated for each cell using the *AddModuleScore()* function in Seurat. The following genes were used for each module:

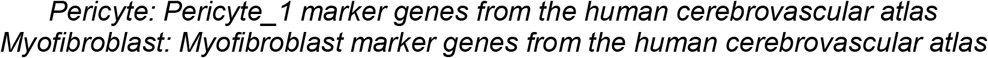

TGF-β signaling in mural cells:

A a module score was calculated for each cell using the *AddModuleScore()* function in Seurat. The following genes were used for the module:

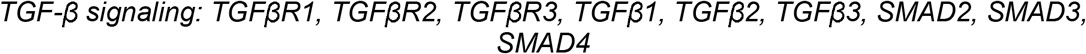

For comparison of TGF-β gene expression in APOE4/4 vs. APOE3/3 miBrains, the module score was then averaged across all cells per miBrain for sample-level comparisons between APOE genotypes.

#### CytoTRACE2 and CellRank2

To estimate differentiation potential in miBrain mural cells, CytoTRACE2^147^ was implemented in R using default parameters. For trajectory inference, the miBrain mural cell object was converted to an AnnData object for downstream analysis in Python. A k-nearest neighbor graph was generated in Scanpy^159^ from the Harmony embedding (*X_harmony*) using n*_neighbors* = 30 and the Gaussian kernel method (*method = “gauss“*). CytoTRACE2 scores were then inverted to represent pseudotime progression and incorporated into CellRank2^144^ using the *PseudotimeKernel*. Transition probabilities were computed using the soft thresholding scheme with nu = 0.3 and visualized by projection onto the UMAP embedding. The TGF-β signaling signature score for correlation analysis with cytoTRACE was assigned using the hallmark TGF-β signaling gene set as described in the human methods section ‘Pseudotime Analysis.’

### Mouse single-cell RNA-seq analysis

#### Dataset processing and clustering

Raw data was downloaded from a previous study^165^. To pass quality control, nuclei had to meet the following criteria: (1) more than 200 features, (2) less than 4000 features, and (3) less than 25% mitochondrial RNAs. Doublets were filtered out using DoubletFinder^150^. The standard Seurat pipeline of *NormalizeData(), FindVariableFeatures()* (nfeatures = 2000), and *ScaleData()* was performed. The top 20 principal components (PCs) were used for UMAP generation at a resolution of 0.2. Harmony was used for batch correction^154^. Clusters were annotated using canonical cell type marker genes. Clusters with high expression of markers of two or more cell types were excluded from further analysis. After excluding a cluster, the Seurat pipeline and harmony integration was repeated. Clusters annotated as astrocytes, endothelial cells, and mural cells were then extracted for downstream vascular cell analysis.

For mural cell subclustering, *FindNeighbors(), FindClusters(),* and *RunUMAP()* were re-run with the top 15 PCs at a resolution of 0.2. Subclusters were annotated using previously described marker genes^62,63^.

#### Differential abundance analysis

DAseq:

The DAseq package was implemented in R^148^. Differential abundance analysis was performed for aged iE4/Cre+ (>250 days old) versus all others. Harmony embeddings were used as input to DAseq along with APOE genotype labels for each mouse. A range of neighborhood sizes *(k.vector = seq(50, 250, 50))* was used to compute differential abundance (DA). DA regions were identified using the *getDAregion()* function at a resolution of 0.05.

Temporal analysis of pericyte and myofibroblast abundance:

Cells in DA region 2 were labeled as myofibroblasts and cells in DA region 5 were labeled as pericytes. For those DA regions, the following binomial logistic mixed-effects model was fit separately:

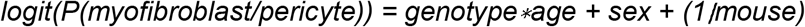

#### Differential gene expression (DGE) analysis

DGE was performed with MAST by using the Seurat *FindMarkers()* function. A pairwise comparison was performed between the cells in the aged iE4/Cre+-enriched DAseq region (DA region 2 in the source code, subsequently labeled myofibroblast) and all other mural cells. A myofibroblast marker gene was defined as logFC > 0.75 and an adjusted p-value < 0.05.

#### Pathway analysis

For Gene Ontology (GO) enrichment analysis, differentially expressed genes were input into the ClusterProfiler R package^145^ using the enrichGO() function with *ont = “ALL”.* Redundant pathways were merged with the *simplify()* function. Significant pathways were defined as an adjusted p-value < 0.05.

#### Unbiased cell type annotation

See section under ‘Cerebrovascular atlas assembly and analysis.’

### Age analysis of APOE-TR mice

The mouse bulk RNA-seq data was downloaded from a previous study^95^. For CIBERSORTx^94^ deconvolution, the web-based application was used. Cells in DAseq region 2 and DAseq region 5 from the ‘Mouse single-cell RNA-seq analysis’ section were used as the reference. A signature matrix was generated using standard parameters, with a sampling ratio of 0.99 and inclusion of all genes. Absolute cell type scores were then estimated from the bulk RNA-seq gene expression data using default CIBERSORTx settings.

For each absolute score (pericyte or myofibroblast), age and genotype differences were tested using a fixed-effects ANOVA of the form:

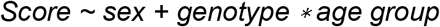

Post hoc comparisons between APOE3/3- vs APOE4/4-TR within each age group were performed using estimated marginal means (emmeans) with Tukey adjustment for multiple testing.

### Myofibroblast gene activity in APOE-TR mice

Gene set variation analysis was performed using the GSVA R package^153^. The processed and normalized 4-month-old mouse bulk RNA-seq data was downloaded from Synapse (syn26561824)^89^. GSVA scores were calculated for each sample using the myofibroblast marker genes from our cerebrovascular atlas (see myofibroblast marker genes in the DGE analysis methods section, all genes with adjusted p-value < 0.05 were used). The human genes were converted to their mouse orthologs prior to GSVA input.

### RNA-seq analysis of iPSC-derived mural cells

FPKM values were analyzed from a previous study^50^. After log-transforming the FPKM values, differential expression analysis between APOE3/3 and APOE4/4 iPSC-derived mural cells (iMCs) was performed using the standard limma pipeline. Array weights were estimated using the *arrayWeights()* function and statistics were calculated for each gene using the *eBayes()* function with *trend = TRUE* and *robust = TRUE.* A differentially expressed gene was defined as |logFC| > 0.25 and an adjusted p-value < 0.05. Differentially expressed genes were input into ClusterProfiler for GO pathway analysis. Significant pathways were defined as an adjusted p-value < 0.05.

For CIBERSORTx^94^ deconvolution, the web-based application was used. Mural cell subclusters from the cerebrovascular atlas were used as the reference. Specifically, 2,000 mural cells were randomly downsampled from the atlas, and their gene counts were extracted. A signature matrix was generated using standard parameters, with a sampling ratio of 0.5 and inclusion of genes expressed in at least 10% of cells. Relative cell type proportions were then estimated from the bulk RNA-seq gene expression data using default CIBERSORTx settings. Pericyte (Pericyte 1 and Pericyte 2) and SMC (SMC 1 and SMC 2) subcluster relative proportions were added to obtain overall pericyte and SMC proportions. For downstream analysis, cell type proportions in APOE4/4 samples were expressed as fold changes relative to the mean proportions in APOE3/3 samples.

### Human iPSC cultures

Pre-coated Geltrex ™ Matrix plates were seeded with iPSCs. iPSC colonies were then grown in StemFlex ™ medium until 60-70% confluency. Subsequently, maintenance iPSCs were passaged using 0.5 mM EDTA to gently lift colonies. iPSCs for differentiation were harvested with Accutase ™ cell detachment solution for 5-10 minutes at 37C and differentiated as single cells according to the specified differentiated protocol.

### Differentiation of iPSCs

#### Differentiation of human iPSCs into neurons

Protocol for neuron differentiation was adapted from Zhang et al.^166^ and Lam et al.^167^. In brief, Lipofectamine™ Stem Transfection Reagent was used to transfect iPSCs with PiggyBac plasmids to confer doxycycline-inducible expression of the Neurogenin-2 gene (*NGN2*, Addgene Plasmid #209077). In short, dissociated iPSCs were seeded on day zero at ∼104,000 cells/cm^2^ onto plates coated with Geltrex™. The seeded cells then received StemFlex™ media containing 10µM Y27632 and 5µg/mL doxycycline supplements. On day 1, culture medium was replaced with Neurobasal N2B27 medium (Neurobasal, 1x B-27, 1x N-2, 1x MEM-NEAA, 1x GlutaMax, 1% penicillin-streptomycin) containing 10µM SB431542, 100nM LDN, 5µg/mL doxycycline, and 5µg/mL puromycin supplements. On days 3-6, daily medium changes were performed with Neurobasal N2B27 media containing 1µg/mL puromycin and 5µg/mL doxycycline supplements. On day 6, cells were treated with 0.5 µM Ara-C. On day 7, cells were lifted with accutase for integration into miBrains.

#### Differentiation of human iPSCs into astrocytes

Previously published iPSC-derived NPC (Chambers et al.)^168^ and astrocyte (TCW et al.)^10^ differentiation protocols were followed to generate astrocytes. In summary, dissociated iPSCs were seeded onto Geltrex ™ - coated plates at a cell density of 100,000 cells/cm^2^ in pre-warmed StemFlex ™ media containing 10µM Y27632 supplement. Cells received StemFlex ™ every other day until cells reached 95% confluence. Once confluent, cells received NPC medium (1:1 DMEM/F12: Neurobasal Medium, 1x N-2 Supplement, 1x B-27 Serum-Free supplement, 1x GlutaMAX Supplement, 1x MEM-NEAA, 1% penicillin-streptomycin) containing 10µM SB43152 and 100nM LDN193189 supplements (day 0). From days 1 to 9, cells received daily media changes with the same media as day 0. On day 10, cells were split with Accutase and seeded onto plates coated with Geltrex ™ and were grown in NPC media supplemented with 10 µM Y27632 and 20ng/mL bFGF. From days 11 through 13, cells received NPC media supplemented 20 ng/mL bFGF. On day 14, cells were split with Accutase and replated onto Geltrex ™ -coated plates and received NPC media supplemented with 10µM Y27632 and 20ng/mL bFGF. From day 15 onward, cells received fresh Astrocyte Medium (AM, ScienCell) every 2-3 days and were passaged using Accutase upon reaching 90% confluence. From this point on, NPCs were differentiated into astrocytes over a 30-day time course. NPCs and differentiated astrocytes were then cryopreserved in freezing medium, which consisted of 90% knockout serum replacement (KSR) and 10% dimethyl sulfoxide (DMSO).

#### Differentiation of human iPSC into brain microvascular endothelial cells

Brain endothelial cells were differentiated according to adapted protocols outlined in Blanchard et al.^50^, Qian et al.^169^, and Wang et al.^170^. In summary, doxycycline-inducible expression of the ETS variant transcription factor 2 (*ETV2*, Addgene Plasmid #168805) was conferred in iPSCs by transfecting iPSCs with a PiggyBac plasmid. Inducible *ETV2*-iPSCs were cultured until 60-70% confluence before they were dissociated with Accutase and seeded at 20,800 cells/cm^2^ onto plates coated in Geltrex ™ that contained StemFlex™ supplemented with 10µM Y27632 on day 0. On day 1, culture medium was replaced with DeSR1 medium (DMEM/F12 with GlutaMAX, 1x MEM-NEAA, 1x penicillin-streptomycin) containing 10 ng/mL BMP4, 6µM CHIR99021, and 5µg/mL doxycycline supplements. On days 5 and 7, cell media was changed and replaced with hECSR medium (Human Endothelial Serum-free Medium, Gibco; 1x MEM-NEAA; 1 x B-27, 1% penicillin-streptomycin) that was supplemented with 50 ng/mL VEGF-A, 2µM Forskolin, and 5µg/mL doxycycline. On day 8, cells were lifted with Accutase and re-plated onto fresh plates coated with Geltrex ™. Cells received hECSR media that was supplemented with 50 ng/mL VEGF-A and 5 µg/mL doxycycline. In the week following day 8, the supplemented day 8 medium was given to cells every 2-3 days to maintain cells until they were ready for tissue assembly in miBrains.

#### Differentiation of human iPSCs into mural cells

Mural cell differentiation protocol was based on published protocol from Patsch et al.^171^. Dissociated iPSCs were seeded at 37,000 to 40,000 cells/cm^2^ onto plates coated with Geltrex ™ and grown in StemFlex ™ containing 10µM Y27632 supplement on day 0. On day 1, StemFlex ™ medium was replaced with N2B27 medium (1:1 DMEM/F12, Neurobasal media, 1x B-27, 1x N-2, 1x MEM-NEAA, 1x GlutaMAX, and 1% penicillin-streptomycin) containing 25 ng/mL BMP4 and 8µM CHIR99021 supplements. On days 3 and 4, cultures received a media change with N2B27 media supplemented with 10 ng/mL Activin A and 10 ng/mL PDGF-BB. On day 5, pericytes were lifted with accutase, plated at 35,000 cells/cm^2^ onto plates coated in 0.1% gelatin, and grown in N2B27 containing 10 ng/mL PDGF-BB supplement. Cells received media changes with day 5 media every 2-3 days until confluent, at which point they were either frozen in freezing medium (90% KSR/10%DMSO) in liquid nitrogen tanks or split onto 0.1% coated gelatin plates in N2B27 media and maintained until ready for miBrain tissue assembly. For monoculture experiments, cells were plated onto 96-well µClear plastic-bottom plates (Greiner) coated with 0.1% gelatin at a density of 2500-5000 cells per well, and fixed 96 h post-plating unless specified otherwise.

#### Differentiation of human iPSCs into oligodendrocyte progenitor cells

At nearly 100% confluence, iPSCs in mTeSR1 with 10 µM Y27632 were plated on Matrigel (day-1). Every day until day 7, 2 mL of neural induction media (DMEM/F12 with 1x NEAA, GlutaMAX, 2-mercaptoethanol, and penicillin-streptomycin), with 10 μM SB431542, 250 nM LDN193189, and 100 nM RA (freshly added per use) was added. From day 8 through day 11, media was replaced daily with N2 medium (DMEM/F12 with 1x NEAA, GlutaMAX, 2-mercaptoethanol, and penicillin-streptomycin), along with 1x N2 and 100 nM RA and 1uM SAG (added fresh daily). On day 12, a cell scraper lifted the cells, which were gently transferred to a 6-well ultra-low attachment plate with 3 mL of OPC-N2B27 medium (DMEM/F12 with 1x NEAA, GlutaMAX, 2-mercaptoethanol, penicillin-streptomycin, 1x N2, 1x B27 without Vit A, 25 µg/=L insulin, and freshly added 100 nM RA and 1 µM SAG). About two-thirds of the OPC-NB27 media was changed every two days until day 20, without disturbance of the cell aggregates. From day 21 until day 30, PDGF medium (DMEM/F12 with 1x NEAA, GlutaMAX, 2-mercaptoethanol and penicillin-streptomycin, 1x N2, 1X B27 without Vit A, 25 ug/ml insulin, 100 ng/ml biotin, 1 µM cAMP, 5 ng/ml HGF, 10 ng/ml IGF-1, 10ng/ml NT3, and 10 ng/ml PDGFaa) was added every other day. On day 30, aggregates measuring 300-800 μm in size were picked with a p200 pipette and plated on PLO/laminin-coated 6-well adherent plates. Cells were plated at a density of 20 cells per well in PDGF medium. Until day 75, two-thirds of the PDGF media was changed every other day. PDGF media was then changed every 2-3 days, and the differentiated OPCs were enriched through FACS with a lineage-specific marker. OPC differentiation followed the protocol from Douvaras et al.^172^.

#### Differentiation of human iPSCs into microglia

To generate microglia, iPSCs were first differentiated into hematopoietic progenitor cells (HPCs) using STEMdiff™ Hematopoietic Kit (Catalog #05310). On days 12, 14, and 16 of the differentiation, floating HPCs were collected from media before freezing in CellBanker (AMSBIO). Following week 2 of miBrain assembly, HPCs were seeded into the existing tissue (25000 / 10 μL miBrain). miBrains were then maintained in miBrain media supplemented with 100ng/mL IL34 and 25ng/mL M-CSF. Microglia differentiation protocol was adapted from previous studies^173,174^.

### Assembly of 3D tissues

## 3D#Tissue Assembly for miBrains

Neurons, endothelial cells, mural cells, and OPCs were lifted with Accutase, and astrocytes in TrypLE Select. Cells were then resuspended in their respective media, counted, and resuspended at 1 × 10^6^ cells/ mL. To assemble miBrains, a 15 mL Falcon tube for each miBrain condition was prepared to receive approximately 5 × 10^6^ neurons, 5 × 10^6^ endothelial cells, 1 × 10^6^ astrocytes, 1 × 10^6^ OPCs, and 1 × 10^6^ pericytes per 1 mL. Each tube of pooled cells was spun down for five minutes at RT at 200 x g. After carefully aspirating as not to disturb the cell pellet, each resulting pellet was placed on ice before resuspension in 1 mL Geltrex^TM^ ,10 µM Y27632 and 5 µg/mL doxycycline. Resuspension technique took care to avoid air bubbles, and cell pellets were kept on ice to prevent premature Geltrex polymerization and ensure successful seeding. 10 µL of encapsulated cell suspensions were then plated per well of a 96-well µClear plastic-bottom plate (Greiner). To allow the Geltrex^TM^ to polymerize, seeded plates were transferred into a 37 °C 95%/5% Air/CO2 incubator for 30 minutes. After polymerization, miBrain cultures were completely submerged in miBrain week-1 medium (Human Endothelial Serum-free Medium, 1x Pen/Strep, 1X MEM-NEAA, 1X CD Lipids, 1x Astrocyte Growth Supplement (ScienCell), 1x B27 Supplement, 10ug/mL Insulin, 1µM cAMP-dibutyl, 50µg/mL Ascorbic acid, 10ng/mL NT3, 10ng/mL IGF, 100ng/mL Biotin, 60 ng/mL T3, 50 ng/mL VEGF, 1µM SAG, 5 µg/mL doxycycline). Each well received 200 μL of media. Every 2-3 days, a half-media change was performed until day 8, before changing the media to miBrain week-2 medium (Human Endothelial Serum-free Medium, 1x Pen/Strep, 1X MEM-NEAA, 1X CD Lipids, 1x Astrocyte Growth Supplement (ScienCell), 1x B27 Supplement, 10ug/mL Insulin, 1µM cAMP-dibutyl, 50µg/mL Ascorbic acid, 10ng/mL NT3, 10ng/mL IGF, 100ng/mL Biotin, 60 ng/mL T3, 5 µg/mL doxycycline). After 4 weeks, miBrain cultures were considered ready for downstream assays.

### 3D tissue assembly for iPSC-derived endothelial cell monocultures

This protocol is the same as the miBrain protocol with the following differences. Each 10 µL Geltrex droplet contained 120,000 of only endothelial cells. Cultures were maintained in hECSR media supplemented with 50 ng/mL VEGF and 5 µg/mL doxycycline for two weeks until fixation (see immunofluorescence methods section).

### Transduction of lentiviral shRNAs

FN1 and scramble MISSION lentiviral shRNAs were purchased from Sigma Aldrich. Lentivirus was generated in HEK293T cells following standard protocols. iPSC-derived mural cells were then transduced at a 1:40 dilution in mural cell culture media. Media was replaced 24 hours after transduction, and puromycin selection was initiated 72 hours post-transduction. Puromycin selection was maintained until cells were integrated into miBrains or collected for qRT-PCR validation of *FN1* knockdown.

### Amyloid and fibronectin treatments

3D iPSC-derived endothelial cell monocultures were generated as previously described. For the amyloid treatment group, 20 nM of both amyloid-β 1-40 and 1-42 were incubated in cell culture media for 1 hour at RT. For the amyloid and fibronectin treatment group, 20 nM of both amyloid-β 1-40 and 1-42 were incubated with 50 ng/mL fibronectin for 1 h in cell culture media at RT. Cells were treated with these mixtures for 96 h before fixing (see immunofluorescence methods section).

### TGF-*β* ligand treatments

iPSC-derived mural cells were seeded onto 96-well plates as previously described. 24 h post-plating, cells were treated with 50 ng/mL TGFβ1 for 96 hours followed by fixation (see immunofluorescence methods section).

### TGF-*β* inhibitor treatments

For 2D treatments, iPSC-derived mural cells were seeded onto 96-well plates as previously described. 24 hours post-plating, cells were treated with 10 µM SB431542 for 96 hours followed by fixation. For miBrains, treatments were either initiated at 3-4 days post-plating and maintained for 4 weeks until fixation or initiated at 2 weeks post-plating and maintained for the remaining 2 weeks until fixation, as specified in the figure legends. The concentrations used for miBrains were 10 µM SB431542 and 10 µM Galunisertib.

For TGF-β inhibition in APOE-TR mice, 12-month-old APOE4/4-TR mice were treated with the compound GW788388 for 4 weeks. Specifically, the mice were treated with 10 mg/kg of drug via daily intraperitoneal injections. The drug was prepared in a solution of 10% DMSO, 40% PEG300, 5% Tween-80, and 45% saline. This solution was also used as the DMSO vehicle for the control group. Following the treatment period, mice underwent cardiac perfusion as described in the ‘Immunofluorescence of mouse brain tissue’ section.

### Immunofluorescence of cell cultures

2D cultures were fixed in 4% paraformaldehyde for 15 minutes at room temperature, rinsed with PBS and permeabilized in 0.1% Triton-X100 in PBS for 10 minutes. Cultures were then blocked in 5% normal donkey serum in PBS for 2 h. Primary antibodies diluted 1:500 in blocking solution were added to cultures and incubated overnight at 4°C, before 3 consecutive rinses with PBS for 10, 20, and 30 minutes. Cultures were then incubated for 2h at room temperature with secondary antibodies diluted 1:1000 in blocking solution. After rinsing once with PBS for 10 min, cultures were incubated for 20 min in Hoechst 33258 (Sigma-Aldrich, Cat. no. 94403) diluted at 1:1000 PBS. Cultures were then rinsed once more with PBS for 30 min. The PBS was then replaced prior to imaging. See corresponding image acquisition and quantification section.

3D cultures and miBrains were incubated overnight at 4°C in blocking solution (0.3% Triton-X100, 5% normal donkey serum, in PBS). Primary antibodies, diluted 1:500 in blocking solution, were added the following day and left for 2-3 nights at 4°C. Cultures received 5 x 30 min rinses with 0.3% Triton-X100 PBS before adding secondary antibodies and Hoechst. Both were diluted at 1:1000 in blocking solution and cultures were left to incubate at 4°C for 1-3 nights. Cultures were rinsed for 5 x 30 min with 0.3% Triton-X100 PBS, before rinsing once more and leaving cells in regular PBS for image acquisition. See corresponding image acquisition and quantification section.

### Immunofluorescence of mouse brain tissue

Mice were first anaesthetized via gaseous isoflurane exposure, followed by cardiac perfusion with ice-cold PBS and subsequently 4% paraformaldehyde (PFA). The brains were dissected out and post-fixed in 4% PFA at 4C overnight. Fixed brains were placed in 30% sucrose in PBS until the tissue had sunk and were then embedded in OCT. Slices were cut at a thickness of 40 µm using a Leica cryostat and stored at -20C in cryoprotectant solution until staining.

For free-floating immunofluorescence, sections were carefully transferred into a 24-well plate with fresh PBS and rinsed two more times. Slices were then blocked in a buffer containing 0.3% Triton X-100 and 5% normal donkey serum in PBS overnight at 4° C. Primary antibodies diluted in blocking buffer were added the following day for 2-3 nights at 4° C. Sections were washed 5 x 30 min in PBS, and secondary antibodies diluted in blocking buffer were added for 2 hours at RT. Slices were then washed 5 x 30 min in PBS and mounted on glass slides for imaging. For slices stained with lectin, lectin was diluted 1:200 in PBS as part of the first wash and left for 1 h. Hoechst was added 1:1000 in PBS as part of the second wash and left for 30 min. All images were acquired in the corpus callosum or overlying cortex and in approximately the same locations of each slice to minimize biasing the results due to differences in anatomical regions (see image acquisition and quantification methods section).

## QUANTIFICATION AND STATISTICAL ANALYSIS

Images were acquired using a Nikon AX R confocal microscope. The same parameters were used for each image within an experiment, and the experimenter was blinded to the channels of interest or experimental condition during imaging. Where indicated in the figure legends, images underwent deconvolution using built-in microscope software as part of a standardized acquisition pipeline, with identical parameters applied across all images within each experiment.

Images were analyzed using scripts from the Nikon AX R built-in quantification software, CellProfiler^143^, or FIJI/ImageJ^152^. Prior to measurement, images underwent rolling-ball background subtraction. The background radius was chosen for each imaging experiment based on acquisition parameters and applied uniformly to all images within that experiment. No additional preprocessing was performed for quantitative analyses.

For visualization, some images were displayed after median filtering in FIJI as stated in the figure legends. The filter radius was selected based on imaging parameters and then applied identically to all images within each experiment. Median filtering was not used for quantitative measurements.

Statistical analysis was performed with GraphPad Prism or R and the tests used are specified in each figure caption.

## Extended Data Figure Legends

**Extended Data Figure 1.**
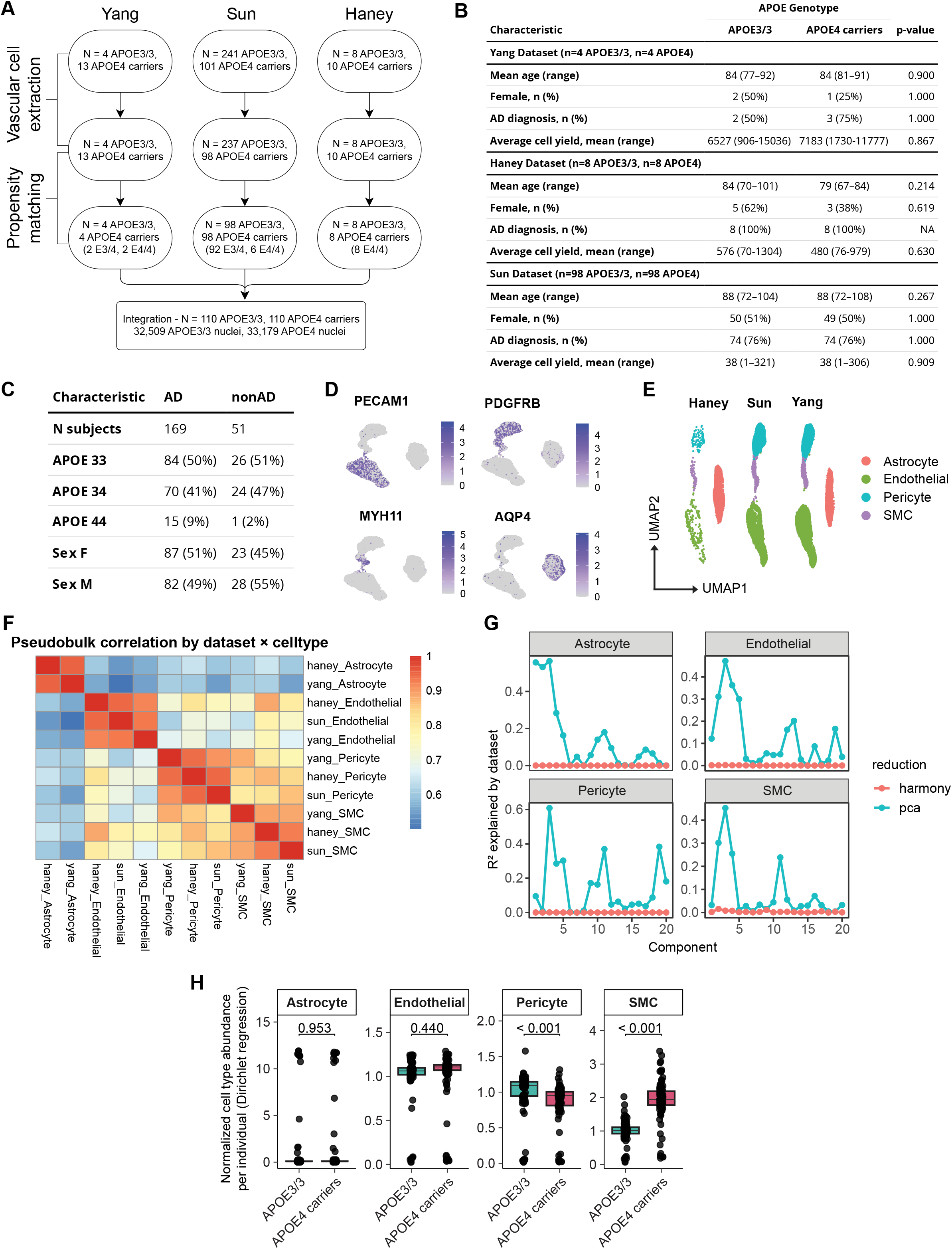
Metadata, quality control, and validation of single-nuclei analyses. **A.** Flowchart describing the generation of the final integrated cerebrovascular atlas used for downstream analyses. Astrocytes, endothelial cells, pericytes, and smooth muscle cells were extracted from each individual. Within each dataset, propensity matching based on age, sex, Alzheimer’s disease (AD) status, and vascular nuclei contribution was performed using the MatchIt package. **B.** Quality control of propensity matching. Unpaired two-sample Student’s t-tests were used to compare age and average vascular cell yield. Chi-squared tests were used to compare age and AD status for the Sun dataset. These two variables were compared using Fisher’s Exact Test in the Yang and Haney datasets. **C.** Subject characteristics stratified by AD and non-AD. **D.** Annotation of blood brain barrier cell types (*PECAM1* = endothelial cells, *PDGFRB* = pericytes, *MYH11* = smooth muscle cells, *AQP4* = astrocytes). **E.** UMAP of each dataset after integration. **F.** Pseudobulk gene-expression profiles were generated by averaging normalized expression values for each dataset × cell type combination, and pairwise Pearson correlations were computed across all pseudobulk profiles. **G.** The proportion of variance in each principal component explained by dataset identity (R² from linear models) was calculated within each major cell type before and after Harmony integration. **H.** Cell-type proportions were modeled using Dirichlet regression. Predicted proportions were normalized to APOE3/3 means for each cell type. Data points represent individual subjects (n = 110 per genotype). Boxes show median and IQR. P-values were derived from Tukey-adjusted pairwise comparisons between APOE3/3 and APOE4 carriers.

**Extended Data Figure 2.**
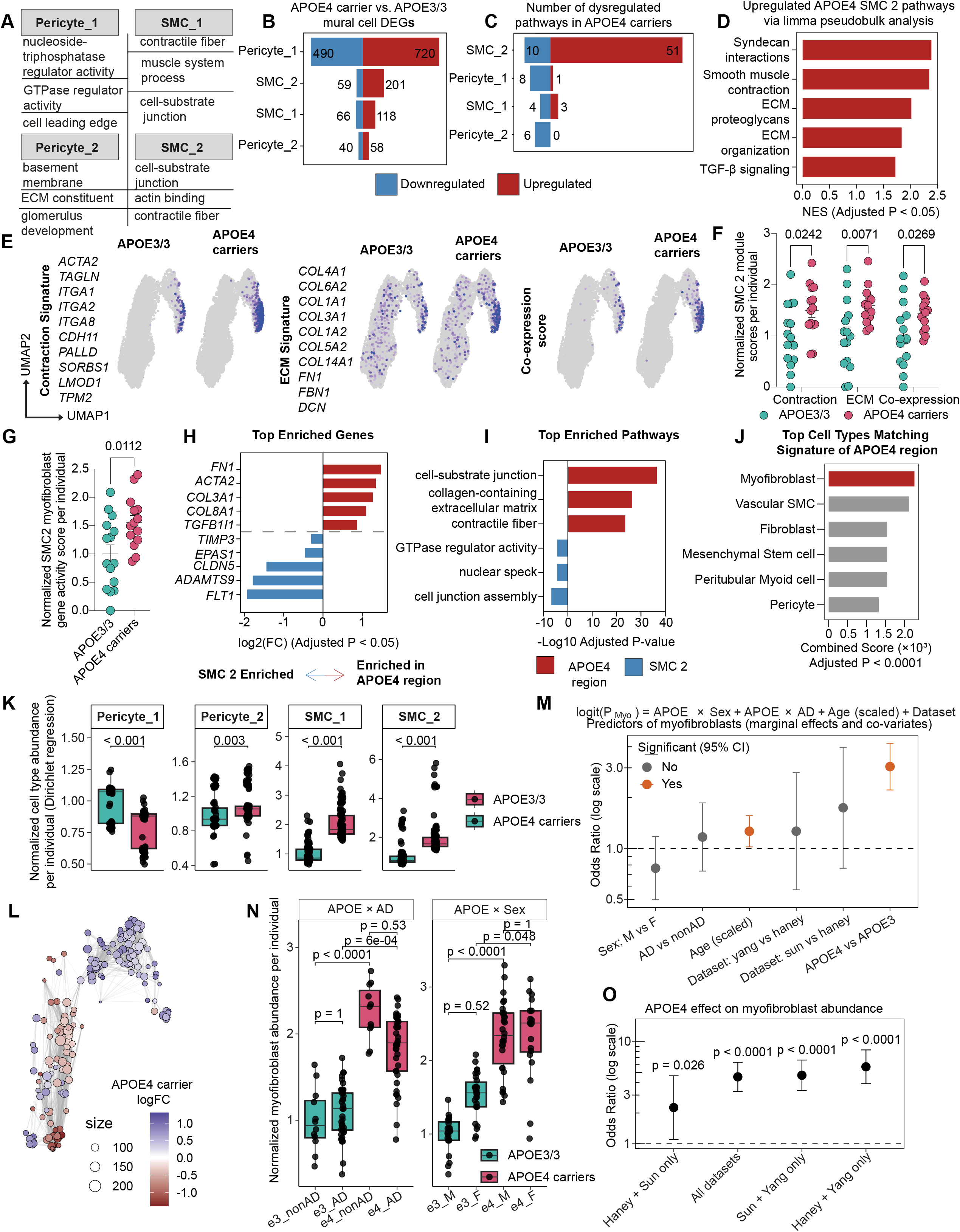
Clustering and analysis of human mural cells. **A.** Top GO pathways of each mural cell subcluster. Marker genes of each cluster were identified via Wilcoxon rank-sum test (logFC > 0.25, adjusted p < 0.05) and input into ClusterProfiler. **B.** Number of differentially expressed genes between APOE4 carriers and APOE3/3 mural cell subclusters. Analysis was performed using the MAST package, and differential expression was defined as adjusted p-value < 0.05 and |logFC| > 0.25. **C.** Number of dysregulated pathways per mural subcluster as described in Figure 1F. **D.** Validation of upregulated pathways in SMC_2 cells from APOE4 carriers using a pseudobulking approach (see Methods). After statistical analysis with the limma R package, genes were ranked according to sign(logFC) * -log10(P-value) and input into the fgsea package using the REACTOME database. Significantly upregulated pathways (NES > 0, FDR < 0.05) related to the extracellular matrix and contraction were then annotated, aligning with the results in Figure 1H. **E.** Feature map of mural cells annotated for expression of contraction or extracellular matrix (ECM) gene programs. Module scores were calculated using Seurat’s *AddModuleScore()* function based on the specified gene sets. A co-expression score was defined for each cell as the minimum of its contraction and ECM scores. **F.** Comparing SMC_2 contraction, ECM, and co-expression signature scores between APOE4 carriers and APOE3/3 individuals. Scores for SMC_2 cells calculated in E. were averaged per individual. Individuals contributing less than 5 cells were excluded from the analysis. Data points represent individuals (N = 15 APOE3/3, N = 14 APOE4 carriers). Bars are group means ± SEM. P-values were calculated using unpaired two-sample Student’s t-tests. **G.** Myofibroblast gene activity in SMC_2 cells by APOE genotype. SMC 2 gene expression was aggregated per individual. Individuals contributing less than 5 cells were excluded from the analysis. Gene set variation analysis (GSVA) was then performed using the GSVA R package. GSVA scores were assigned to each individual using previously described myofibroblast marker genes (see Methods). Data points represent individuals (N = 15 APOE3/3, N = 14 APOE4 carriers). Bars are group means ± SEM. P-values were calculated using unpaired two-sample Student’s t-tests. **H.** Representative marker genes of APOE4-enriched SMC_2 cells and non-enriched SMC_2 cells identified via DAseq in Figure 1H. The MAST package was utilized to identify differentially expressed genes (|logFC| > 0.25, adjusted p < 0.05) between these two subregions. **I.** The top three enriched GO pathways in APOE4-enriched cells compared to the remaining SMC_2 cells. The differentially expressed genes identified in H. were input into ClusterProfiler. **J.** Top cell type annotations of APOE4-enriched SMC_2 cells. The differentially expressed genes upregulated in APOE4 SMC_2 cells from H. were input into EnrichR with the CellMarker database. Redundant cell types were merged to enhance readability. **K**. Mural subcluster proportions were modeled using Dirichlet regression. Predicted proportions were normalized to APOE3/3 means for each cell type. Data points represent individual subjects (n = 106 per genotype). Boxes show median and IQR. P-values were derived from Tukey-adjusted pairwise comparisons between APOE3/3 and APOE4 carriers. **L.** UMAP of significantly differentially abundant (DA) cell neighborhoods in mural cells of APOE4 carriers vs. non-carriers using the miloR package (spatial FDR < 0.05, positive logFC indicates enrichment in APOE4 carriers). **M.** Binomial generalized linear model identifying predictors of myofibroblasts. The proportion of myofibroblasts relative to total SMC_2 cell count per individual (P_myofibroblast_) was modeled. **N.** Myofibroblast abundances stratified by AD diagnosis and sex are shown using model-predicted proportions from the binomial GLM described in M, normalized to APOE3/3 non-AD or APOE3/3 male. Data points represent individuals (n = 12 APOE3/3 non-AD, n = 41 APOE3/3 AD, n = 11 APOE4 non-AD, n = 39 APOE4 AD; n = 23 APOE3/3 males, n = 30 APOE3/3 females, n = 30 APOE4 males, n = 20 APOE4 females). P-values are from Tukey-adjusted emmeans pairwise comparisons. **O.** Leave-one-out dataset sensitivity analysis of the APOE4 effect on myofibroblast abundance.

**Extended Data Figure 3.**
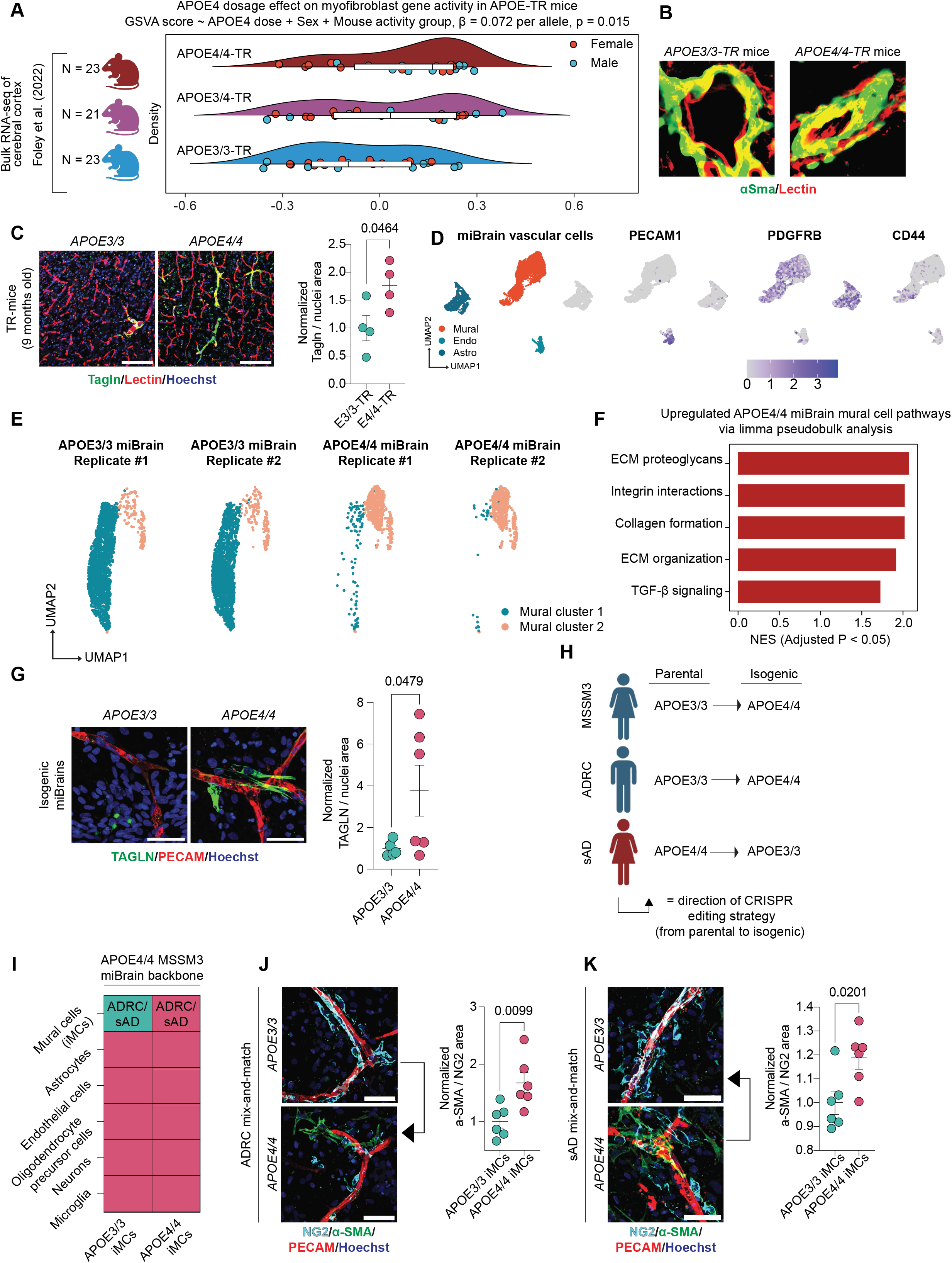
Additional validation of myofibroblast accumulation in APOE4 model systems. **A.** Correlation of APOE4 dosage with myofibroblast gene activity in APOE-TR mice. A processed bulk RNA sequencing dataset from the cerebral cortices of 4-month-old APOE3/3, APOE3/4, and APOE4/4-TR was downloaded from the source publication. Gene set variation analysis (GSVA) was performed using the GSVA R package. GSVA scores were assigned to each sample using the APOE4 myofibroblast marker genes identified in Extended Data Figure 2H. GSVA scores were modeled using linear regression including sex, activity group (from the original study), and APOE4 allele dosage (0 = APOE3/3, 1 = APOE3/4, 2 = APOE4/4). **B.** Representative images of large cerebral vessels stained for lectin and αSma in age- and sex-matched APOE3/3 and APOE4/4-TR mice. **C.** Representative images of cortical vessels of APOE3/3 and APOE4/4-TR mice stained for lectin and Tagln. Scale bar, 100 µm. For visualization only, a median filter was applied to the Tagln channel. The area positive for Tagln was quantified, normalized to nuclei, and expressed relative to APOE3/3. Data points represent mean values from individual mice (n = 6 images per mouse, n = 4 mice per genotype). Bars are mean group values ± SEM. P-values were calculated using unpaired two-sample Student’s t-tests. **D.** UMAP and annotation of miBrain vascular cell types using canonical marker genes. **E.** Per-sample UMAP of miBrain mural cells. **F.** Representative upregulated pathways of APOE4 miBrain mural cells using a pseudobulking approach (see Methods). After statistical analysis with the limma R package, genes were ranked according to sign(logFC) * -log10(P-value) and input into the fgsea package using the REACTOME database. Significantly upregulated pathways (NES > 0, FDR < 0.05) related to myofibroblasts were then annotated, aligning with the results in Figure 1O. **G.** Representative images of isogenic APOE3/3 and APOE4/4 miBrains stained for PECAM and TAGLN. Scale bar, 50 µm. The area positive for TAGLN was quantified, normalized to nuclei, and expressed relative to APOE3/3. Data points represent mean values from individual miBrains (n = 6 images per miBrain, n = 6 miBrains per genotype). Bars are mean group values ± SEM. P-values were calculated using an unpaired two-sample Student’s t-test. **H.** Characterization of iPSC lines utilized in this study. **I.** Schematic of mix-and-match miBrain experiments performed with additional isogenic pairs of iPSC-derived mural cells (iMCs). **J-K.** miBrains were immunostained for αSMA, NG2, and PECAM. Scale bar, 50 µm. The αSMA and NG2 area ratio was quantified and expressed relative to APOE3/3. Data points represent mean values from individual miBrains (n = 6 images per miBrain, n = 6 miBrains per genotype). Bars are mean group values ± SEM. P-values were calculated using unpaired two-sample Student’s t-tests.

**Extended Data Figure 4.**
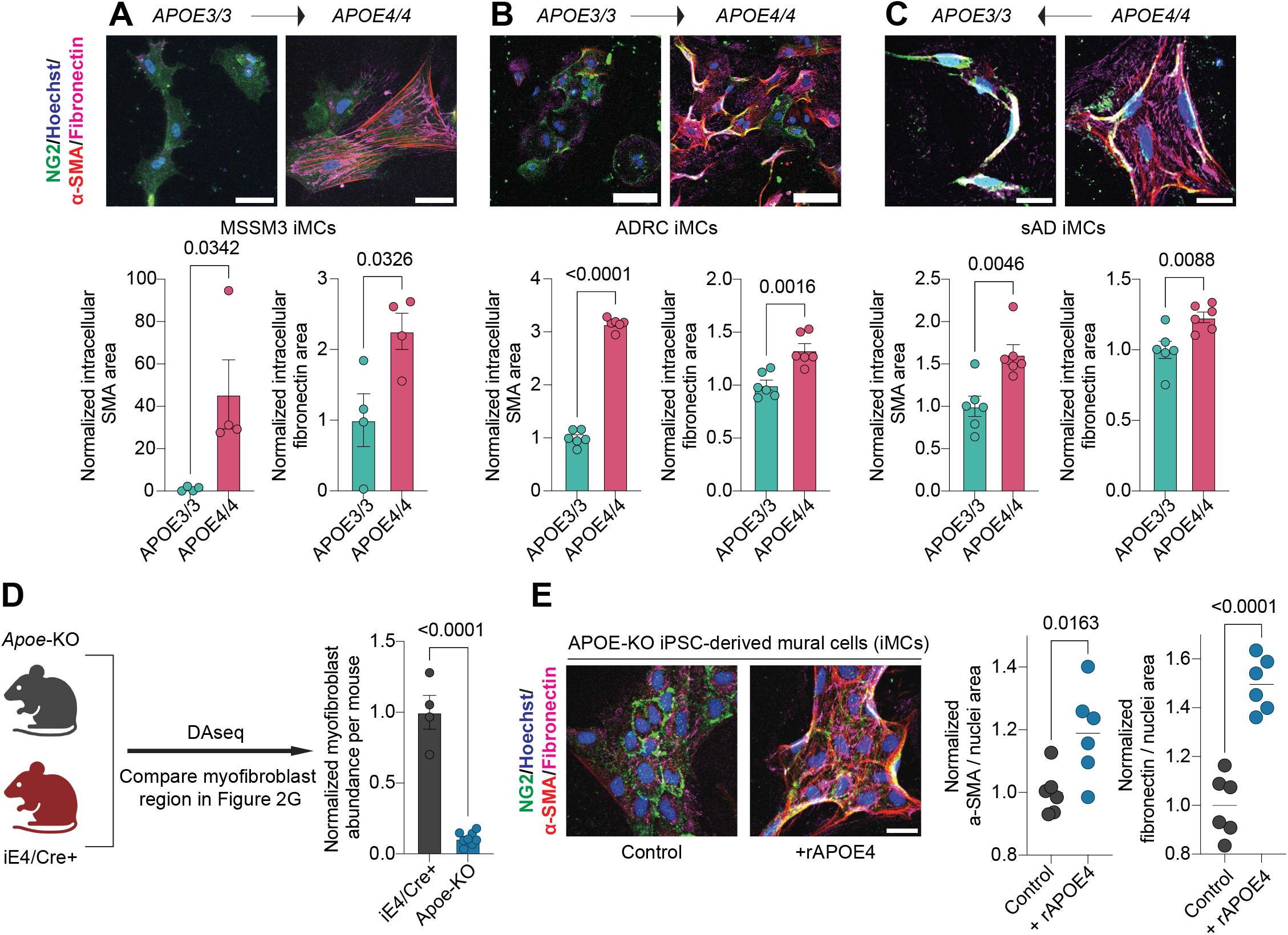
Supporting data for the sufficiency and necessity of APOE4 expression in mural cells for myofibroblast accumulation. A-C. Representative images of isogenic iMCs in monoculture stained for NG2, αSMA, and fibronectin. MSSM3 scale bar, 50 µm. ADRC scale bar, 100 µm. SAD scale bar, 25 µm. Intracellular a-SMA and fibronectin areas were quantified as the positively stained regions within the NG2 mask, normalized to the total NG2 area, and expressed relative to APOE3/3 levels. Data points represent mean values from individual wells (n = 6 images per well, n = 4-6 wells per genotype). Bars are mean group values ± SEM. P-values were calculated using unpaired two-sample Student’s t-tests. **D.** Observed myofibroblast abundance in a previously published mouse scRNA-seq dataset (Figure 2E) for iE4/Cre+ and *Apoe*-KO mice. Differential abundance analysis on the mural cell clusters between aged iE4/Cre+ (n = 4) versus Apoe-KO (n = 8) mice was performed with DAseq. The proportion of cells in the DAseq region corresponding to the myofibroblast region in Figure 2G were quantified per mouse and expressed relative to iE4/Cre+. The p-value shown corresponds to the fixed effect of genotype from a mixed-effects model that included sample as a random effect. **E.** Representative images of APOE-KO iPSC-derived mural cells (iMCs) treated with 200 ng/mL recombinant APOE4 protein. Scale bar, 25 µm. The area positive for a-SMA or fibronectin was quantified, normalized to nuclei, and expressed relative to control. Data points represent mean values from individual wells (n = 6 images per well, n = 6 wells per genotype). Bars are mean group values ± SEM. P-values were calculated using unpaired two-sample Student’s t-tests.

**Extended Data Figure 5.**
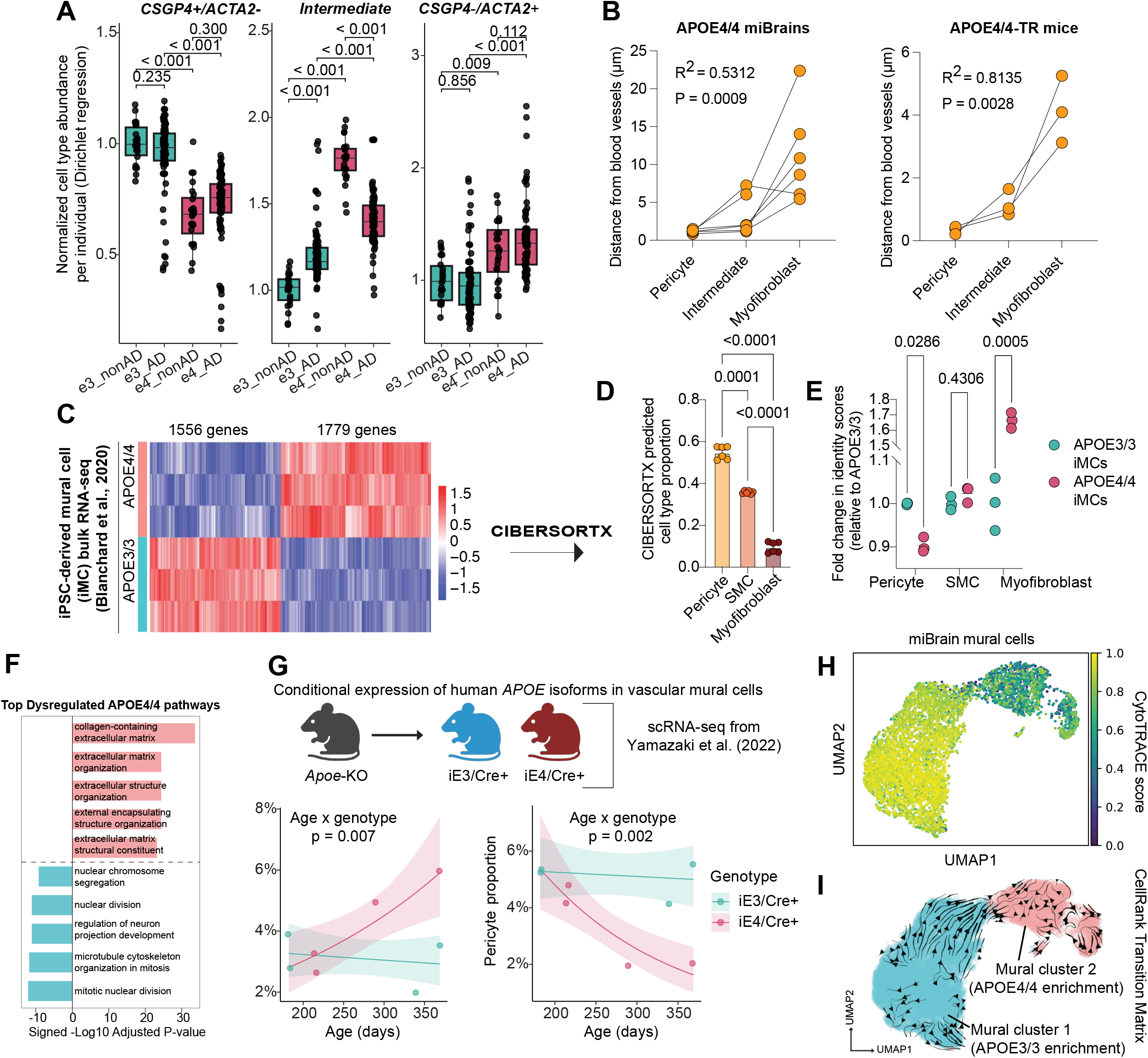
Supporting data for the APOE4 pericyte-to-myofibroblast transition. **A.** Subject-level predicted proportions of each mural cell state from the covariate-adjusted Dirichlet regression model stratified by AD diagnosis as described in the Methods. Data points represent individuals. Values were normalized to APOE3/3 non-AD. P-values are from Tukey-adjusted emmeans pairwise comparisons. **B.** Blood vessel distance measurements for pericytes, intermediate cells, and myofibroblasts in APOE4/4 miBrains and APOE4/4-TR mice. Cells were classified as described in Figure 2H. The *morph()* function in CellProfiler was used to calculate the distance of each pixel from the nearest PECAM+/lectin+ vascular region. For each classified cell, vascular distance was defined as the median pixel distance of all pixels within that cell. Distances were then averaged per miBrain or per mouse. Data points represent mean values from individual APOE4/4 miBrains or APOE4/4-TR mice (n = 6 miBrains, n = 3 mice). P-values were calculated using a repeated measures one-way ANOVA with a post-hoc test for a linear trend. **C.** CIBERSORTx deconvolution of bulk RNA-seq data from iPSC-derived mural cells (iMCs) using the mural cell subclusters from the single-nuclei cerebrovascular atlas as the reference (see Methods). **D.** Predicted relative abundance of mural cell subclusters in iMC bulk RNA-seq data via CIBERSORTx deconvolution. **E.** Relative abundance of mural cell subclusters stratified by APOE genotype. Bars are mean fold-change values ± SEM in APOE4/4 identity scores relative to APOE3/3. Data points represent individual bulk RNA-sequencing replicates (n = 3 per genotype). P-values were calculated using a two-way repeated measures ANOVA with Geisser-Greenhouse correction and Sidak’s multiple comparisons test. **F.** Top dysregulated GO pathways in APOE4/4 vs. APOE4/4 iMCs. Differentially expressed genes upregulated in APOE4/4 iMCs (|logFC| > 0.25, adjusted p < 0.05, calculated from limma package) were input into ClusterProfiler. **G.** Observed myofibroblast and pericyte proportions over time in a previously published mouse scRNA-seq dataset. Data points represent individual mice (n = 4 mice per genotype). Ribbons represent 95% confidence intervals. P values were calculated from the model described in the Methods. **H.** CytoTRACE score of miBrain mural cells. A higher score indicates more differentiation potential (a less differentiated state) whereas a lower score indicates less differentiation potential (a more differentiated state) (see Methods). **I.** CellRank transition matrix displaying predicted differentiation trajectories based on CytoTrace scores and k-nearest neighbor graph (see Methods).

**Extended Data Figure 6.**
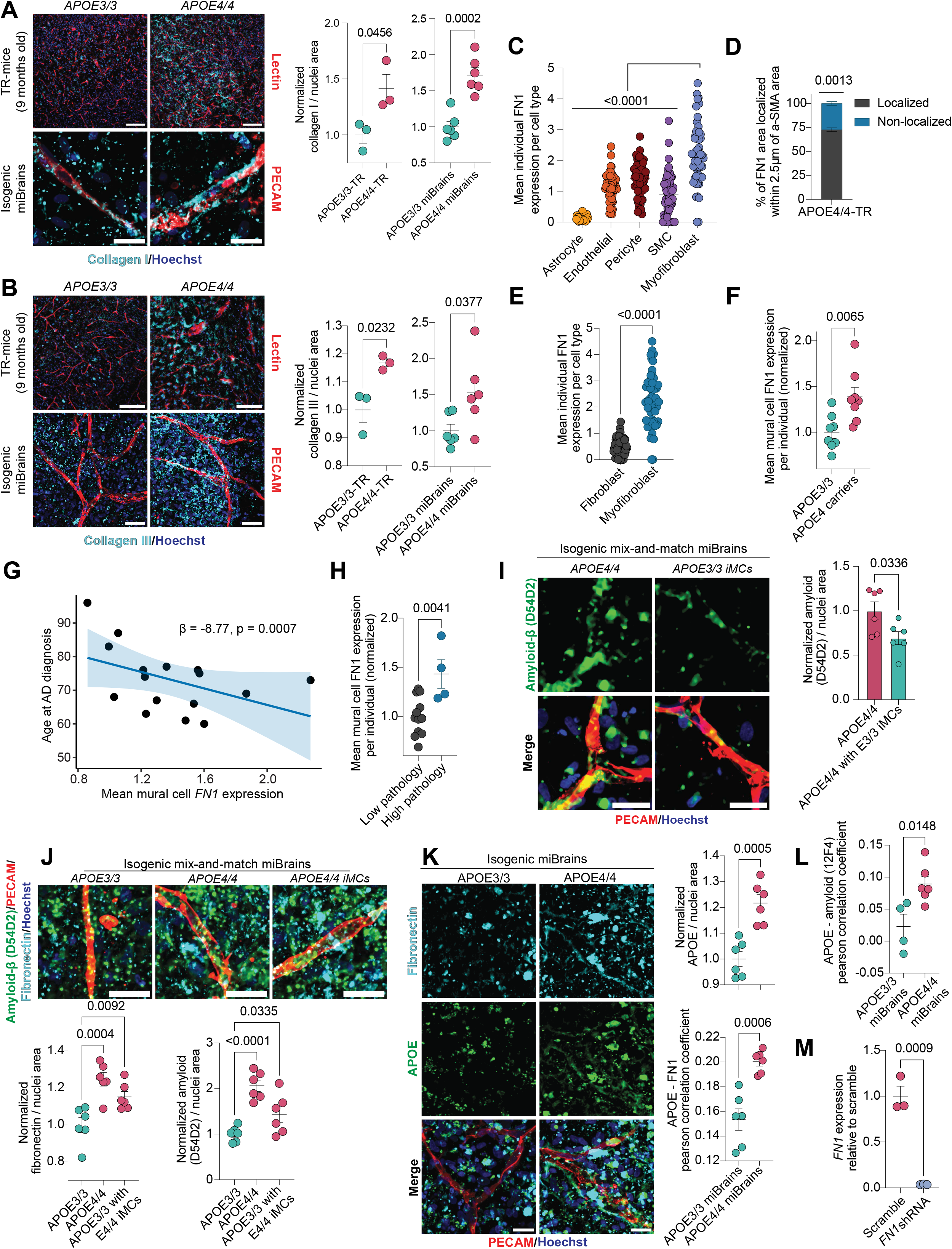
Additional markers and analysis of APOE4 cerebrovascular fibrosis and AD pathology. A-B. Representative images of cortical vessels from APOE3/3 and APOE4/4-TR mice and isogenic APOE3/3 and APOE4/4 miBrains stained for markers of blood vessels (lectin/PECAM) and collagen I or collagen III. Scale bar: APOE-TR, 100 µm; miBrain, 25 µm (A) or 50 µm (B). Mouse images were deconvolved using identical parameters. The area positive for collagen I or collagen III was quantified, normalized to nuclei, and expressed relative to APOE3/3. Data points represent mean values from individual mice or miBrains (n = 6 images per mouse or miBrain, n = 3 mice and 6 miBrains per genotype). Bars are mean group values ± SEM. P-values were calculated using unpaired two-sample Student’s t-tests. **C.** Quantification of FN1 expression across cell types in the cerebrovascular atlas. FN1 expression was aggregated per cell type for each individual. Individuals missing more than one cell type were excluded from the analysis. Data points represent individuals (n = 57). Bars are group means ± SEM. P-values were calculated using a repeated measures mixed-effects analysis with the Geisser-Greenhouse correction and Dunnett’s multiple comparisons test. **D.** Fibronectin deposition near αSMA staining in APOE4/4-TR mice. The percentage of fibronectin area within 2.5 µm of αSMA-positive area relative to the total fibronectin-positive area was quantified. Data points represent mean values from individual mice (n = 6 images per mouse, n = 4 mice). Bars are mean group values ± SEM. P-values were calculated using a one-sample t-test comparing the APOE4/4-TR mean to 50%. **E.** Comparison of *FN1* gene expression in fibroblasts versus myofibroblasts. Gene expression was aggregated per individual. See Methods. Data points represent individuals. Bars are group means ± SEM. P-values were calculated using a paired two-sample Student’s t-test. **F.** Mural cell *FN1* expression in APOE3/3 vs. APOE4 individuals (n = 8 per genotype). *FN1* expression was normalized to APOE3/3 Data points represent individuals. Bars are group means ± SEM. P-values were calculated using an unpaired two-sample Student’s t-test. **G.** The association between individual mural cell *FN1* expression and age at AD diagnosis (n = 16) was assessed via Theil–Sen robust linear regression. **H.** A global pathology score was calculated for each individual by averaging their amyloid plaque, neurofibrillary tangle, infarct, and Lewy body scores reported in the source metadata. These pathology scores were classified as low (bottom 75%) or high (top 25%). Mural cell *FN1* expression was then compared between the low (n = 12) and high (n = 4) pathology groups. *FN1* expression was normalized to the low pathology group. Data points represent individuals. Bars are group means ± SEM. P-values were calculated using an unpaired two-sample Student’s t-test. **I.** Representative images of APOE4/4 miBrains and APOE4/4 miBrains integrated with APOE3/3 iMCs stained for PECAM and an additional amyloid antibody (D54D2). Scale bar, 25 µm. For visualization only, a median filter was applied to the amyloid channel. The area positive for amyloid was normalized to nuclei and expressed relative to APOE4/4. Data points represent mean values from individual miBrains (n = 6 images per miBrain, n = 6 miBrains per genotype). Bars are group means ± SEM. P-values were calculated using an unpaired two-sample Student’s t-test. **J.** Representative images of isogenic APOE3/3, APOE4/4, and APOE3/3 miBrains integrated with APOE4/4 iMCs stained for PECAM, fibronectin, and amyloid (D54D2). Scale bar, 25 µm. For visualization only, a median filter was applied to the amyloid channel. The area positive for fibronectin and amyloid was quantified, normalized to nuclei, and expressed relative to APOE3/3. Data points represent mean values from individual miBrains (n = 6 images per miBrain, n = 6 miBrains per genotype). Bars are mean group values ± SEM. P-values were calculated using a one-way ANOVA with Holm-Sidak’s multiple comparisons test. **K-L.** Representative images of isogenic miBrains stained for APOE, fibronectin, and amyloid (12F4 antibody). Scale bars, 25 µm. The area positive for APOE was quantified, normalized to nuclei, and expressed relative to APOE3/3. Both APOE-fibronectin and APOE-amyloid colocalizations were quantified using Pearson’s correlation coefficient. Data points represent mean values from individual miBrains (n = 6 images per miBrain, n = 4-6 miBrains per genotype). Bars are mean group values ± SEM. P-values were calculated using unpaired two-sample Student’s t-tests. **M.** Validation of *FN1* shRNA knockdown. qRT-PCR was performed on APOE4/4 iMCs in monoculture. Data points represent mean values from individual wells of iMCs normalized to scramble (3 qRT-PCR replicates per well, n = 3 wells per condition). Bars are mean group values ± SEM. P-values were calculated using unpaired two-sample Student’s t-tests.

**Extended Data Figure 7.**
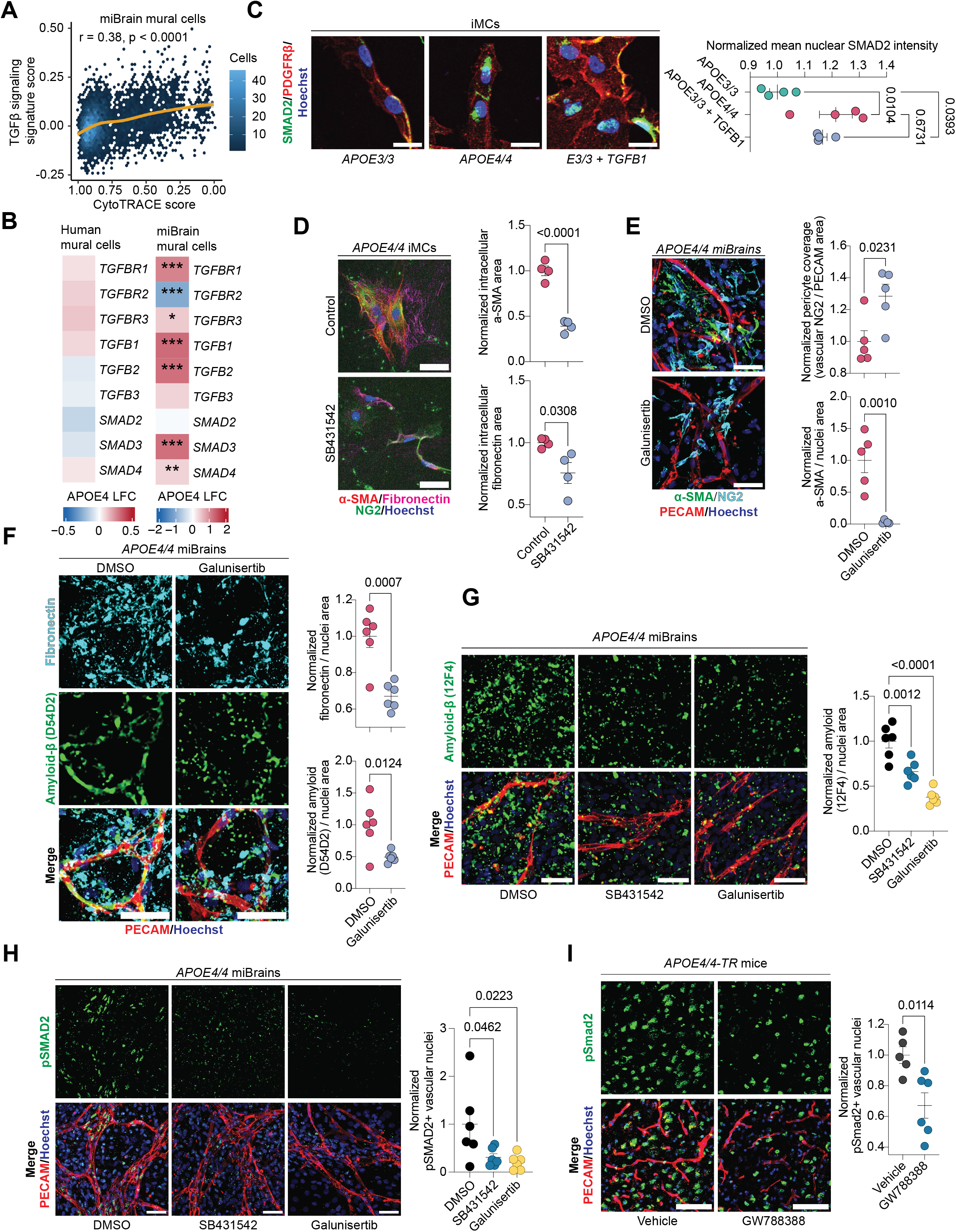
Supporting data of TGF-β signaling in the APOE4 pericyte-to-myofibroblast transition. **A.** TGF-β signaling activity in miBrain mural cells across cytoTRACE2 scores (see Methods). Correlation analysis was performed using Pearson’s correlation coefficient. **B.** Differential gene expression analysis of key TGF-β genes in human and miBrain mural cells. Gene log fold-changes of APOE4 mural cells vs. APOE3/3 and corresponding p-values were calculated via a pseudobulking approach (limma). **C.** Representative images of iMCs with and without TGFβ1 stimulation. Scale bar, 25 µm. For the control group, APOE3/3 and APOE4/4 iMCs were left untreated. For the experimental group, APOE3/3 iMCs were treated with 50 ng/mL TGFβ1 for 96 hours. iMCs were then fixed and stained for PDGFRβ and SMAD2. SMAD2 intensity inside the nuclear mask was quantified and expressed relative to APOE3/3 control. Data points represent mean values from individual wells of iMCs (6 images per well, 4 wells per condition). Bars are mean group values ± SEM. P-values were calculated using a one-way ANOVA with Dunnett’s test for multiple comparisons. **D.** Representative images of myofibroblast phenotypes in iMC monocultures after TGF-β inhibition. Scale bar, 50 µm. For the control group, APOE4/4 iMCs were left untreated. For the experimental group, APOE4/4 iMCs were treated with 10 µM SB431542 for 96 hours. iMCs were then fixed and stained for NG2, αSMA, and fibronectin. Intracellular αSMA and fibronectin areas were quantified as the positively stained regions within the NG2 mask, normalized to the total NG2 area, and expressed relative to APOE4/4 levels. Data points represent mean values from individual wells (n = 6 images per well, n = 4 wells per condition). Bars are mean group values ± SEM. P-values were calculated using unpaired two-sample Student’s t-tests. **E-F**. Representative images of APOE4/4 miBrains stained for PECAM, αSMA, NG2, fibronectin, and amyloid (D54D2) after TGF-β inhibition with Galunisertib. Scale bar, 50 µm. For visualization only, a median filter was applied to the amyloid channel. APOE4/4 miBrains were treated with either DMSO vehicle or 10 µM Galunisertib for four weeks. Three-dimensional pericyte coverage was measured by the overlapping NG2 and CD144 volume, normalized to the total CD144 volume, and expressed relative to APOE3/3. The area positive for αSMA, fibronectin, and amyloid was quantified, normalized to nuclei, and expressed relative to DMSO vehicle. Data points represent mean values from individual miBrains (n = 6 images per miBrain, n = 6 miBrains per condition). Bars are mean group values ± SEM. P-values were calculated using unpaired two-sample Student’s t-tests. **G.** Representative images of APOE4/4 miBrains stained for PECAM and an additional amyloid antibody (12F4) after TGF-β inhibition. Scale bar, 50 µm. For visualization only, a median filter was applied to the amyloid channel. APOE4/4 miBrains were treated with either DMSO vehicle, 10 µM SB431542, or 10 µM Galunisertib for four weeks. The area positive for αSMA, fibronectin, and amyloid was quantified, normalized to nuclei, and expressed relative to DMSO vehicle. Data points represent mean values from individual miBrains (n = 6 images per miBrain, n = 6 miBrains per condition). Bars are mean group values ± SEM. P-values were calculated using a one-way ANOVA with Dunnett’s test for multiple comparisons. **H.** Representative images of APOE4/4 miBrains stained for PECAM and pSMAD2 after TGF-β inhibition. Scale bar, 50 µm. APOE4/4 miBrains were treated with either DMSO vehicle, 10 µM SB431542, or 10 µM Galunisertib for four weeks. Nuclei along the vasculature were classified as pSMAD2+ using an intensity-based threshold in CellProfiler, normalized to the total number of vascular nuclei, and expressed relative to DMSO vehicle. Data points represent mean values from individual miBrains or mice (n = 6 images per miBrains, n = 6 miBrains per condition). Bars are mean group values ± SEM. P-values were calculated using a one-way ANOVA with Dunnett’s test for multiple comparisons. **I.** Representative images of aged APOE4/4-TR mice stained for lectin and pSMAD2 after TGF-β inhibition. Scale bar, 75 µm. Images were deconvolved using identical parameters. APOE4/4-TR mice were treated with either DMSO vehicle or 10 mg/kg of GW788388 daily for four weeks. Nuclei along the vasculature were classified as pSMAD2+ using an intensity-based threshold in CellProfiler, normalized to the total number of vascular nuclei, and expressed relative to DMSO vehicle. Data points represent mean values from individual mice (n = 6 images per mouse, n = 5 vehicle mice - one outlier removed via Grubbs’ test, n = 6 drug mice). Bars are mean group values ± SEM. P-values were calculated using an unpaired two-sample Student’s t-test.

## List of Supplementary Materials

a. Supplementary Information Guide
b. Supplementary Information
c. Source data

