## Supplementary Information Guide for "APOE4 promotes cerebrovascular fibrosis and amyloid deposition via a pericyte-to-myofibroblast transition"

**Supplementary Table S1. Cluster markers of the major cerebrovascular cell types, related to Figure 1.**

| **Column** | **Explanation** |
| --- | --- |
| gene | Gene name |
| p_val | Nominal p-value |
| avg_log2FC | Log2(fold change) in gene expression in the cluster of interest vs. the other clusters |
| pct.1 | Percent of cells in the cluster of interest expressing the gene |
| pct.2 | Percent of cells outside the cluster of interest expressing the gene |
| p_val_adj | Adjusted p-value |
| cluster | Cluster of interest |

**Supplementary Table S2. Significant differentially abundant cell neighborhoods in APOE4 carriers vs. non-carriers for the major cerebrovascular cell types via miloR analysis, related to Figure 1.**

| **Column** | **Explanation** |
| --- | --- |
| Nhood | The number assigned to that neighborhood in miloR |
| logFC | The log fold change in abundance for that neighborhood in APOE4 carriers vs. non-carriers (logFC > 0 indicates enrichment in APOE4 carriers) |
| logCPM | Average log normalized cell counts across all individuals |
| F | The f-statistic from the quali-likelihood F-test |
| PValue | Nominal p-value |
| FDR | P-value adjusted with Benjamini & Hochberg method |
| SpatialFDR | The FDR adjusted for spatial graph overlapds between neighborhoods |
| celltype | The cell type in which the neighborhood is assigned to |
| celltype_fraction | The proportion of cells in that neighborhood that belong to the assigned cell type |

**Supplementary Table S3. Cluster markers of the mural cell subtypes, related to Figure 1.**

| **Column** | **Explanation** |
| --- | --- |
| gene | Gene name |
| p_val | Nominal p-value |
| avg_log2FC | Log2(fold change) in gene expression in the cluster of interest vs. the other clusters |
| pct.1 | Percent of cells in the cluster of interest expressing the gene |
| pct.2 | Percent of cells outside the cluster of interest expressing the gene |
| p_val_adj | Adjusted p-value |
| cluster | Cluster of interest |

**Supplementary Table S4. ClusterProfiler pathway enrichment analysis of mural subcluster marker genes from S3, related to Figure 1.**

| **Column** | **Explanation** |
| --- | --- |
| ONTOLOGY | Ontology aspect in which that term belongs (cellular compartment [CC], molecular function [MF], or biological process [BP]) |
| ID | Ontology term ID |
| Description | Description of the ontology term |
| Gene ratio | Ratio of provided genes overlapping with the genes in the GO term |
| BgRatio | Ratio of all genes overlapping with the genes in the GO term |
| pvalue | Nominal p-value |
| p.adjust | Adjusted p-value |
| geneID | Overlapping genes between provided gene list and genes in the ontology term |
| cluster | Cluster in which the ontology term is enriched |

**Supplementary Table S5. Differential gene expression analysis in mural cell subclusters of APOE4 carriers vs. non-carriers via MAST, related to Figure 1.**

| **Column** | **Explanation** |
| --- | --- |
| gene | Gene name |
| p_val | Nominal p-value |
| avg_log2FC | Log2(fold change) in gene expression in APOE4 carriers vs. non-carriers for the given cluster (avg_log2FC > 0 indicates upregulation in APOE4 carriers) |
| pct.1 | Percent of APOE4 carrier cells in the given cluster expressing the gene |
| pct.2 | Percent of APOE3/3 cells in the given cluster expressing the gene |
| p_val_adj | Adjusted p-value |
| cluster | Cluster of interest |

**Supplementary Table S6. Gene set enrichment analysis of APOE4 carrier vs. non-carrier MAST results from S5, related to Figure 1.**

| **Column** | **Explanation** |
| --- | --- |
| pathway | Name of pathway |
| pval | Nominal p-value |
| padj | Adjusted p-value |
| ES | Enrichment score |
| NES | Normalized enrichment score in APOE4 carriers vs. non-carriers (NES > 0 indicates enrichment in APOE4 carriers) |
| size | Size of the pathway (number of genes in that pathway) |
| leadingEdge | Genes that are driving the enrichment |
| Cluster | Cluster of interest |

**Supplementary Table S7. Pseudobulked differential gene expression analysis in SMC_2 cluster of APOE4 carriers vs. non-carriers via limma, related to Figure 1.**

| **Column** | **Explanation** |
| --- | --- |
| gene | Gene name |
| logFC | Log2(fold change) in gene expression in APOE4 carriers vs. non-carriers (logFC > 0 indicates upregulation in APOE4 carriers) |
| AveExpr | Log2 expression level across all individuals |
| t | Moderated t-statistic |
| P.Value | Nominal p-value |
| adj.P.Val | Adjusted p-value |
| B | Log-odds that the gene is differentially expressed |

**Supplementary Table S8. Gene set enrichment analysis of APOE4 carrier vs. non-carrier SMC_2 limma results from S7, related to Figure 1.**

| **Column** | **Explanation** |
| --- | --- |
| pathway | Name of pathway |
| pval | Nominal p-value |
| padj | Adjusted p-value |
| ES | Enrichment score |
| NES | Normalized enrichment score in APOE4 carriers vs. non-carriers (NES > 0 indicates enrichment in APOE4 carriers) |
| size | Size of the pathway (number of genes in that pathway) |
| leadingEdge | Genes that are driving the enrichment |

**Supplementary Table S9. Cell type annotation of APOE4 SMC_2 gene signature via enrichR, related to Figure 1.**

| **Column** | **Explanation** |
| --- | --- |
| Term | Name of cell type |
| Overlap | Fraction of genes overlapping between the APOE4 gene list and the cell type |
| P.value | Nominal p-value |
| Adjusted.P.value | Adjusted p-value |
| Odds.Ratio | Association between APOE4 gene list and the cell type gene list |
| Combined.Score | Natural log of p-value multiplied by the z-score (deviation from expected rank) |
| Genes | Overlapping genes between the APOE4 gene list and the cell type gene list |

**Supplementary Table S10. Differential gene expression analysis in APOE4-enriched DAseq subregion vs. remaining SMC_2 cells via MAST, related to Figure 1.**

| **Column** | **Explanation** |
| --- | --- |
| gene | Gene name |
| p_val | Nominal p-value |
| avg_log2FC | Log2(fold change) in gene expression in APOE4-enriched DAseq subregion vs. remaining SMC_2 cells (avg_log2FC > 0 indicates upregulation in APOE4 region) |
| pct.1 | Percent of cells in APOE4 region expressing the gene |
| pct.2 | Percent of remaining SMC_2 cells expressing the gene |
| p_val_adj | Adjusted p-value |

**Supplementary Table S11. ClusterProfiler pathway enrichment analysis of differentially expressed genes in APOE4-enriched DAseq subregion vs. remaining SMC_2 cells from S10, related to Figure 1.**

| **Column** | **Explanation** |
| --- | --- |
| ONTOLOGY | Ontology aspect in which that term belongs (cellular compartment [CC], molecular function [MF], or biological process [BP]) |
| ID | Ontology term ID |
| Description | Description of the ontology term |
| Gene ratio | Ratio of provided genes overlapping with the genes in the GO term |
| BgRatio | Ratio of all genes overlapping with the genes in the GO term |
| pvalue | Nominal p-value |
| p.adjust | Adjusted p-value |
| geneID | Overlapping genes between provided gene list and genes in the ontology term |
| enriched_celltype | Region in which the ontology term is enriched (Myofibroblast = APOE4-enriched DAseq subregion [genes with avg_log2FC > 0 in S9]; SMC_2 = rest of SMC_2 cells [genes with avg_log2FC < 0 in S9)] |

**Supplementary Table S12. Cell type annotation of APOE4-enriched DAseq subregion via enrichR, related to Figure 1.**

| **Column** | **Explanation** |
| --- | --- |
| Term | Name of cell type |
| Overlap | Fraction of genes overlapping between the APOE4 gene list and the cell type |
| P.value | Nominal p-value |
| Adjusted.P.value | Adjusted p-value |
| Odds.Ratio | Association between APOE4 gene list and the cell type gene list |
| Combined.Score | Natural log of p-value multiplied by the z-score (deviation from expected rank) |
| Genes | Overlapping genes between the APOE4 gene list and the cell type gene list |

**Supplementary Table S13. Cluster markers of the miBrain mural cell subtypes, related to Figure 1.**

| **Column** | **Explanation** |
| --- | --- |
| gene | Gene name |
| p_val | Nominal p-value |
| avg_log2FC | Log2(fold change) in gene expression in mural cluster 2 vs. mural cluster 1 (positive logFC indicates enrichment in cluster 2) |
| pct.1 | Percent of cells in the cluster of interest expressing the gene |
| pct.2 | Percent of cells outside the cluster of interest expressing the gene |
| p_val_adj | Adjusted p-value |

**Supplementary Table S14. ClusterProfiler pathway enrichment analysis of miBrain mural cluster 2 marker genes from S13, related to Figure 1.**

| **Column** | **Explanation** |
| --- | --- |
| ONTOLOGY | Ontology aspect in which that term belongs (cellular compartment [CC], molecular function [MF], or biological process [BP]) |
| ID | Ontology term ID |
| Description | Description of the ontology term |
| Gene ratio | Ratio of provided genes overlapping with the genes in the GO term |
| BgRatio | Ratio of all genes overlapping with the genes in the GO term |
| pvalue | Nominal p-value |
| p.adjust | Adjusted p-value |
| geneID | Overlapping genes between provided gene list and genes in the ontology term |

**Supplementary Table S15. Pseudobulked differential gene expression analysis of APOE4/4 vs. APOE3/3 miBrain mural cells, related to Figure 1.**

| **Column** | **Explanation** |
| --- | --- |
| gene | Gene name |
| logFC | Log2(fold change) in gene expression in APOE4/4 vs. APOE3/3 (logFC > 0 indicates upregulation in APOE4/4 miBrains) |
| AveExpr | Log2 expression level across all individuals |
| t | Moderated t-statistic |
| P.Value | Nominal p-value |
| adj.P.Val | Adjusted p-value |
| B | Log-odds that the gene is differentially expressed |

**Supplementary Table S16. Gene set enrichment analysis of APOE4/4 vs. APOE3/3 miBrain mural cell limma results from S15, related to Figure 1.**

| **Column** | **Explanation** |
| --- | --- |
| pathway | Name of pathway |
| pval | Nominal p-value |
| padj | Adjusted p-value |
| ES | Enrichment score |
| NES | Normalized enrichment score in APOE4/4 miBrains (NES > 0 indicates enrichment in APOE4/4 miBrains) |
| size | Size of the pathway (number of genes in that pathway) |
| leadingEdge | Genes that are driving the enrichment |

**Supplementary Table S17. Cluster markers of the aged iE4/Cre+ enriched SMC region, related to Figure 2.**

| **Column** | **Explanation** |
| --- | --- |
| gene | Gene name |
| p_val | Nominal p-value |
| avg_log2FC | Log2(fold change) in iE4/Cre+ enriched region (positive logFC indicates enrichment in that region compared to the other cells) |
| pct.1 | Percent of cells in the cluster of interest expressing the gene |
| pct.2 | Percent of cells outside the cluster of interest expressing the gene |
| p_val_adj | Adjusted p-value |

**Supplementary Table S18. ClusterProfiler pathway enrichment analysis of aged iE4/Cre+ enriched region from S17, related to Figure 2.**

| **Column** | **Explanation** |
| --- | --- |
| ONTOLOGY | Ontology aspect in which that term belongs (cellular compartment [CC], molecular function [MF], or biological process [BP]) |
| ID | Ontology term ID |
| Description | Description of the ontology term |
| Gene ratio | Ratio of provided genes overlapping with the genes in the GO term |
| BgRatio | Ratio of all genes overlapping with the genes in the GO term |
| pvalue | Nominal p-value |
| p.adjust | Adjusted p-value |
| geneID | Overlapping genes between provided gene list and genes in the ontology term |

**Supplementary Table S19. Cell type annotation of aged iE4/Cre+ enriched region via enrichR, related to Figure 2.**

| **Column** | **Explanation** |
| --- | --- |
| Term | Name of cell type |
| Overlap | Fraction of genes overlapping between the APOE4 gene list and the cell type |
| P.value | Nominal p-value |
| Adjusted.P.value | Adjusted p-value |
| Odds.Ratio | Association between APOE4 gene list and the cell type gene list |
| Combined.Score | Natural log of p-value multiplied by the z-score (deviation from expected rank) |
| Genes | Overlapping genes between the APOE4 gene list and the cell type gene list |

**Supplementary Table S20. Differential gene expression analysis of APOE4/4 iPSC-derived mural cells (iMCs) vs. APOE3/3 iMCs via limma, related to Figure 3.**

| **Column** | **Explanation** |
| --- | --- |
| gene | Gene name |
| logFC | Log2(fold change) in gene expression in APOE4/4 iMCs vs. APOE3/3 iMCs (logFC > 0 indicates upregulation in APOE4/4 iMCs) |
| AveExpr | Log2 expression level across all samples |
| t | Moderated t-statistic |
| P.Value | Nominal p-value |
| adj.P.Val | Adjusted p-value |
| B | Log-odds that the gene is differentially expressed |

**Supplementary Table S21. ClusterProfiler pathway enrichment analysis of differentially expressed genes in APOE4/4 iMCs vs. APOE3/3 iMCs from S20, related to Figure 3.**

| **Column** | **Explanation** |
| --- | --- |
| ONTOLOGY | Ontology aspect in which that term belongs (cellular compartment [CC], molecular function [MF], or biological process [BP]) |
| ID | Ontology term ID |
| Description | Description of the ontology term |
| Gene ratio | Ratio of provided genes overlapping with the genes in the GO term |
| BgRatio | Ratio of all genes overlapping with the genes in the GO term |
| pvalue | Nominal p-value |
| p.adjust | Adjusted p-value |
| geneID | Overlapping genes between provided gene list and genes in the ontology term |
| direction | Whether the ontology term is upregulated or downregulated in APOE4/4 iMCs (Upregulated = genes with logFC > 0 in S12; Downregulated = genes with logFC < 0 in S12) |

**Supplementary Table S22. Differential gene expression analysis in fibroblasts of APOE4 carriers vs. non-carriers via MAST, related to Figure 4.**

| **Column** | **Explanation** |
| --- | --- |
| gene | Gene name |
| p_val | Nominal p-value |
| avg_log2FC | Log2(fold change) in gene expression in APOE4 carriers vs. non-carriers (avg_log2FC > 0 indicates upregulation in APOE4 carriers) |
| pct.1 | Percent of APOE4 carrier fibroblast cells expressing the gene |
| pct.2 | Percent of APOE3/3 fibroblast cells expressing the gene |
| p_val_adj | Adjusted p-value |

**Supplementary Table S23. Gene set enrichment analysis of APOE4 carrier vs. non-carrier fibroblast MAST results from S22, related to Figure 4.**

| **Column** | **Explanation** |
| --- | --- |
| pathway | Name of pathway |
| pval | Nominal p-value |
| padj | Adjusted p-value |
| ES | Enrichment score |
| NES | Normalized enrichment score in APOE4 carriers vs. non-carriers (NES > 0 indicates enrichment in APOE4 carriers) |
| size | Size of the pathway (number of genes in that pathway) |
| leadingEdge | Genes that are driving the enrichment |

**Supplementary Table S24. NicheNet ligand activities in APOE4 myofibroblasts, related to Figure 5.**

| **Column** | **Explanation** |
| --- | --- |
| test_ligand | Ligand name |
| aupr | Area under the precision-recall curve (AUPR) |
| aupr_corrected | Corrected AUPR value |
